# Benzofuran sulfonates and small self-lipid antigens activate type II NKT cells via CD1d

**DOI:** 10.1101/2021.03.05.433980

**Authors:** CF Almeida, D Smith, T-Y Cheng, C Harpur, E Batleska, T Nguyen, T Hilmenyuk, CV Nguyen-Robertson, SJJ Reddiex, I Van Rhijn, J Rossjohn, AP Uldrich, DB Moody, SJ Williams, DG Pellicci, DI Godfrey

## Abstract

Natural Killer T (NKT) cells detect lipids presented by CD1d. Most studies focus on type I NKT cells that express semi-invariant αβ T cell receptors (TCR) and recognise α-galactosylceramides. However, CD1d also presents structurally distinct lipids to NKT cells expressing diverse TCRs (type II NKT cells) but our knowledge of the antigens for type II NKT cells is limited. An early study identified an NKT cell agonist, phenyl pentamethyldihydrobenzofuransulfonate (PPBF), which is notable for its similarity to common sulfa-drugs, but its mechanism of NKT-cell activation remained unknown. Here we demonstrate that a range of pentamethylbenzofuransulfonate (PBFs), including PPBF, activate polyclonal type II NKT cells from human donors. Whereas these sulfa drug-like molecules might have acted pharmacologically on cells, here we demonstrate direct contact between TCRs and PBF-treated CD1d complexes. Further, PBF-treated CD1d-tetramers identified type II NKT cell populations cells expressing αβ and γδTCRs, including those with variable and joining region gene usage (TRAV12-1–TRAJ6) that was conserved across donors. By trapping a CD1d-type II NKT TCR complex for direct mass spectrometric analysis, we detected molecules that allow binding of CD1d to TCRs, finding that both PBF and short-chain sphingomyelin lipids are present in these complexes. Furthermore, the combination of PPBF and short-chain sphingomyelin enhances CD1d tetramer staining of PPBF-reactive T cell lines over either molecule alone. This study demonstrates that non-lipidic small molecules, that resemble sulfa-drugs implicated in systemic hypersensitivity and drug allergy reactions, activate a polyclonal population of type II NKT cells in a CD1d-restricted manner.

**Significance Statement:** Whereas T cells are known to recognize peptide, vitamin B metabolite or lipid antigens, we identify several non-lipidic small molecules, pentamethylbenzofuransulfonates (PBFs), that activate a population of CD1d-restricted NKT cells. This represents a breakthrough in the field of NKT cell biology. This study also reveals a previously unknown population of PBF-reactive NKT cells in healthy individuals with stereotyped receptors that paves the way for future studies of the role of these cells in immunity, including sulfa-drug hypersensitivity.

## Introduction

Natural Killer T (NKT) cells are defined as T cells that are restricted to the lipid-antigen presenting molecule, CD1d. The most extensively studied are type I NKT cells, which typically express an invariant T cell receptor (TCR)-α chain consisting of TRAV10–TRAJ18 in humans (TRAV11– TRAJ18 in mice) paired with a constrained repertoire of TCR-β chains, enriched for TRBV25 in humans (TRBV13, 29, 1 in mice) (reviewed in^1^). Type I NKT cells are defined by their strong responses to α-galactosylceramide (α-GalCer) and structurally-related hexosyl ceramides presented by CD1d, while in contrast, type II NKT cells are defined as CD1d-restricted T cells that express diverse TCRs and do not recognize α-GalCer (reviewed in^1, 2^). Very little is known about the chemical identity of antigens for type II NKT cells; however, some studies suggest that these cells are abundant in humans^3, 4, 5^ and, by virtue of their greater TCR diversity, they can interact with a broader range of antigens compared to type I NKT cells^2, 6, 7, 8, 9, 10, 11, 12, 13, 14^.

In 2004, a non-lipidic molecule, phenyl pentamethyldihydrobenzofuransulfonate (PPBF), was described that stimulated a human TRAV10^-^ T (type II) NKT cell line in a CD1d-dependent manner^15^. These observations were notable because PPBF resembles various sulfonamide drugs: furosemide (diuretic), sulfasalazine (disease-modifying anti-rheumatic drug) and celecoxib (anti-inflammatory), as well as ‘sulfa’ antibiotics such as sulfonamide, sulfapyridine, sulfamethoxazole, sulfadiazine, sulfadoxine^16^. These drugs cause systemic delayed-type hypersensitivity reactions, which are thought to be mediated by T cells^17, 18, 19, 20^. Most of our limited understanding of drug hypersensitivity reactions comes from work with HLA-restricted conventional T cells, which has led to the proposal of four main mechanisms for small-drug immune-activity, as reviewed in^21^: 1) hapten/prohapten formation, whereby the drug reacts with self-Ags to generate a neo-product that undergoes processing and presentation to T cells; 2) non-covalent/labile pharmacological interaction (p.i. concept) with immune receptors on the cell surface; 3) superantigen mediating direct linkage of TCRs and Ag-presenting molecules; 4) anchor-site occupation by small-molecules in Ag-presenting molecules inducing an altered-self-Ag repertoire^22^. Whether the CD1d-NKT cell axis can be implicated in drug hypersensitivity remains unclear. Whereas most antigens in the CD1d system are lipids that use their linear alkyl chains to bind to CD1d, PPBF is a polycyclic small molecule and so might act through an atypical display mechanism. As previous attempts to use CD1d-PPBF tetramers to stain the T cell line were unsuccessful, the atypical drug-like structure of PPBF raised the possibility of direct pharmacological action on T cells, rather than presentation of CD1d-PPBF complexes to TCRs. However, the mechanism of PPBF-mediated type II NKT cell activation remains undefined^15^.

Here, we used TCR-transduced cell lines, CD1d tetramers treated with PPBF, and new analogues in the pentamethylbenzofuransulfonate (PBF) family to discover that several molecules stimulate polyclonal NKT cells. Using CD1d tetramers treated with a newly identified and more potent analogue of PPBF, we identified populations of type II NKT cells that comprise a polyclonal repertoire of both αβ and γδ T cells, including those with conserved TCR sequences. This enigmatic nature of T cell responses to small molecules was resolved using TCR-trap technology^23, 24^. Mass spectrometric analysis of all molecules present in CD1d-PBF-TCR complexes indicates that CD1d binds PBFs and small self-lipids that promote CD1d-TCR binding. These data support a model of type II NKT cell recognition of small sulfa-drug like compounds in association with CD1d and flag a possible mechanism where such cells may be involved in sulfa-drug hypersensitivity.

## Results

### Role of the TCR in recognition of PPBF molecules

Previously, it was demonstrated that PPBF can stimulate a TRAV10^-^ αβ T cell clone, ABd, which was derived from polyclonal T cells expanded under PPBF stimulation of human PBMCs^15^. While activation of ABd required CD1d^15^, attempts to stain the line with PPBF-treated tetramers were unsuccessful. PPBF is a structurally unconventional stimulant of CD1d-reactive T cells, and these observations are consistent with either low affinity binding of the TCR to PPBF-treated CD1d complexes, or that PPBF acts by an atypical mechanism such as direct pharmacological activation of cells via non-TCR mechanisms. We used the ABd TCR sequence (TRAV12-1–TRAJ6, TRBV11) to generate a T cell line by transducing β2m-deficient SKW3 cells (SKW3.β2m^-/-^, lacking endogenous CD1d) to conduct assays to directly examine the role of the ABd TCR in PPBF responsiveness.

Given the unusual, non-lipidic structure of PPBF, we studied the structural requirements for T cell recognition by using analogues of PPBF. Given that our previous work demonstrated that 3-methyl-PPBF was more potent than the 2- and 4-methyl isomers^15^, we focused on substitutions at the phenyl 3-position. Using a one-step direct method for synthesis of these molecules, we altered the methyl group to substituents varying in electron density (F, Cl, Br), steric bulk (isopropyl, iPr; *tert*-butyl, tBu; phenyl, Ph), and electron-donating/withdrawing ability (Me_2_N, MeO, NO_2_, CF_3_PPBF). Additional analogues included 2,3-dimethyl and 3,5-dichloro derivatives (DMePPBF and DClPPBF, respectively) to examine whether substituent effects are additive. We increased the ring size of the benzofuran to a benzochroman and studied linker variation by replacing the sulfonate with a sulfonamide (APBF, mAPBF and MAPBF). We also included the unsubstituted cores: PBF, PBC and a water-soluble analogue, Trolox (**Figure 1A**). These compounds were assessed for their ability to activate the ABd-TCR transduced cell line, or a control type I human NKT TCR-expressing line (NKT15)^25^ in co-cultures with C1R Ag-presenting cells. CD69 upregulation was measured as a readout for cellular activation, and strong dose-dependent responses were detected to many of these analogs by the ABd cell line (**Figure 1B**), but not in the control NKT15 cell line, which responded only to α-GalCer.

**Figure 1.**
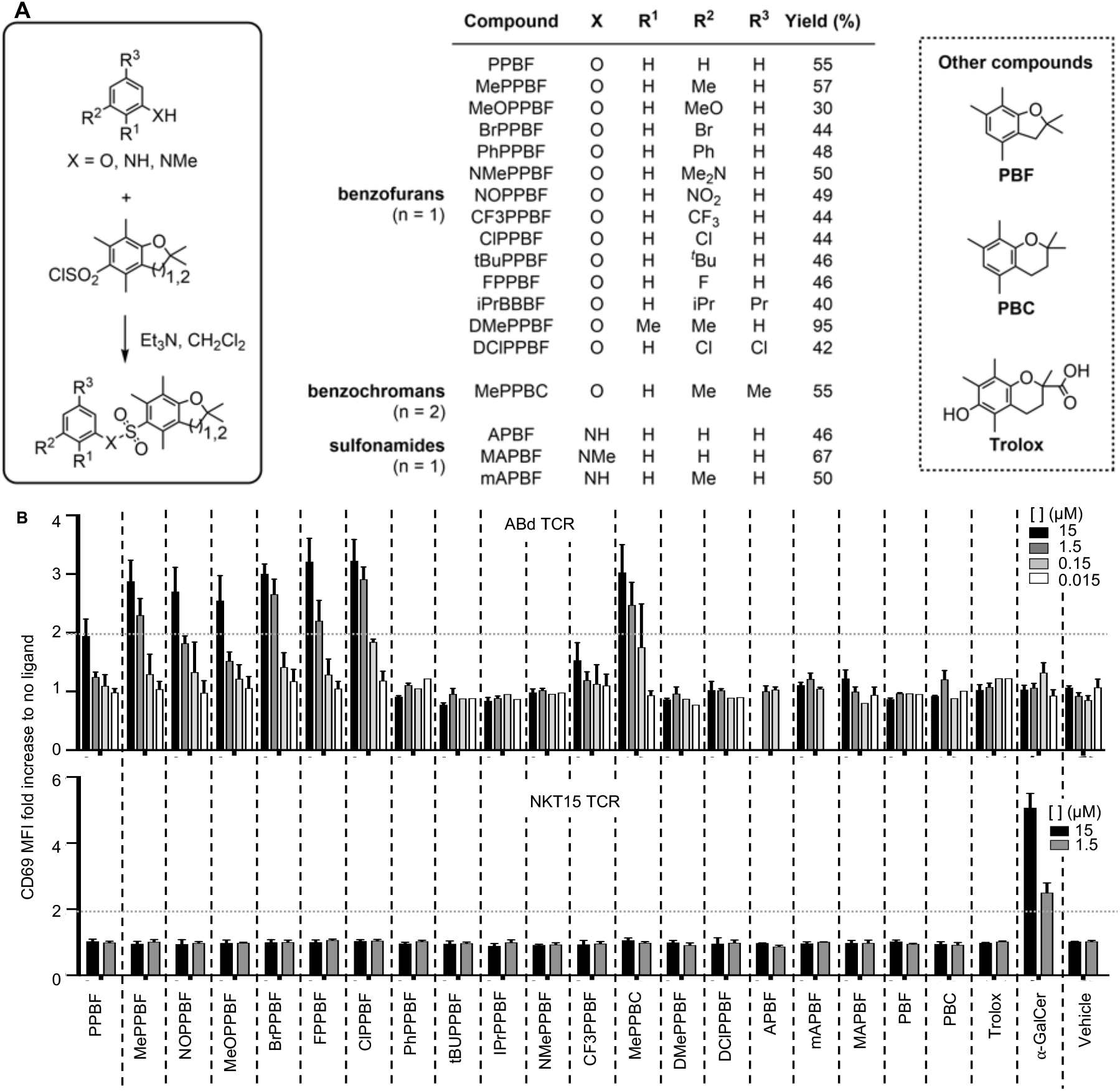
Structure activity relationship study of PPBF analogs on type II NKT cells. **A.** Synthesis of various PPBF analogs: including modifications at positions 2’ and 3’ of the phenyl ring, the sulfonate linkage and ring expansion of the benzofuran. Values depict reaction yields. **B.** These were serially diluted (10-fold dilutions ranging 15μM to 0.015μM) and incubated with C1R antigen presenting cells (APC) and assessed for their ability to activate a SKW3.β2m^-/-^ cell line stably transduced with the ABd type II NKT TCR (PPBF-reactive^15^). α-GalCer and the NKT15 type I NKT TCR^+^ cell line (α-GalCer reactive^25^). Graphs depict fold increase of CD69 upregulation in comparison to conditions with no ligand after 16h activation. Dotted horizontal line marks a 2-fold increase over no ligand control. Data is from 3 independent experiments + SEM.

Both compounds from our initial study^15^, PPBF and 3-methyl PPBF (MePPBF), activated ABd-TCR^+^ cells, eliciting a 2- and 3-fold increase in CD69 mean fluorescence intensity (MFI), respectively, when used at 15 μM (**Figure 1B**). This effect was largely lost when PPBF was used at 1.5 μM, but MePPBF was still active at this lower concentration, consistent with previous work that showed this analog was more potent than PPBF^15^. This finding is notable because ABd was derived in the presence of PPBF, yet the resulting T cell line responds better to MePPBF. Stronger responses than the unsubstituted parental PPBF were also elicited by NOPPBF, MeOPPBF, FPPBF and, most notably, by BrPPBF and ClPPBF, which retained activity even at 0.15 μM. Conversely, PhPPBF, tBuPPBF, iPrPPBF, NMePPBF and CF_3_PPBF failed to induce responses, indicating a preference for a small group at the 3-position. The benzochroman MePPBC (and the 3-chloro analogue thereof (**Supplemental Figure 1**)), was as potent as ClPPBF, whereas isomerization of the sulfonate linkage to give the 3-methyl ‘isosulfonate’ led to loss of activity and binding (**Supplemental Figure 1**), indicating the importance of linker orientation. The disubstituted compounds (DMePPBF and DClPPBF) did not elicit a response, highlighting the preference for substitution only at the 3-position. Finally, the sulfonamides APBF, mAPBF, MAPBF and the unsubstituted core analogues PBF, PBC (a benzochroman), and Trolox, did not induce a response, indicating that a sulfonate linker and the phenyl group are important for T cell activation (**Figure 1B**).

Next, we challenged the ABd TCR-transduced cell line with PPBF, mPPBF and α-GalCer in the presence of blocking anti-CD1d or isotype control antibodies (**Figure 2A**). PPBF and mPPBF stimulated the ABd TCR-transduced cells in a CD1d-dependent manner and were blocked by anti-CD1d antibody (αCD1d) but not by an isotype control. PBF compounds did not activate the NKT15 type I NKT cell line, suggesting no general mitogenic effect. Conversely, NKT15 TCR-transduced cells were highly responsive to α-GalCer, and blocked by anti-CD1d, but were unresponsive to PPBF or mPPBF, demonstrating that the response to these compounds is TCR- and CD1d-dependent.

**Figure 2.**
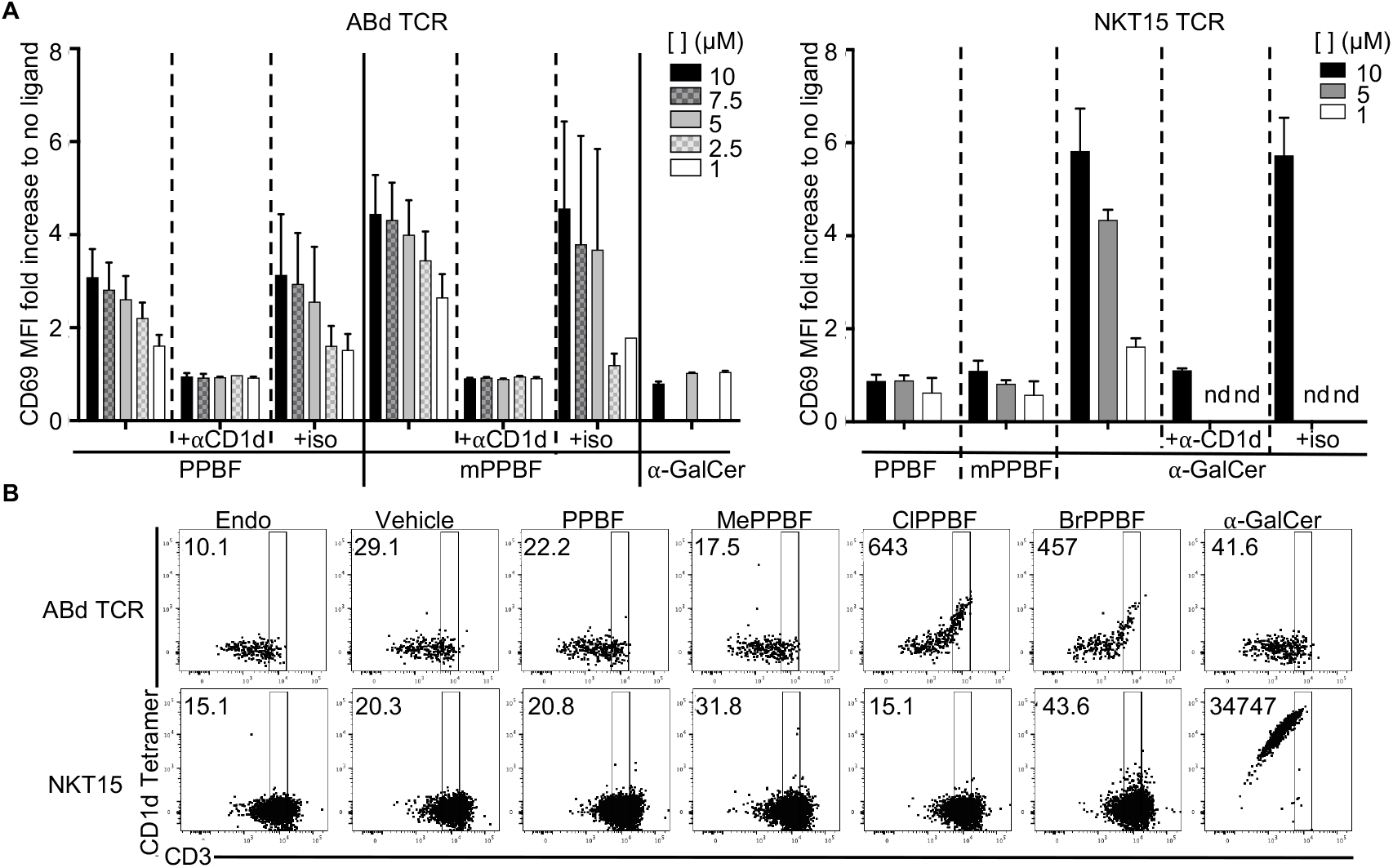
PPBF-mediated type II NKT cell activation is due to TCR recognition of CD1d-PPBF complexes. **A.** PPBF or mPPBF were incubated with C1R antigen presenting cells at concentrations ranging from 1–10 μM, and assessed for their ability to activate a SKW3.β2m^-/-^ cell line stably transduced with the ABd type II NKT TCR, in the presence or absence of αCD1d blocking antibody, or isotype control. α-GalCer and the NKT15 type I NKT TCR^+^ cell line. Graphs depict fold increase of CD69 upregulation in comparison to conditions with no ligand after 16h activation. Data is from 3 independent experiments ^+^ SEM. **B.** CD1d tetramers were loaded with candidate PPBF analogues and assessed for their ability to stain the ABd type II NKT TCR^+^ cell line by flow cytometry. α-GalCer-loaded or ‘unloaded’ (Endo)-CD1d tetramer, and the NKT15 type I NKT TCR^+^ cell line were included as controls. Numbers represent the MFI of tetramer staining of cells with similar TCR levels. Dot plots are representative of n=4 experiments performed.

We also stained the ABd TCR-transduced cell line with CD1d tetramers treated with PPBF or two of the higher potency analogues (**Figure 1B and 2B**). Similar to our previous work^15^, CD1d-PPBF or CD1d-MePPBF tetramers failed to stain the ABd-TCR^+^ cell line above background with untreated CD1d tetramers (**Figure 2B**). However, ClPPBF, BrPPBF or ClPPBC treatment enabled CD1d tetramer staining of CD3/TCR^hi^ cells (**Figure 2 and Supplemental Figure 1**). Conversely, CD1d tetramers loaded with α-GalCer stained the control NKT15 type I NKT cell line but not the ABd type II NKT cell line (**Figure 2B**). These data can potentially be explained by the ‘affinity-avidity gap’ whereby certain molecules generate complexes with MHC or CD1 proteins that are of sufficient TCR affinity to functionally activate T cells but do not mediate tetramer binding to TCRs ^26, 27, 28, 29^. Accordingly, the two less potent agonists, PPBF and MePPBF, appear to fall within the affinity-avidity gap, whereas ClPPBF, ClPPBC and BrPPBF induce formation of higher avidity CD1d complexes that permit tetramer staining. Thus, the ABd TCR directly binds CD1d and does so in a PBF-dependent manner.

### Identification of CD1d-PPBF reactive NKT cells in PBMC

Having established that CD1d-endo tetramers treated with some PBF analogs enable staining of the ABd TCR^+^ SKW3.β2m^-/-^ cells, we used CD1d-ClPPBF tetramers to search for PBF-reactive cells in PBMCs from human blood donors. Compared to the frequency of staining of type I NKT cells with CD1d-α-GalCer tetramers amongst CD3^+^CD14^-^CD19^-^7AAD^-^ human lymphocytes (0.079 to 0.23%), ClPPBF-treated CD1d tetramers stained larger numbers (~1%) of lymphocytes, but with much lower mean fluorescence intensity (MFI) **(Figure 3A)**. To enrich CD1d-ClPPBF tetramer^+^ cells with high tetramer staining to allow functional testing of TCRs and CD1d recognition, we used tetramer-associated magnetic enrichment (TAME)^30^. One round of TAME with CD1d-ClPPBF increased the frequency of CD1d-ClPPBF staining of CD3^+^ T cells (6 to 15%) and yielded more defined clusters of brightly stained populations in most cases.

**Figure 3.**
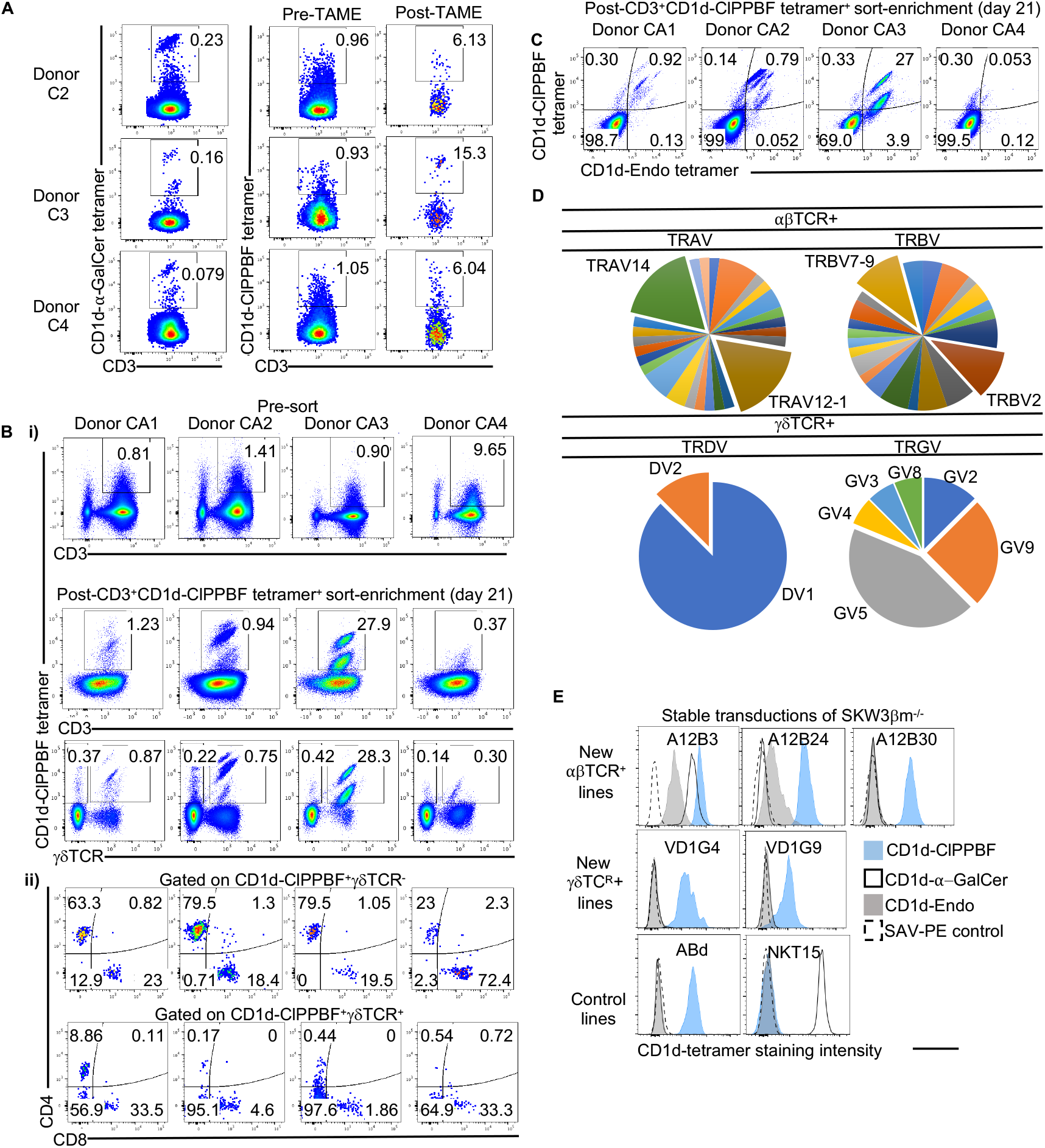
Identification of human CD1d-CLPPBF tetramer^+^ T cells in peripheral blood mononuclear cells (PBMC). **A.** Identification of CD1d–α-GalCer tetramer^+^ (type I NKT) (left), or CD1d–ClPPBF tetramer^+^ cells in PBMCs (right). Flow cytometry plots of 3 donors showing CD3 versus CD1d–α-GalCer tetramer (left) or CD1d-ClPPBF tetramer staining (right) pre- and post-CD1d-ClPPBF tetramer associated magnetic enrichment (TAME), on gated 7AAD^−^CD14^−^CD19^−^ single lymphocytes. Data is representative of *n* = 10 individual donors across 3 independent experiments. **B**. CD1d-ClPPBF^+^CD3^+^ cells were bulk sorted and expanded *in vitro* for 21 days with plate bound anti-CD3 and anti-CD28, in the presence of IL-2, IL-7 and IL-15. After expansion, samples were reassessed for their ability to bind CD1d-ClPPBF tetramers. Representative plots of 4 donors showing pre- and post-sort enriched CD3 or γδTCR versus CD1d–ClPPBF tetramer staining (i). CD4 Vs CD8 expression on αβ T cells (CD3^+^CD1d^-^ClPPBF^+^γδTCR^-^) or γδ T cells (CD3^+^CD1d-ClPPBF^+^γδTCR^-^) (ii). Plots are representative of *n*=13 individual donors across 6 independent experiments. **C.** Dual tetramer labelling showing CD1d-endo versus CD1d–ClPPBF tetramers on sort-enriched cells showing 7AAD^−^CD19^−^CD14^−^CD3^+^ cells. **D.** Single positive CD1d-ClPPBF^+^CD1d-Endo-CD3^+^ cells were single cell sorted into individual wells for TCR gene PCR amplification. Pie charts with TCR genes used were derived from analysis of 13 donors (αβTCRs) or 7 donors (γδTCRs), of 6 independent sorting experiments. **E.** SKW3.β2m^-/-^ cells were stably transduced with TCR sequences for which CD1d-reactivity had been validated in Supplemental Figure 1 and stained with anti-CD3 and ClPPBF-, α-GalCer-loaded or ‘unloaded’ (Endo)-loaded CD1d tetramers. Histograms show tetramer staining of cells previously gated to a confined range of TCR expression levels to ensure that different TCR expression between cell lines was not a variable, and are representative of n=3 independent experiments. SAV-PE control staining was also included (dashed line).

Flow cytometric cell sorting of CD1d-ClPPBF tetramer^+^ cells, followed by *in vitro* expansion in the presence of anti-CD3 and anti-CD28 plus cytokines (IL-2, IL-7 and IL-15) for 21 days, generated large CD1d-ClPPBF tetramer^+^ T cell populations (~0.3%-30% of CD3+ cells) from all donors tested **(Figure 3B).** More so than pre-sort staining, these enriched and expanded cells exhibited clear CD1d-ClPPBF tetramer staining, which correlated with CD3 intensity as expected when tetramers bind TCRs. Unexpectedly, in most donors, these cells included both αβ T cells (CD3^+^γδTCR^-^) and γδ T cells (CD3^+^γδTCR^+^) cells (**Figure 3B**), identifying the first γδ T cells to recognise PBF-treated CD1d. Most CD1d-ClPPBF tetramer^+^ αβ T cell clones were CD4^+^ or CD8^+^, whereas γδ T cells that bound CD1d-ClPPBF tetramer were CD4^-^CD8^-^ double negative (DN), typical of γδ T cells, including those described to interact with CD1d^31, 32, 33^. Thus, human PMBC from healthy donors contain populations of αβ TCR^+^ and γδTCR^+^ cells that can recognise CD1d treated with non-lipidic PBF molecules.

The above data could not distinguish between T cells that were reactive to CD1d alone from those that specifically react with CD1d in association with PBF antigen. To investigate this question, *in vitro*-expanded tetramer^+^ cells were dual-stained with CD1d-ClPPBF tetramers and CD1d tetramers that were not treated with CIPPBF, but likely still carrying endogenous lipids (‘CD1d-endo’) derived from the mammalian expression system. In all subjects (4 shown, representative of 13 donors analyzed), the 2D staining pattern showed several distinct T cell clusters, which likely represent oligoclonal expanded subpopulations of NKT cells (**Figure 3C**). Most clusters co-stained with CD1d-endo and CD1d-ClPPBF, showing a positive correlation of intensity on a diagonal, suggesting both tetramers bound the same TCR. However, in some cases, CD1d-ClPPBF tetramers stained more brightly relative to CD1d-endo, while in other cases, the opposite was observed and both examples were seen even within the same donor. Thus, the presence of ClPPBF may modulate the recognition of some self (endo) lipids present in CD1d-tetramers, or CD1d itself. Importantly, some T cells were much more strongly labelled with the ClPPBF-loaded CD1d tetramers compared to CD1d endo tetramers, appearing in the upper-left quadrant of the two dimensional FACS plot.

### Diverse but biased CD1d-ClPPBF tetramer^+^ NKT TCRs

To dissect the molecular basis of CD1d-PBF TCR reactivity, we sought to clone αβTCRs and γδTCRs from CD1d-ClPPBF-reactive T cells and measure their interactions with CD1d, self-lipids, ClPPBF and α-GalCer. Using dual tetramer staining, we sorted single cells from the upper left quadrant, that were stained with CD1d-ClPPBF tetramer at higher levels than by CD1d-endo tetramers (**Figure 2C**) from 13 blood donors for multiplex single-cell TCR sequencing^34^ to identify 47 paired αβTCR sequences and 16 paired γδTCR sequences (**Table 1, Figure 3D**). None of the sorted cells expressed the defining TRAV10–TRAJ18 rearrangement for human type I NKT cells. Of the 16 unique paired γδTCR sequences and 4 unpaired sequences (not shown), 13 displayed *TRDV1* rearrangements consistent with previous studies that described the same bias amongst CD1d-restricted γδTCR^+31, 32, 33^ or δ/αβTCRs^35^ T cells. For the αβ T cells, *TRAV14*, was observed in 8 paired and 4 unpaired TCRs and *TRAV12-1* was present in 8 clones with paired sequences and in 2 unpaired TCRα sequences. Amongst the *TRAJ* genes, *TRAJ6* and *TRAJ52* were also preferred. Further, 3 out of 8 *TRAV12-1*^+^ TCRs preferentially rearranged with *TRAJ6*. These variable and joining region gene patterns are notable because the ABd TCR uses *TRAV12-1*– *TRAJ6*^15^, suggesting a new interdonor TCR conservation pattern among human type II NKT cells (**Table 1**). Within the TCR β-chain rearrangements, there was a preference for *TRBV7-9* and *TRBV2* genes, which differed to the TRBV25 chain used by type I NKT cells (reviewed in ^1^). Together, these data suggest that CD1d-ClPPBF tetramers identify a population of T cells expressing diverse but biased αβTCRs (*TRAV12-1* and *TRAV14*) or *TRDV1*-biased γδTCRs, and that these are distinct from type I NKT TCRs.

**Table 1.**
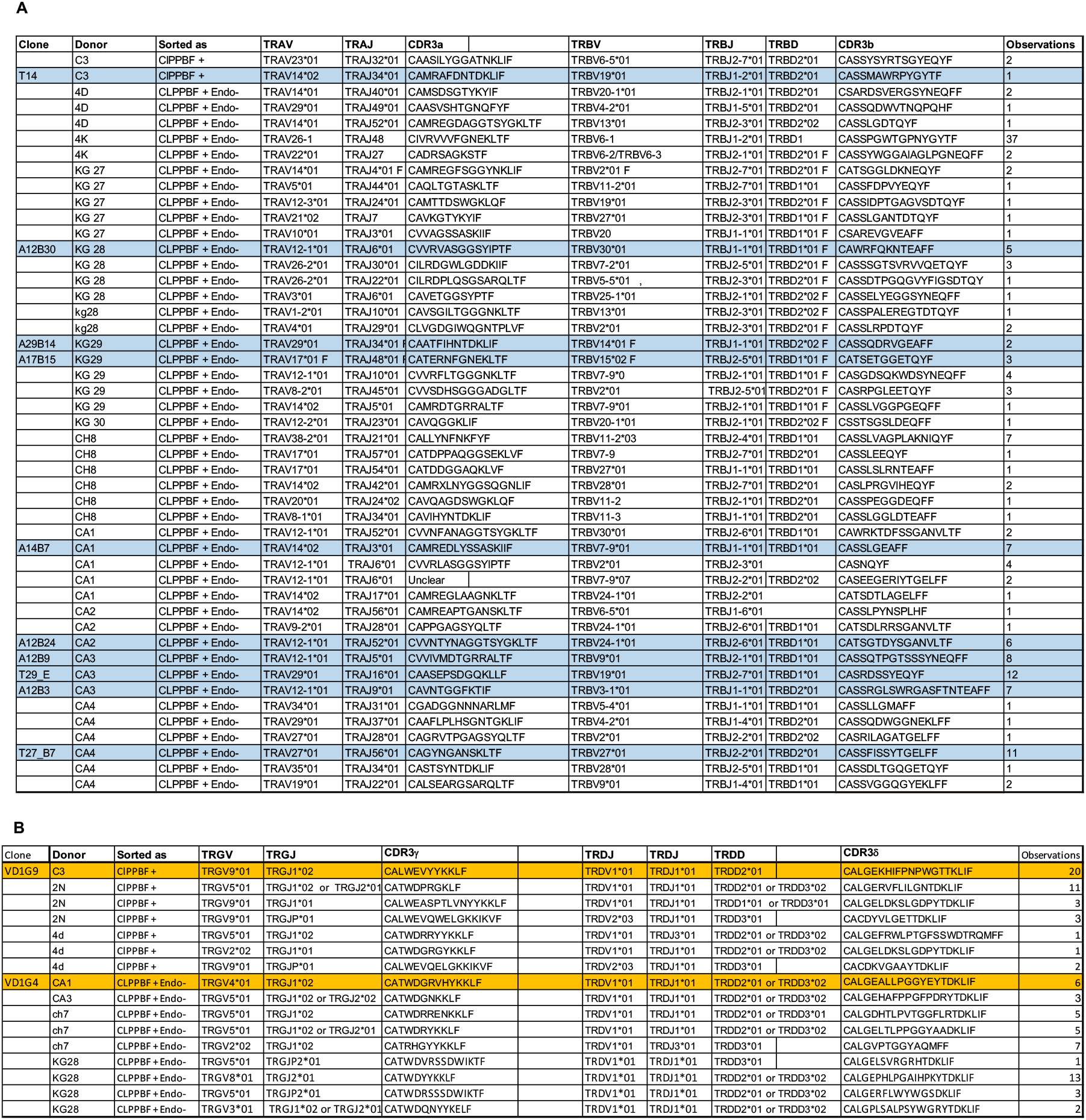
Paired sequences of CD1d-ClPPBF tetramer^+^ single-sorted cells. IMGT TCR gene nomenclatures and associated complementary determining region (CDR) 3 loop amino acid sequences are shown or αβ and γδ T cells sorted with CD1d-ClPPBF tetramers post tetramer-associated magnetic enrichment - Figure 2A (donor C3) -or as CD1d-ClPPBF tetramer^+^CD1d-Endo tetramer-after CD1d-ClPPBF tetramer sort/expansion in culture for 1 days as per figure 2C (upper left quadrant). The number of observations column refers to the frequency that each unique clonotype was observed within each experiment including analysis of 13 donors (αβTCRs) or 7 donors (γδTCRs). A total of 6 sorting experiments were performed.

### Specificity of the CD1d-ClPPBF tetramer^+^ T cells

To test the specificity of the putative CD1d-ClPPBF-reactive TCRs, we selected nine αβTCR and two γδTCR sequences, from 7 donors (**Table 1**) of which three TRAV12^+^ αβTCRs (clones A12B3, A12B24 and A12B30) and both γδTCR sequences (clones VD1G9 and VD1G4) recapitulated CD1d-ClPPBF tetramer staining using HEK293T-based TCR transfectants (**Supplemental Figure 2**). These validated TCR sequences were stably transduced into SKW3.β2m^-/-^clones that were assessed for binding to CD1d-ClPPBF, CD1d-endo CD1d-α-GalCer tetramers (**Figure 3E**) along the original PPBF-reactive ABd TCR and type I NKT TCR cell lines, which served as controls for this experiment. Three TRAV12^+^ TCRs, both γδTCR clones, and the ABd clone were stained strongly by CD1d-ClPPBF at levels above CD1d-endo tetramers or streptavidin (SAV)-control (**Figure 3E**). Despite that, some clones were also stained by CD1d-endo tetramers above SAV-control **(Supplemental Figure 2 and Figure 3E),**showing that PBFs increase CD1d-TCR binding but are not absolutely required for CD1d-TCR interaction. Conversely, the NKT15 TCR cell line was only stained by CD1d-α-GalCer tetramers. Interestingly, α-GalCer-loaded CD1d tetramers also stained the A12B3 clone more brightly than CD1d-endo, but not as brightly as CD1d-ClPPBF tetramers, suggesting that this is an atypical (type Ia) NKT TCR whose CD1d-reactivity is also enhanced by the presence of α-GalCer, similar to our previous reports ^36, 37, 38^, although to a lower extent than the presence of ClPPBF. Taken together, we define a population of CD1d-endo and PBF reactive T cells using tetramers in the circulation of healthy blood donors, which respond to CD1d and PPBF in a TCR-dependent manner.

### Potential docking mode of PPBF-reactive TCRs

Next, we investigated the likely CD1d docking modality of PPBF-reactive αβ and γδ TCRs, using the ABd and VD1G9) TCRs compared to NKT15 TCR clones, respectively. We used a panel of CD1d-transduced C1R cell lines, each individually modified by alanine substitution of residues that are distributed on the solvent exposed surface of CD1d, where TCRs have been previously shown to bind^31,39^. With the exception of the Val72 CD1d mutant, which only had 75% of the CD1d MFI surface expression of WT CD1d, all other mutant cell lines displayed similar CD1d surface expression (<25% variation to WT C1R.CD1d) (**Supplementary Figure 3**). The footprint of NKT15 TCR (**Figure 4**) is known from a ternary crystal structure^39^, and was used as a reference to interpret the effects of CD1d mutation on response to α-GalCer-loaded C1R cells. As expected, CD1d mutations affecting residues involved in binding of the type I NKT TCR to CD1d-α-GalCer such as Glu83-Ala, Val147-Ala, Lys86-Ala, Met87-Ala, which form a ring around the F′ antigen portal, had a moderate (orange) or severe (red) impact on CD69 upregulation of the NKT15 cell line. Cellular activation of both VD1VG9 and ABd TCR^+^ CD1d-PBF-reactive cell lines was influenced by the mutation of a common residue, Trp160, which sits in the central A′ roof of the α2 helix of CD1d. This mutation did not affect the activation of the type I NKT15 TCR, suggesting distinct CD1d footprints. ABd activation was altered by mutations located towards in the A′ roof (Val72, Arg79) and near the F′ roof (Lys86, Val147), which suggests that ABd TCR binds over the central position of CD1d that covers the antigen portal. When PPBF was not included, similar results were observed for both the ABd and the VD1 TCR (data not shown). Overall, the CD1d docking footprint of the VD1G9 and ABd PPBF-reactive TCRs were distinct to the F′-oriented docking of type I NKT TCRs (**Figure 4**)^39, 40^, α-GlcADAG, α-GalCer reactive (type Ia) TRAV13–TRAJ50^+^ TCRs^36, 41^, and other type II NKT TCRs^38^ or the A′-oriented docking of atypical^37^ or sulfatide-reactive type II NKT^42, 43^ and TRDV1^+^ CD1d-restricted TCRs^33^. Thus, PPBF-reactive TCRs potentially dock over the top of the antigen-binding cleft of CD1d, which might allow productive analysis of molecules that are trapped between CD1d and TCR using TCR trap technology^23, 24^.

**Figure 4.**
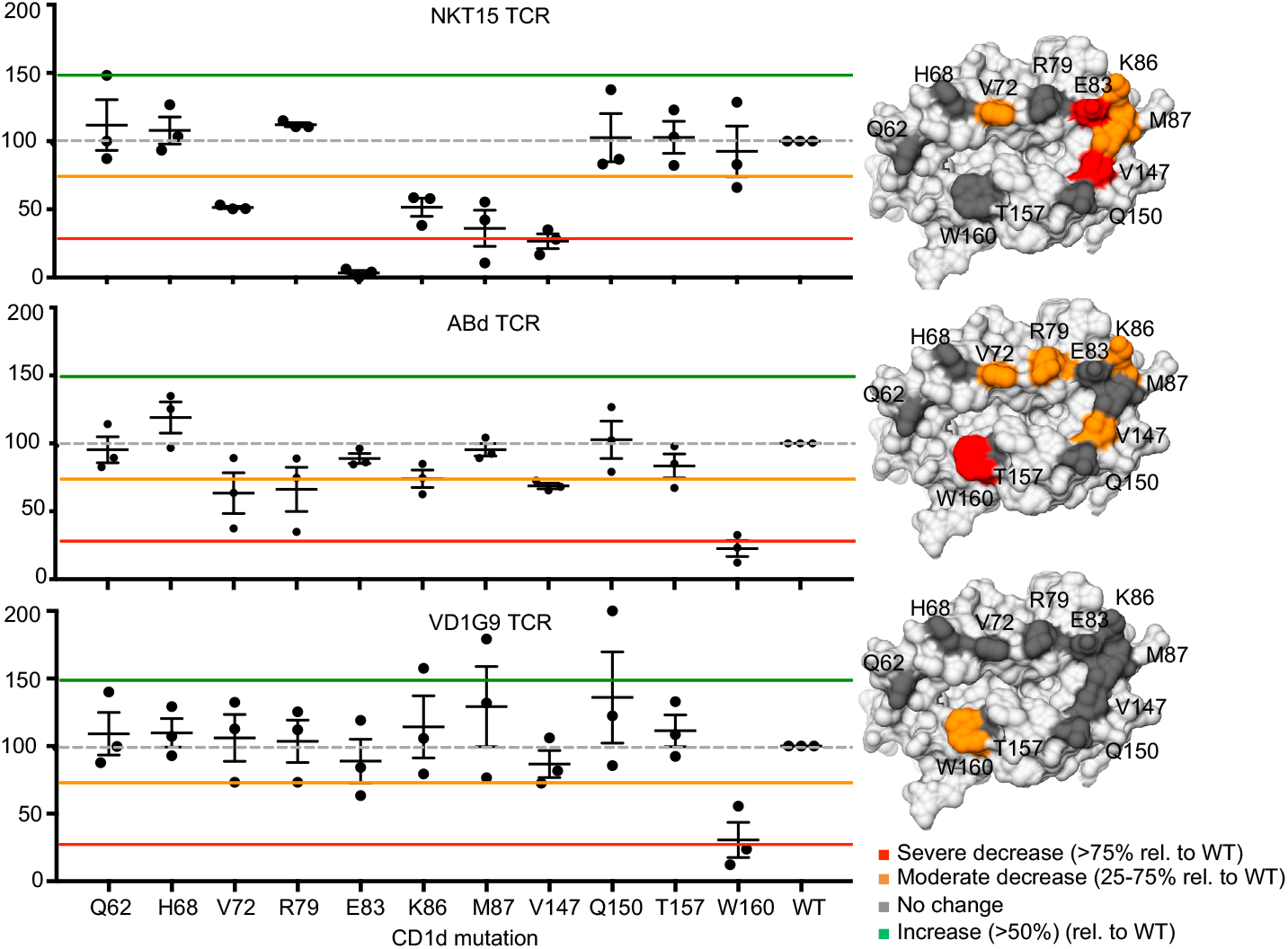
CD1d docking modes of PPBF-reactive type II NKT cells. ABd or VD1G9 NKT TCR-transduced SKW3β2m^-/-^ cell lines, were co-cultured with C1R cells each transduced with a single (Ala) mutant of CD1d. The level of activation (CD69) elicited by each mutant is normalised to the response elicited by WT CD1d. The responses of a NKT15-TCR expressing cell line to α-GalCer stimulation were also analysed as control. Graphs show the average of duplicate wells ± SEM within one experiment, where 3 independent assays were performed. Corresponding CD1d surface maps (PDB code: 1Zt4) are shown to the right of each graph, depicting residues that when mutated had no effect (dark grey).

### Trapping lipids in CD1d-PPBF-TCR complexes

Since PBFs promote CD1d-TCR binding (**Figure 3C, E**), this raises the question of whether PBFs either alter the structure of CD1d, influence lipid ligand content or are captured at the CD1d-TCR interface. To address this, we generated soluble ABd TCR to trap and identify compounds within CD1d-TCR complexes. The TCR trap approach^23, 24^ involves mixing purified recombinant CD1d proteins and TCRs followed by size exclusion chromatography to enrich for CD1d-TCR complexes, which are larger and thereby elute earlier than CD1d and TCR monomers (**Figure 5A**). Next, mass-spectrometric (MS) analysis of lipid eluents derived from early eluting protein (fractions 30-34) allows direct identification of antigen molecules within CD1d-TCR complexes, which are compared to ligands detected in eluents from CD1d monomers, and present within intermediate (fractions 35-36) and later fractions that co-elute with protein monomers (fractions 37-41) ^23, 24^. Addition of BrPPBF to mixtures of CD1d and the ABd TCR facilitated CD1d-TCR complex formation as observed by early elution of material (fractions 30-34) that resolved from the profiles of CD1d alone or TCR alone (**Figure 5A**). This supports that BrPPBF increased formation of CD1d-TCR complexes.

**Figure 5.**
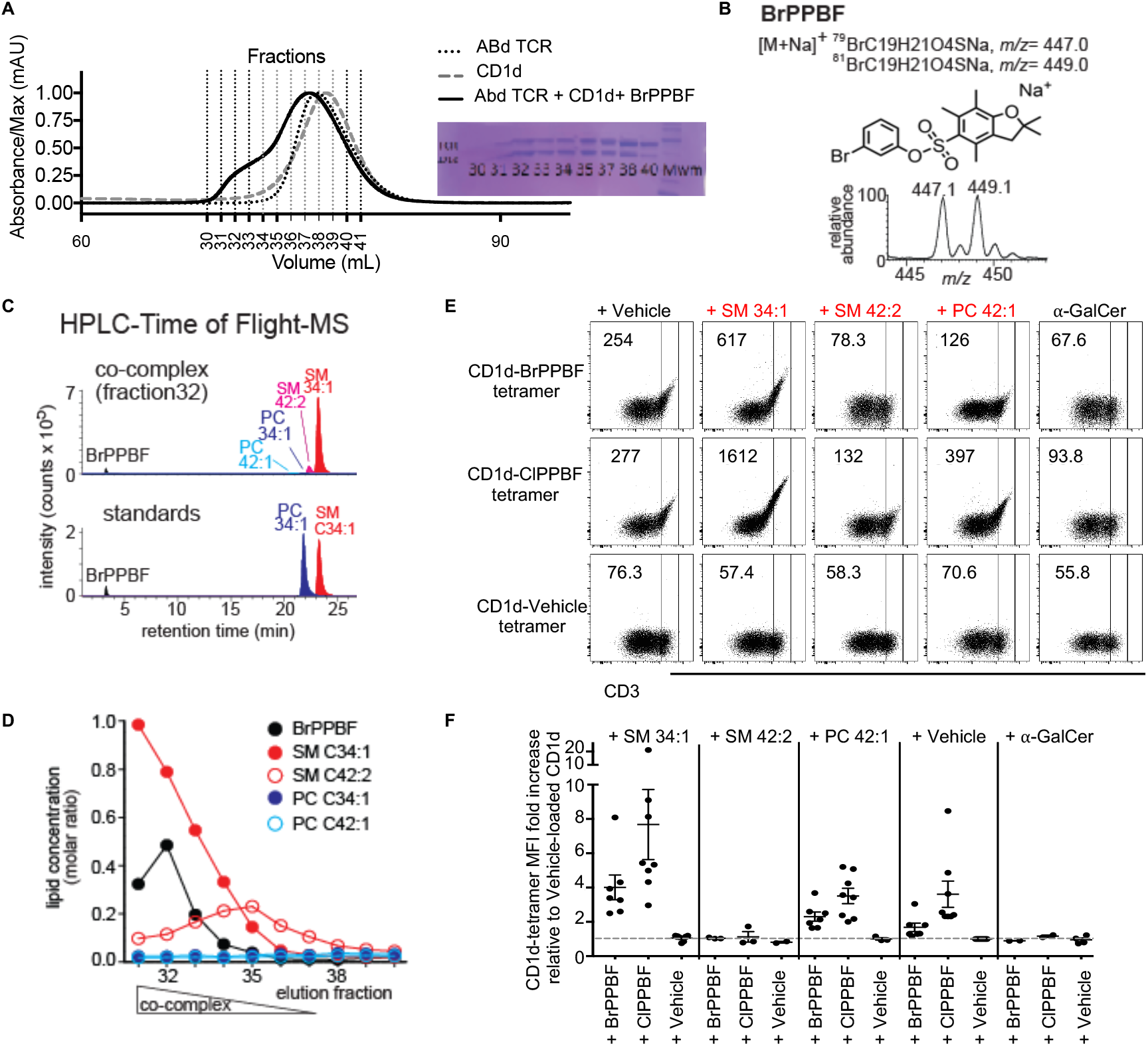
TCR-trap determination of CD1d-BrPPBF-TCR-complexes. **A.** CD1d-TCR-complexes were separated from monomers by size using fast performance liquid chromatography (FPLC) and protein composition and validated by polyacrilimde gel electrophoresis. Fractions were normalized to protein input and then subject to lipid elution and analysed by MS. **B.** BrPPBF was identified by shotgun nano-ESI MS in the early co-complex (fraction 32). **C.** Lipids extracted from the co-complex (fraction 32) were analyzed by HPLC-QTof-MS to obtain the accurate mass and retention time in comparison with the indicated standards. **D.** The relative quantity of 5 detected lipids in the TCR-trap fractions were estimated by fitting the chromatogram area to an external standard curve. **E.** ABd TCR^+^ cell line was stained with CD1d-ClPPBF or BrPPBF pre-loaded with short chain 34:1 SM or long chain 42:2 SM or 42:1 PC. Representative flow cytometry plots and MFI of CD1d tetramer staining of cells with similar TCR levels (top). The fold increase in CD1d tetramer MFI is shown with each combination of lipid and PBF (bottom). Each point represents one individual experiment.

### Nano-electrospray ionization of lipids in CD1d-TCR complexes

Sensitive shotgun nano-electrospray MS analysis of the trace lipids eluted from proteins generated an overview of the compounds present TCR-CD1d complexes (fraction 32), intermediate (fraction 35) and non-complexed (fraction 38) fractions (Supplemental **Figure 4 A-B**). Diagnostic ions corresponding to the sodium adduct of BrPPBF (*m/z* 447.1 and 449.1) and matching the 1:1 isotopic ratio of ^79^Br and ^81^Br were detected in fraction 32 (**Figure 5B**), demonstrating BrPPBF within the CD1d-TCR complexed fraction. CD1d-endo complexes also carry endogenous lipids acquired during synthesis in the mammalian expression system that we used, which were identified in all fractions (**Figure 5B, Supplemental Figure 4B**). Collision induced dissociation (CID) MS revealed a prominent ion at *m/z* 725.4 from the CD1d-TCR complex fraction identified as short-chain sphingomyelin (SM) with a combined chain length of C34 with one unsaturation (34:1 SM) (**Supplemental Figure 4B-D**). We also detected weak signals for two phosphatidylcholines, 34:1 PC and 42:2 PC (*m/z* 782.4 and 894.6) **(Supplemental Figure 4B-C)**, and 40:2 SM and 42:2 SM (*m/z* 807.5 and 835.5) **(Supplemental Figure 4B)**. While 34:1 SM was enriched within CD1d-TCR fractions, the other ions were enriched in uncomplexed fractions (**Figure 4B)**, indicating BrPPBF is both within the CD1d-TCR complex and influences the spectrum of endogenous lipids found within the TCR-CD1d complex.

### Quantitative analysis of trapped lipids by HPLC-MS

We used high performance liquid chromatography (HPLC)-MS, which provides higher mass accuracy and uses chromatography to assess the retention time of eluted ligands versus standards. This approach confirmed the identity of BrPPBF and 34:1 SM in the CD1d-TCR complex fractions, as well as trace amounts of 42:2 SM, 34:1 PC and 42:1 PC (**Figure 5C**). Time of flight (TOF) HPLC-MS detection allows quantitation of these molecules using external standards (**Figure 5D**, **Supplemental Figure 5**). These measurements demonstrated a clear pattern for selective capture of 34:1 SM and BrPPBF in early co-complex fractions. 42:2 SM and PCs were present only at low levels and showed modest variations, but less clear enrichment patterns than seen with 34:1 SM (**Figure 5D**). Importantly, control runs of CD1d alone indicated that these patterns of lipid or BrPPBF enrichment in early fractions did not occur in the absence of TCR (**Supplemental Figure 6**), so fractionation results from lipid/BrPPBF entrapment by the TCR rather than direct interaction with the column.

To investigate whether BrPPBF was required for CD1d-TCR interaction, we repeated the TCR trap experiment in the absence of BrPPBF. Lower levels of complexation were observed, with much less material obtained in the earliest fractions (**Supplemental Figure 6**). Both 34:1 SM and 42:1 SM were detected in early CD1d-TCR complex fractions, but the ratio of short (34:1) to long (42:2) SM was markedly increased in the early fractions, in the presence of BrPPBF (**Supplemental figure 4B**). Overall, these experiments indicated that BrPPBF promotes early elution of CD1d-TCR complexes, but that short chain sphingomyelin can also appear in CD1d-TCR complexes. Finally, we studied whether 34:1 SM augments CD1d tetramer staining when added as an exogenous ligand. For CD1d-endo tetramers, exogenous 34:1 SM alone did not enhance staining of the ABd TCR (**Figure 5E**) or the VD1G9 TCR (**Supplemental Figure 7**). However, when CD1d tetramers were co-loaded with BrPPBF or ClPPBF and 34:1 SM, staining augmentation was observed suggesting synergistic interaction of BrPPBF with a self-lipid. Conversely, co-loading of BrPPBF or ClPPBF with 42:2 SM led to inhibition of CD1d tetramer staining, when compared to co-loading with vehicle, whilst co-loading 34:1 PC had no impact. These results suggest that the ability of PBF molecules to enhance CD1d-TCR interactions is lipid-sensitive.

## Discussion

Whereas most studies of CD1 emphasize T cell recognition of lipids comprised of alkyl chains, this study provides a different view of NKT cell activation by constrained, non-lipidic small molecules in the benzyl sulfonate family. Further, this work extends the diversity of the type II NKT cell family by defining a new public TCR motif among polyclonal T cells and provides detailed information about the identity and action of non-lipidic NKT cell agonists. Whereas initial views favored an indirect or pharmacological action of these atypical molecules, we identified and characterised a polyclonal population of NKT cells that can recognise, via their TCR, CD1d molecules associated with an exogenously loaded non-lipidic PBF molecules and an endogenous self-lipid, such as 34:1 SM. Overall polyclonal αβ and γδ T cells reactive to CD1d and PBF are present among type II NKT cells in all donors tested.

Notably one of the biased TCR motifs (TRAV12–TRAJ6) that emerged among type II NKT cells resembled the TCR from the original ABd clone isolated 17 years prior to this study^15^. This finding and the monomorphic nature of CD1d hint at conserved modes of interaction of the TCR-PBF-CD1d interaction. The ABd TCR and polyclonal TCRs stained by CD1d-ClPPBF tetramers showed tolerance to some, but not all, variations in the PBF molecular structure, suggesting that the 3-substituted phenyloxysufonyl motif is critical for supporting the TCR-CD1d interaction, while the nature of the benzofuran/chroman system is more flexible.

Models of glycolipid display by CD1d generally predict that the two alkyl chains present in most glycolipid antigens occupy the two pockets of CD1d, and the carbohydrate head group protrudes for TCR recognition^44^. However, PPBF variants are small (*m/z* 350-400) and rigid molecules constrained by one phenyl and one sulfonyl group; these non-lipidic molecules lack alkyl chains and head groups and therefore do not neatly fit within the amphipathic lipid-display model paradigm. Although the precise mode of lipid and small molecule positioning on CD1d awaits further structural confirmation, our data provide specific insights into candidate mechanisms. Firstly, our data demonstrate that PBFs are not mitogens that directly or broadly activate T cells. Instead CD1d tetramers and TCR^+^ cell line assays indicate that PBFs are directly involved in promoting CD1d-TCR interactions. However, some T cells identified using PBF-loaded CD1d tetramers did not absolutely require PBF for tetramer staining or CD1d-TCR complex formation, suggesting that they have basal reactivity towards CD1d presenting endogenous-ligands that is enhanced in the presence of PBFs. Intriguingly, these findings are extended by the unbiased discovery of molecules eluted from a CD1d-αβTCR complex, which detected BrPPBF and a self-lipid 34:1 SM in eluates. This finding shows that PBF both increases CD1d-TCR complex formation and places a PBF present within the CD1d-TCR complex, raising the question of whether PBF and 34:1 SM can jointly or separately support TCR binding. The TCR trap assay without BrPPBF also revealed an enrichment for a short chain SM in early eluting fractions where CD1d-TCR complexes are likely to emerge, suggesting that a self-lipid can support CD1d-TCR binding at a threshold that does not detectably activate or enable CD1d tetramer staining of NKT cells such as the ABd clone.

The action of PBFs to increase CD1d-mediated activation, tetramer binding and TCR trapping/co-complexation are subject to two possible interpretations to address the key question of whether PPBF directly engages the TCR at the binding interface, or whether it indirectly acts on CD1d and/or endogenous lipids. One is based on the fact that stimulatory SMs and PBFs are both relatively small molecules. The mass of PBFs (~400 amu) is about half the size of most lipids bound to human CD1d (~800 amu, ^45^), and the ratio of 34:1 SMs to 42:2 SMs is increased in complexes. Recent studies show how small ‘permissive’ ligands buried within CD1a can expose the CD1a surface for direct recognition^46, 47, 48, 49, 50^. In this interpretation, the reactivity of ABd and other TCRs might be explained by the small size of short-chain SM and PBF molecules, allowing them to bind largely within the CD1d cleft to expose the outer surface of CD1d to a TCR. Consistent with this interpretation, footprint analysis shows that one αβ and one γδ TCR binds over the F’ portal of CD1d where larger ligands would protrude. However, this analysis does not explain why SM alone does not lead to T cell staining with CD1d tetramers, whereas PPBF and SM (C34:1) synergise to enhance the TCR-CD1d interaction. The other interpretation is a combined action since with the TCR trap experiments show that both SM and PBF were in CD1d-TCR complexes, although this does not necessarily mean that the two molecules are present together within the same complexes. However, it is possible that the ABd TCR is recognising a trimolecular complex comprising CD1d, SM (C34:1) and PPBF, which is also consistent with the combination of these two ligands enhancing CD1d tetramer staining of PPBF-reactive cell lines. In this model, the TCR may directly contact both SM or PPBF, or, SM alters the orientation of PPBF, or vice versa, in order to create a better TCR epitope with one or the other.

Modulation of type I NKT cell responses to CD1d-lipid antigen complexes through lipid-tail-^51^ or -headgroup modifications^52, 53, 54^ has been extensively studied and can impact on immune response and disease outcomes^55^. Our results show that a family of structurally related non-lipidic small molecules can also modulate CD1d-dependent T cell responses and could be used as drugs to transiently modulate CD1d-mediated T cell responses for immunotherapy. Finally, PBF compounds share chemical features with commonly used sulfa-drugs that generate hypersensitivity reactions in humans, including sulfisomidine, sulfadiazine, sulfasalazine, and celecoxib^16^. In a similar vein, small lipid allergens can enhance CD1a-mediated T cell autoreactivity by displacing certain lipids from CD1a and altering the remaining lipid repertoire, enhancing CD1a-autoreactivity by some T cell clones^49^. Given that hypersensitivity to sulfa-drugs limits the use of these otherwise valuable therapeutics^18, 19^, these data support investigation of whether PBF-like sulfa-drugs can bind CD1 proteins, and whether there is a role for type II NKT cells in sulfa-drug induced hypersensitivity.

## Materials and Methods

### Flow cytometry

Buffy coats from blood donors (Australian Red Cross Blood Service agreement 13-04VIC-07) were studied in accordance with the University of Melbourne Human Research and Ethics Committee (approval number 1035100). PBMCs were prepared by density gradient centrifugation Ficoll-Paque (GE Healthcare). Cells were first incubated for 10 min on ice with Fc-receptor block (BD Pharmingen), followed by CD1d-ClPPBF tetramers for 30min. Where described, CD1d–ClPPBF tetramer^+^ cells were then enriched by TAME, using anti-phycoerythrin (PE) magnetic beads and LS columns (Miltenyi Biotec), followed by surface antibody staining and/or by secondary CD1d tetramer staining (CD1d-Endo tetramers). Antibodies used include: CD3e (UCHT1, eBioscience and Becton Dickinson), CD4 (RPA-T4, BD Pharmingen), CD8α (SK1, BD Pharmingen), CD19 (HIB19, BioLegend), CD14 (MφP9, BD Pharmingen), TCRγδ (11F2, BD Pharmingen), and 7-aminoactinomycin D (7AAD) viability dye (Sigma). Antibody dilution factors were empirically determined. CD3^+^ CD1d–ClPPBF tetramer^+^ cells were sorted using a FACSAria (BD Biosciences). Where described, sorted were then expanded for 14–21 days using anti-CD3 (10 μg/mL, UCHT1, BD Pharmingen), anti-CD28 (2 μg/mL, CD28.2, BD Pharmingen), IL-2 (20 U/mL, Prepotech) IL-7 (5 ng/mL, eBioscience) and IL-15 (40 pg/mL), in complete RPMI-1640 supplemented with 10% (v/v) FBS (JRH Biosciences), penicillin (100 U/ml), streptomycin (100 μg/ml), Glutamax (2 mM), sodium pyruvate (1 mM), nonessential amino acids (0.1 mM), HEPES buffer (15 mM, pH 7.2-7.5) (all from Invitrogen, Life Technologies) and 2-mercaptoethanol (50 μM, Sigma). Expanded cells were re-stained with CD1d-ClPPBF tetramers alone, followed by surface antibodies and secondary (CD1d-endo) tetramer staining. CD1d-ClPPBF tetramer^+^ cells were then index-sorted as single cells for TCR sequencing. Data analysis was completed using FlowJo (BD), and graphs generated using GraphPad Prism.

### CD1d production

Soluble human CD1d was produced in mammalian HEK-293S.N acetylglucosaminyltransferase-I−(*GnTI−*) cells by co-transfection with pHLsec vectors encoding truncated human CD1d ectodomain with a C-terminal biotinylation motif and His6-tag (amino acid sequence at the C-terminus: GSGLNDIFEAQKIEWHEHHHHHH) and β2-microglobulin, using polyethylenimine as described CD1d^56^. Soluble CD1d was purified from culture supernatant by immobilised metal-affinity (Ni/NTA) chromatography and size-exclusion chromatography. For CD1d tetramer generation, CD1d was enzymatically biotinylated using biotin ligase (produced in-house), and further purified by size-exclusion chromatography, followed by storage at −80 °C. Following ligand-loading (as described below), CD1d was tetramerised using SAV-PE, or SAV-BV421 (BD Pharmingen).

### PPBF and analogues

PBF analogues were synthesized by the reaction of 2,2,4,6,7-pentamethyldihydrobenzofuran-5-sulfonyl chloride or 2,2,5,7,8-pentamethylbenzochromane-6-sulfonyl chloride and an appropriate phenol or aniline with triethylamine as base in CH_2_Cl_2_ at 0°C.The solution was stirred for 24 h and then concentrated. The residue was purified by flash chromatography (ethyl ether/pet. sprits) to afford the sulfonate or sulfonamide.

### Lipids

24:1 (PBS44) was kindly provided by P. Savage (Brigham Young University). α-GalCer 26:0 was supplied by Alexis Biochemicals, and SM (42:2), SM (34:1), PC (42:1); from Avanti Polar Lipids. CD1d-ligands were dissolved in tris buffer Saline (TBS) alone (pH8) or TBS containing 0.05% v/v Tyloxapol (Sigma), or buffer containing 0.5% v/v tween-20, 57 mg/ml, sucrose and 7.5 mg/ml histidine, and incubated with CD1d at 6-12-fold molar excess overnight. Lipid-incubations occurred with CD1d at pH8 (RT), whilst non-lipid (eg. PPBF and PBF analogues) incubations occurred with CD1d in pH6 at 4°C, as this was determined to provide optimal tetramer staining. For co-loading, CD1d was first loaded with lipids at pH8 (RT) overnight followed by PPBF at pH6 (4deg) overnight. All CD1d-ligands were stored at −20C and sonicated for ~30 min prior to each use.

### TCR identification

CD3^+^CD1d–ClPPBF tetramer^+^ cells, were single-cell sorted from CD1d–ClPPBF tetramer-enriched or *in vitro* expanded NKT cells (as described above), and cDNA generated using 0.1% Triton X-100 (Invitrogen) and SuperScript VILO, following manufacturer’s instructions. Paired TCRα and TCRβ or TCRγ and TCRδ chain transcripts were amplified as previously described^57^. PCR products were sequenced (AGRF, Australia) and analysed using IMGT. TCR nomenclature is presented in accordance with the IMGT guidelines^58^. Unproductive TCR gene rearrangements were excluded from analysis.

### Production of β2-microglobulin KO SKW3 cell lines

To knock out β2m in SKW-3 cells, the short guide RNAs (sgRNAs) targeting the signaling peptide (5’-GAGTAGCGCGAGCACAGCTA - 3’) was cloned into lentiCRISPR v2 (a gift from Feng Zhang; Addgene plasmid # 52961; http://n2t.net/addgene:52961; RRID:Addgene_52961) as described^59^. Cells were transduced, selected based on puromycin resistance, then single-cell cloned. These were selected based on lack of CD1b/d surface expression, and β2m knockout was validated by immunoblotting.

### Generation of stable cell lines

TCR constructs containing full-length TCRα and TCRβ or TCRγ and TCRδ chains separated by a 2A-cleavable linker were produced (Genscript) and cloned into the pMIG II plasmid. TCR-deficient SKW3.β2m^-/-^ (SKW3 cells in which the β2-microglobulin gene had been deleted) were retrovirally transduced with both TCR and a 2A-cleavable human CD3 construct^60^, the packaging vectors pEQ-Pam-3-E pVSV-G and using HEK293T cells as packaging cells in the presence of FUGENE 6, as previously described^61^. The pMIG II expression and packaging vectors were provided by Dr. Dario Vignali (St. Jude’s Research Hospital, USA), and the CD3 expression vector was provided by Prof. Stephen Turner (Monash University, Australia). CD3(GFP)^hi^ cells were sorted and assessed for their ability to bind CD1d tetramers using flow cytometry. For assays with transiently transfected TCRs, TCR- and CD3-encoding pMIG II vectors were used to transiently transduce HEK293T cells using FUGENE (Promega). Cells were harvested after 48h and assessed for CD3/TCR expression and CD1d-tetramer binding using flow cytometry. C1R cells were transduced to express human wild type CD1d or a mutated version of CD1d (each line containing a single CD1d mutation as per Figure 4 and Supplementary Figure 3, akin to TCR-transduced SKW3.β2m^-/-^ cells, and purified by flow cytometric sorting to produce stable cells lines with similar levels of surface-expressed CD1d (Supplementary figure 3).

### Stimulation assays

TCR-expressing SKW3.β2m^-/-^ cells were co-cultured overnight in complete RPMI (as described above), with or without C1R cells (either C1R WT or C1R.CD1d transduced) or THP-1 cells, with graded concentrations of lipid in round-bottom 96-well plates, and TCR-transduced cells were analysed after 16h by flow cytometry for CD69 upregulation using, anti-CD69 (FN50, BD Pharmingen).

### Generation of soluble TCRs

Individual TCRα and TCRβ of the ABd TCR were synthesized (Integrated DNA Technologies) and cloned into the pET-28 or pET-30 vectors (Novagen), respectively. Sequence verified TCR-containing vectors were expressed and produced as inclusion bodies (IBP) in *Escherichia coli* BL21 (DE3) pLysS. Soluble TCRα- and TCRβ-chain IBPs were injected into refold buffer (containing 0.1 M Tris, 2 mM EDTA, 0.4 M arginine, 0.5 mM oxidised glutathione, 5 mM reduced glutathione, and 5 M urea, with a final pH 8.5) to allow refolding overnight, with gentle stirring, at 4C)^25^. Samples were dialysed for 4 h into 100 mM Urea, 10 mM Tris–HCl pH 8.0 followed by two consecutive dialysis into 10 mM Tris–HCl at pH 8.0 (first for 4 h and second overnight). Refolded TCRs were purified by diethylaminoethanol (DEAE) sepharose anion exchange, followed by Superdex-75 16/60 gel-filtration (GE healthcare), anion exchange Mono Q 10/100 GL (GE healthcare), and hydrophobic interactions with a HiTrap Phenyl HP column (GE healthcare. The individual TCR δ- and γ-chains of the VD1G9 TCR were ordered (IDT), cloned into a pHLSEC vectors (Novagen), expressed in HEK-293S.N acetylglucosaminyltransferase-I−(*GnTI−*) cells and purified by Ni/NTA chromatography, size exclusion chromatography and anion exchange chromatography.

### Lipid elution and Mass Spectrometry

Proteins separated by the size exclusion column were collected and subjected to lipid extraction by chloroform, methanol, and water according to the method of Bligh & Dyer ^62^.The lipid containing organic phase was separated and collected by centrifugation at 850 g for 10 min. The organic solvent was dried under a nitrogen stream and the residue was redissolved in chloroform/methanol (1:2) and stored at −20C. For non-quantitative lipid detection, the extracted lipids were analyzed by nano ESI-MS using LXQ (Thermo Scientific linear ion trap mass spectrometer. The spray voltage and capillary temperature were set to 0.8 kV and 200C. The collision energy for collision-induced dissociation tandem mass spectrometry (CID-MS/MS) was 20-30% of maximum and product ions were trapped with a q valve of 0.25. To obtain accurate mass, retention time, and relative amount information, the samples were normalized based on the input proteins (10 μM) and 20 μl were injected to an Agilent 6520 Accurate-Mass Q-TOF spectrometer equipped with a 1200 series HPLC system using a normal phase Inertsil Diol column (150 mm × 2 mm, GL Sciences), running at 0.15 ml/min according to the published method with minor modifications^63^. The quantification of BrPPBF, PC, and SM was estimated by comparison of the detected chromatogram peak area to an external standard curve fitted with quadratic equation.

## Supporting information

Chemistry Experimental

## Acknowledgments

We are grateful to Dr. Paul Savage (Brigham Young University, UT, USA) for providing α-galactosylceramide analogue PBS44 used for production of CD1d-α-GalCer tetramers. We thank staff from the flow cytometry facilities at the Department of Microbiology and Immunology at the Peter Doherty Institute and the Melbourne Brain Centre at the University of Melbourne. This work was supported by the Australian Research Council (ARC; DP170104386 and CE140100011), the National Health and Medical Research Council of Australia (NHMRC; 1113293, 1083885, 1145373), the NIH (R01 AR048632), the Allergy and Immunology Foundation of Australia (AIFA; 2021) the University of Melbourne (UoM ECR grant; 2021). DGP is supported by a CSL Centenary Fellowship. DIG was supported by an NHMRC Senior Principal Research Fellowship (1117766).

