## Supplemental Figures for "Benzofuran sulfonates and small self-lipid antigens activate type II NKT cells via CD1d"

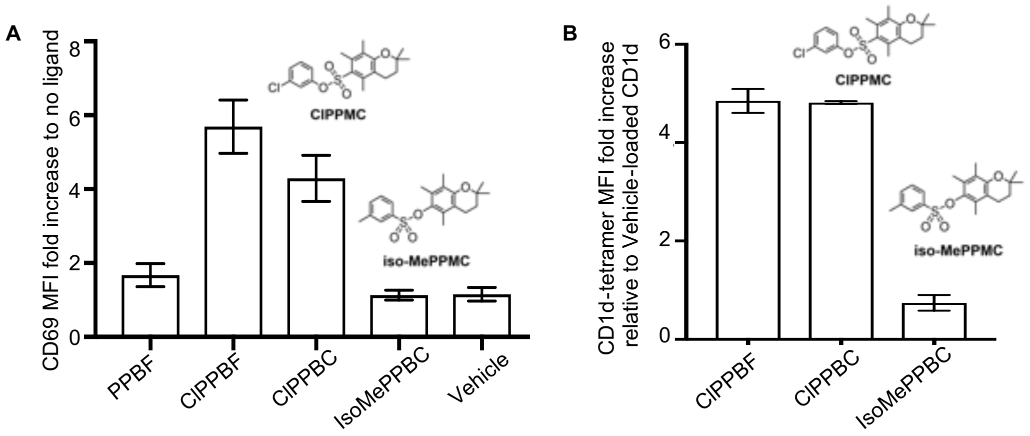


**Supplemental Figure 1 Additional structural variants: sulfonate linker orientation is important for ABd TCR activation. A**. PPBF, a ring expanded analogue of ClPPBF, ClPPBC and an isomer of MePPBC where the orientation of the $SO_3$ linkage has been inverted, IsoMePPBC, were assessed for their ability to activate ABd type II NKT $TCR^+$ cell line when co-cultured with THP-1 cells at 10μM. Graphs depict fold increase of CD69 upregulation in comparison to conditions with no ligand after 16h activation. **B.** CD1d tetramers loaded with ClPPBF, ClPPBC or IsoMePPBC were assessed for their ability to stain the ABd cell line by flow cytometry. Graphs show MFI fold increase of tetramer staining relative to vehicle-loaded CD1d of cells with similar TCR levels. Data is from 3 independent experiments + SEM.

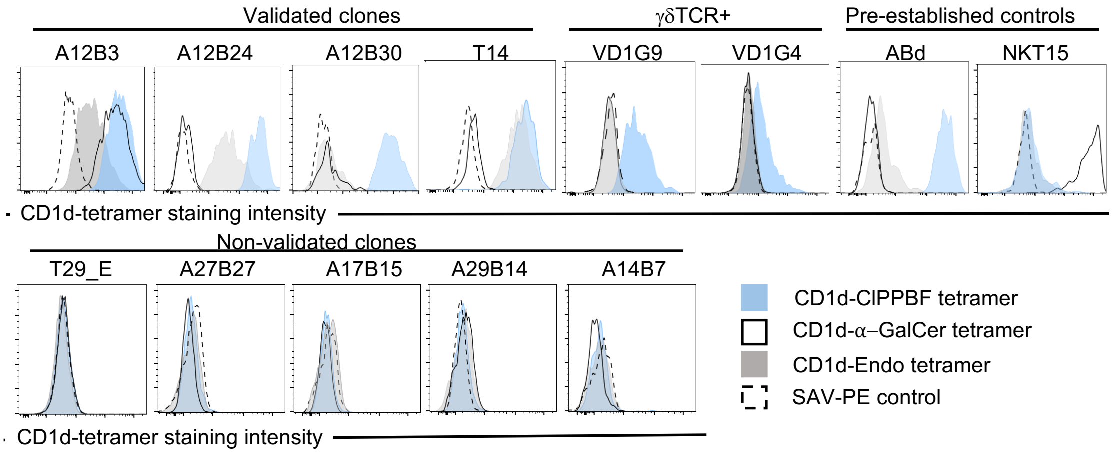


**Supplemental Figure 2. Validating the CD1d-reactivity of the TCRs expressed by CD1d-ClPPBF tetramer$^{+}$ sorted cells.** HEK 293T cell lines were transiently transfected with PMIG-constructs with selected TCR sequences from Table 1 and stained with anti-CD3 and ClPPBF-, α-GalCer-loaded or unloaded (Endo)-CD1d tetramers. Histograms depict tetramer staining of cells with similar TCR levels. SAV-PE control staining was also included (dashed line).

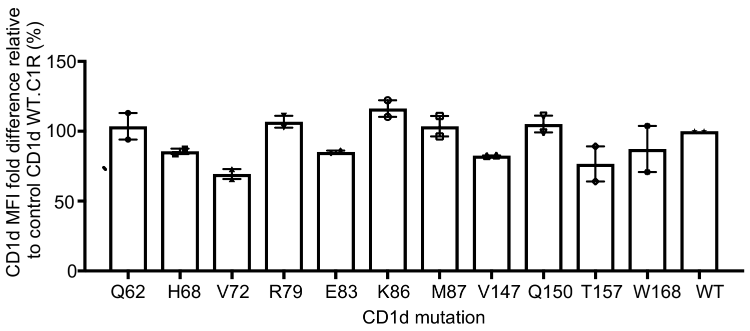


**Supplemental Figure 3. CD1d expression amongst CD1d mutant C1R cell lines**. CD1d expression C1Rs transduced with single Alanine mutated version of CD1d was assessed by flow cytometry. Graphs show CD1d MFI fold difference to WT CD1d-transduced C1Rs (CD1d WT.C1R). Data is from 2 independent experiments ± SEM.

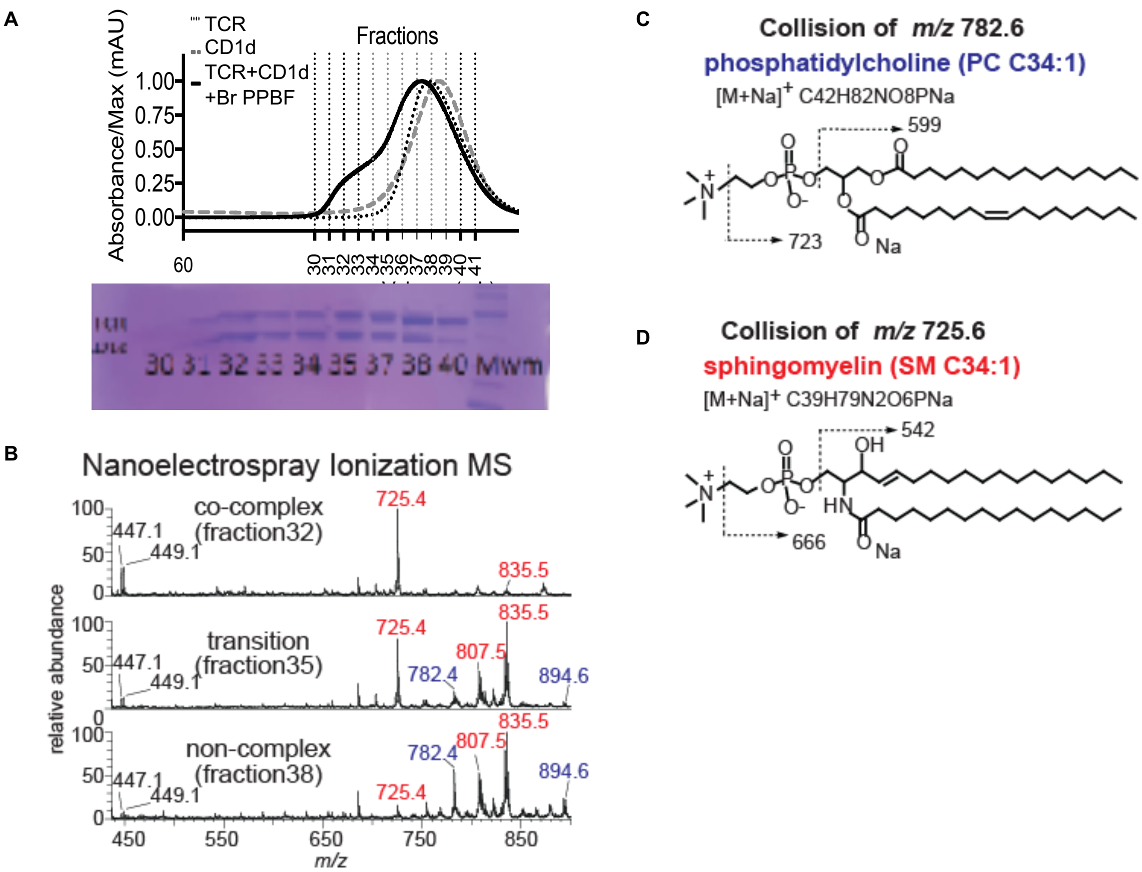


**Supplemental Figure 4. Nano-ESI MS identification of lipids in the TCR trap. A.** Polyacrylamide gel electrophoresis of protein composition of CD1d-BrPPBF-TCR fractions generated by fast performance liquid chromatography (FPLC). **B.** Lipid extracted from the TCR trap fractions indicated in A were analyzed by nano-ESI MS shotgun MS analysis to detect BrPPBF (*m/z* 447.1 and 449.1) phosphatidylcholine (PC) with an overall chain length of 34:1 (*m/z* 782.6) and 42:1 (*m/z* 894.6) as well as sphingomyelin (SM) with a chain length of 34:1 (*m/z* 725.6) 40:2 (*m/z* 807.5) or 42:2 (*m/z* 835.5) were identified in the early co-complex (fraction 32), middle transition (fraction 35), or late non-complex (fraction 38) fractions. **C-D.** The identities of PC and SM were established by CID-MS.

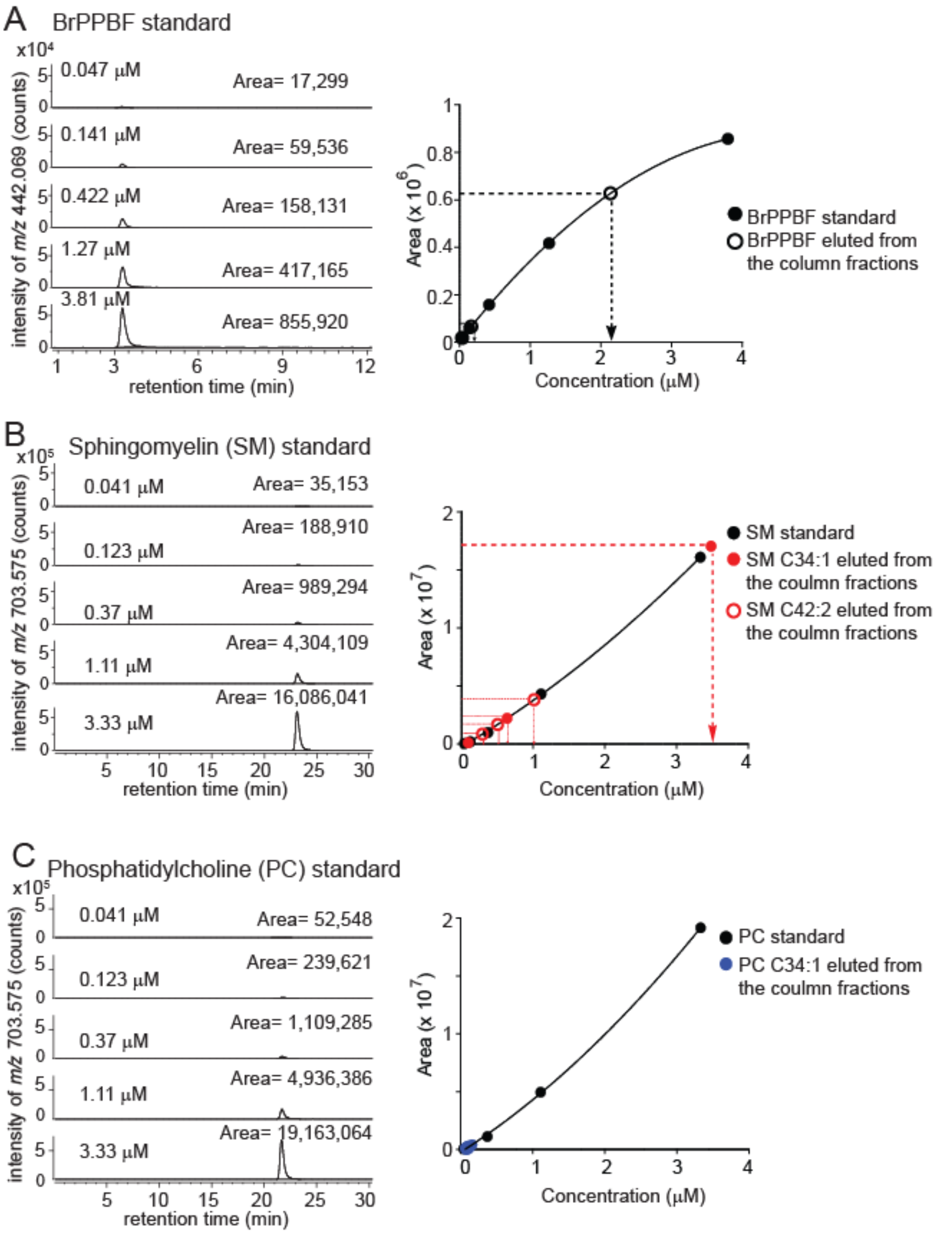


**Supplemental Figure 5. Mass spectrometry-based quantitation of lipids eluted from TCR-BrPPBF-ABd-TCR-complexes.** Serial dilutions of authentic lipid standards for BrPPBF (**A**), SM (**B**) , and PC (**C**) were injected for HPLC-TOF-MS to generate ion chromatograms from which the area under the ion chromatogram is expressed in arbitrary units (counts, left panel, solid circles), which were used to generate external standard curves from which the measured count values from CD1 or CD1-TCR eluted lipids were used to estimate absolute lipid concentrations (dotted lines) and molar ratios.

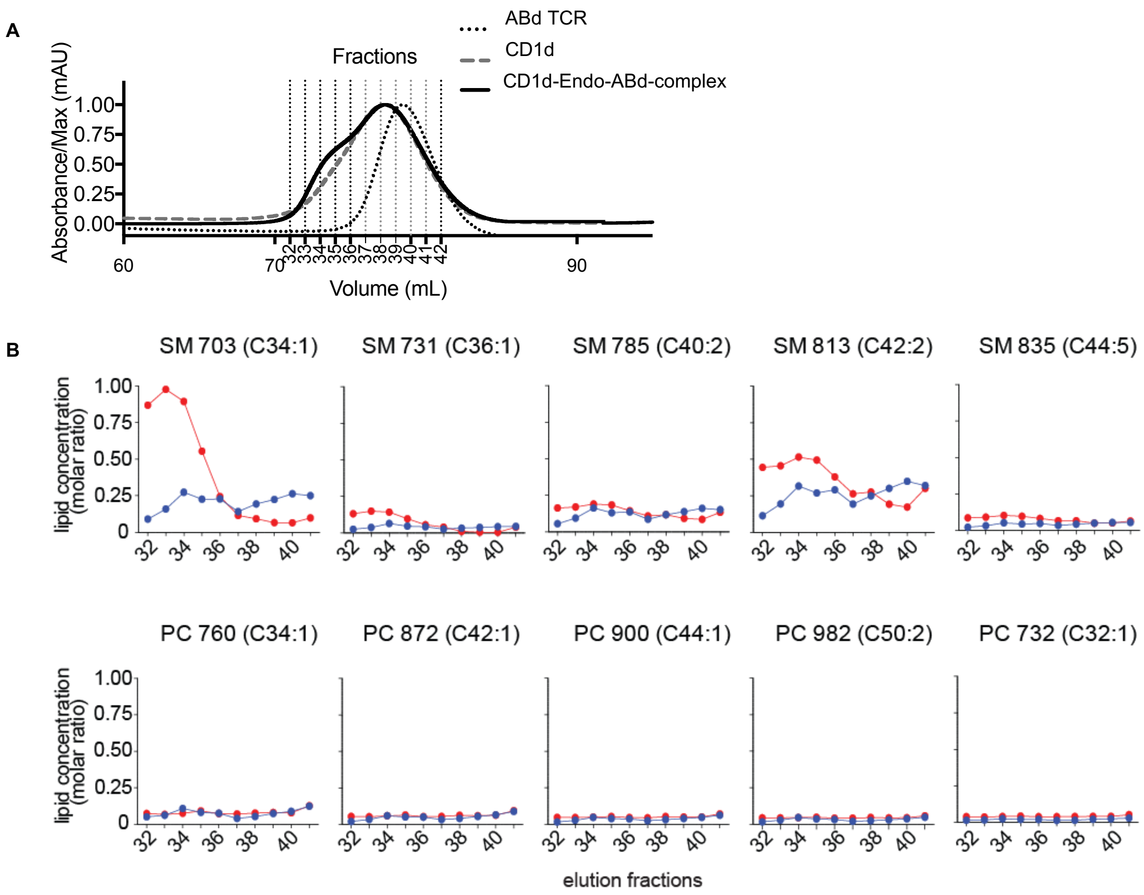


**Supplemental Figure 6. HPLC-TOF- MS analysis of lipids eluted from CD1d-ABd TCRCD1d alone. A.** CD1d alone, ABd TCR or CD1d-ABd TCR-complexes were fractionated by size using fast performance liquid chromatography. **B.** Individual fractions from the co-complex run (red) and CD1d alone run (blue) were normalized based on protein amount and analyzed by HPLC-TOF-MS using external standards to estimate molar ratios. The quantities of each sphingomyelin (SM) and phosphatidylcholine (PC) were separately reported based on the nominal mass (703-835) and overall alkyl chain length (34-44) and unsaturation (1-2) as described in Figure 5.

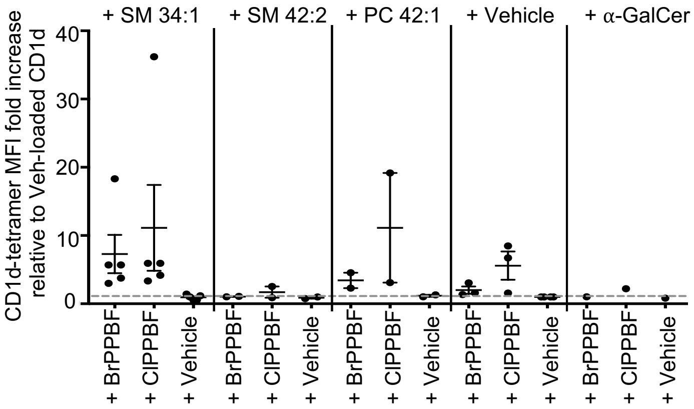


**Supplemental Figure 7. Co-presentation of short chain SM and ClPPBF results in enhanced CD1d-tetramer binding.** VD1G9 TCR+ cell line was stained with CD1d-ClPPBF or BrPPBF pre-loaded with short chain 34:1 SM or long chain 42:2 SM or 42:1 PC. Graphs show fold increase in CD1d tetramer MFI relative to vehicle-loaded CD1d. Each dot represents one individual experiment.
