## Supplementary material for "Benzofuran sulfonates and small self-lipid antigens activate type II NKT cells via CD1d": Chemistry Experimental

### General

Pyridine was distilled over KOH before use. Dichloromethane was dried over alumina according to the method of Pangborn *et al*.^1^ Reactions were monitored using TLC, performed with silica gel 60 F_254_. Detection was effected by charring in a mixture of 5% sulfuric acid in methanol, 10% phosphomolybdic acid in EtOH, and/or visualizing with UV light. Flash chromatography was performed according to the method of Still et al.^2^ using silica gel 60. [α]_D_ values are given in deg 10^−1^ cm^2^ g^−1^. NMR experiments were conducted on a 400 MHz instrument, with chemical shifts referenced relative to residual protiated solvent and are in ppm. The chemical shift for CDCl_3_ was referenced against residual chloroform at δ 7.26 ppm. Mass spectra were acquired in the ESI-QTOF mode.

**2,2,4,6,7-Pentamethyldihydrobenzofuran (1)**

Isobutylaldehyde (12.1 mL, 132 mmol) and H_2_SO_4_ (250 μL) were added to a stirred solution of 2,3,5-trimethylphenol (15 g,110 mmol) in toluene (25 mL). The reaction mixture was refluxed for 4 h, then was concentrated. Flash chromatography of the residue (EtOAc/pet. spirits) afforded **1** as a white solid (15.9 g, 76%), mp 43–44 °C; ^1^H NMR (400 MHz, CDCl_3_) δ 1.48 (6 H, s), 2.09 (3 H, s), 2.16 (3 H, s), 2.21 (3 H, s), 2.92 (2 H, s), 6.50 (1 H, s); HRMS (ESI^+^) calcd for C_13_H_18_O (M + Na)^+^ *m*/*z* 213.1250. Found 213.1261.

**2,2,4,6,7-Pentamethyldihydrobenzofuran-5-sulfonyl chloride (2)**

Chlorosulfuric acid (1.09 g, 9.31 mmol) was added dropwise to a solution of **1** (0.500 g, 2.63 mmol) in CH_2_Cl_2_ (5 mL) at 0 °C. The reaction was stirred at rt for 1.5 h. The mixture was quenched with ice, then the organic phase was separated and washed successively with 5% Na_2_CO_3_ solution, saturated NaHCO_3_, and water, then dried (MgSO_4_) and concentrated. The crude was recrystallized (pet. spirits) to afford **2** as a grey solid (0.418 g, 55%) which was used without further purification.

**2,2,5,7,8-Pentamethylbenzochromane (3)**

2-Methyl-3-buten-2-ol (348 mg, 4.04 mmol) was added dropwise to a solution of 2,3,5-trimethylphenol (500 mg, 3.67 mmol) in TFA (2 mL). The solution was stirred at rt for 2 h then concentrated. The residue dissolved in Et_2_O and washed with saturated NaHCO_3_, water and brine, dried (MgSO_4_) and concentrated. Flash chromatography of the residue (CH_2_Cl_2_/pet. spirits) afforded **3** as a yellow oil (123 g, 16%). ^1^H NMR (400 MHz, CDCl_3_) δ 1.31 (6 H, s), 1.88 (2 H, t, *J* 6.9 Hz), 2.08 (3 H, s), 2.17 (3 H, s), 2.20 (3 H, s), 2.61 (2 H, t, *J* 6.8 Hz), 6.55 (1 H, s); ^13^C NMR (100 MHz, CDCl_3_) δ 11.5, 19.0, 19.9, 20.6, 27.0, 32.9, 73.2, 116.7, 122.1, 122.4, 133.5, 134.8, 151.8; HRMS (ESI^+^) calcd for C_14_H_20_O (M + Na)^+^ *m*/*z* 227.1406. Found 227.1405.

**2,2,5,7,8-Pentamethylbenzochromane-6-sulfonyl chloride (4)**

Chlorosulfuric acid (150 μL, 2.08 mmol) was added dropwise to a solution of **3** (120 mg, 0.587 mmol) in CH_2_Cl_2_ (2 mL) at 0 °C. The reaction was stirred at rt for 1.5 h. The mixture was quenched with ice, then the organic phase was separated and washed with 5% Na_2_CO_3_ solution, saturated NaHCO_3_, and water, then dried (MgSO_4_) and concentrated to afford **4** as a yellow solid (50 mg, 28%) which was used without further purification.

**General Procedure for the synthesis of benzofuran and benzochroman sulfonates**

**and sulfonamides:**

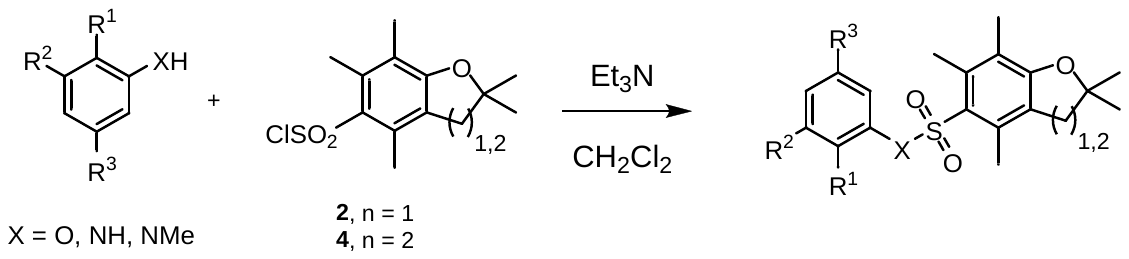

A phenol or aniline (0.58 mmol) was added to a solution of sulfonyl chloride (**2** or **4**, 0.69 mmol) and Et_3_N (100 μL) in CH_2_Cl_2_ (5 mL) at 0 °C. The reaction was stirred for 24 h and then concentrated. The residue was purified by flash chromatography (ethyl ether/pet. sprits) to afford the sulfonate or sulfonamide.

**Phenyl 2,2,4,6,7-pentamethyldihydrobenzofuran-5-sulfonate (5)**

Phenol (55 mg, 0.58 mmol) was reacted with **2** (200 mg, 0.69 mmol) according to the General Procedure to afford **5** (105 mg, 55%). ^1^H NMR (400 MHz, CDCl_3_) δ 1.48 (6 H, s), 2.13 (3 H, s), 2.33 (3 H, s), 2.56 (3 H, s), 2.95 (2 H, s), 7.00–7.06 (2 H, m), 7.19–7.25 (1 H, m), 7.28 (2 H, t, *J* 7.8 Hz); ^13^C NMR (100 MHz, CDCl_3_) δ 12.6, 18.0, 19.2, 28.6, 43.0, 87.4, 118.3, 122.5, 125.3, 126.8, 129.6, 135.2, 141.0, 149.8, 160.8; HRMS (ESI^+^) calcd for C_19_H_22_O_4_S (M + Na)^+^ *m*/*z* 369.1131. Found 369.1139.

**3-Methylphenyl 2,2,4,6,7-pentamethyldihydrobenzofuran-5-sulfonate (6)**

m-Cresol (60 μl, 0.58 mmol) was reacted with **2** (200 mg, 0.69 mmol) according to the General Procedure to afford **6** (120 mg, 57%). ^1^H NMR (400 MHz, CDCl_3_) δ 1.49 (6 H, s), 2.14 (3 H, s), 2.30 (3 H, s), 2.35 (3 H, s), 2.57 (3 H, s), 2.96 (2 H, s), 6.75–6.81 (1 H, m), 6.89–6.93 (1 H, m), 7.00–7.06 (1 H, m), 7.14 (1 H, t, *J* 7.9 Hz); ^13^C NMR (100 MHz, CDCl_3_) δ 12.6, 18.1, 19.3, 21.4, 28.6, 43.1, 87.4, 118.3, 119.1, 123.1, 125.3, 125.4, 127.6, 129.3, 135.2, 139.9, 141.0, 149.7, 160.8; HRMS (ESI^+^) calcd for C_20_H_24_O_4_S (M + Na)^+^ *m*/*z* 383.1288. Found 383.1277.

**3-Fluorophenyl 2,2,4,6,7-pentamethyldihydrobenzofuran-5-sulfonate (7)**

3-Fluorophenol (60 μl, 0.58 mmol) was reacted with **2** (200 mg, 0.69 mmol) according to the General Procedure to afford **7** (124 mg, 46%). ^1^H NMR (400 MHz, CDCl_3_) δ 1.49 (6 H, s), 2.14 (3 H, s), 2.35 (3 H, s), 2.56 (3 H, s), 2.96 (2 H, s), 6.78 (1 H, dt, *J* 9.4, 2.4 Hz), 6.86 (1 H, dd, *J* 8.3, 2.2 Hz), 6.95 (1 H, td, *J* 8.4, 2.5 Hz), 7.22–7.31 (1 H, m); ^13^C NMR (100 MHz, CDCl_3_) δ 12.6, 18.1, 19.3, 28.6, 43.0, 87.6, 110.4, 110.6, 113.9, 114.1, 118.3, 118.6, 124.8, 125.5, 130.3, 130.4, 135.3, 141.1, 161.1; HRMS (ESI^+^) calcd for C_19_H_21_FO_4_S (M + Na)^+^ *m*/*z* 387.1037. Found 387.1030.

**3-Chlorophenyl 2,2,4,6,7-pentamethyldihydrobenzofuran-5-sulfonate (8)**

3-Chlorophenol (82 μl, 0.58 mmol) was reacted with **2** (200 mg, 0.69 mmol) according to the General Procedure to afford **8** (97 mg, 44%). ^1^H NMR (400 MHz, CDCl_3_) δ 1.49 (6 H, s), 2.14 (3 H, s), 2.35 (3 H, s), 2.56 (3 H, s), 2.97 (2 H, s), 6.92–7.00 (1 H, m), 7.02–7.07 (1 H, m), 7.19–7.24 (2 H, m); ^13^C NMR (100 MHz, CDCl_3_) δ 12.5, 17.9, 19.1, 28.5, 42.9, 87.4, 118.4, 120.7, 122.9, 124.6, 125.3, 130.2, 134.7, 135.1, 150.0, 160.9; HRMS (ESI^+^) calcd for C_19_H_21_ClO_4_S (M + Na)^+^ *m*/*z* 403.0741. Found 403.0753.

**3-Bromophenyl 2,2,4,6,7-pentamethyldihydrobenzofuran-5-sulfonate (9)**

3-Bromophenol (61 μl, 0.58 mmol) was reacted with **2** (200 mg, 0.69 mmol) according to the General Procedure to afford **9** (111 mg, 45%). ^1^H NMR (400 MHz, CDCl_3_) δ 1.49 (6 H, s), 2.14 (3 H, s), 2.35 (3 H, s), 2.56 (3 H, s), 2.97 (2 H, s), 6.96–7.02 (1 H, m), 7.16 (1 H, t, *J* 8.1 Hz), 7.19 (1 H, t, *J* 2.0 Hz), 7.33–7.38 (1 H, m); ^13^C NMR (100 MHz, CDCl_3_) δ 12.6, 18.0, 19.3, 28.6, 42.9, 87.6, 118.5, 121.2, 122.4, 124.6, 125.5, 125.8, 130.0, 130.6, 135.2, 141.1, 150.1, 161.1; HRMS (ESI^+^) calcd for C_19_H_21_BrO_4_S (M + Na)^+^ *m*/*z* 447.0236. Found 447.0237.

**3-Isopropylphenyl 2,2,4,6,7-pentamethyldihydrobenzofuran-5-sulfonate (10)**

3-Isopropylphenol (48 μl, 0.35 mmol) was reacted with **2** (200 mg, 0.69 mmol) according to the General Procedure to afford **10** (54 mg, 40%). ^1^H NMR (400 MHz, CDCl_3_) δ 1.12 (6 H, d, *J* 6.9 Hz), 1.48 (6 H, s), 2.13 (3 H, s), 2.31 (3 H, s), 2.55 (3 H, s), 2.81 (1 H, p, *J* 6.9 Hz), 2.94 (2 H, s), 6.72–6.79 (1 H, m), 6.86–6.91 (1 H, m), 7.05–7.10 (1 H, m), 7.19 (1 H, t, *J* 7.9 Hz); ^13^C NMR (100 MHz, CDCl_3_) δ 12.6, 18.0, 19.2, 23.8, 28.6, 33.9, 43.1, 87.3, 118.3, 119.9, 120.2, 125.0, 125.22, 125.25, 129.4, 135.3, 141.1, 149.9, 150.7, 160.8; HRMS (ESI^+^) calcd for C_22_H_28_O_4_S (M + NH_4_)^+^ *m*/*z* 406.2047. Found 406.2051.

**3-*tert*-Butylphenyl 2,2,4,6,7-pentamethyldihydrobenzofuran-5-sulfonate (11)**

3-*tert*-Butylphenol (48 μl, 0.35 mmol) was reacted with **2** (200 mg, 0.69 mmol) according to the General Procedure to afford **11** (65 mg, 46%). ^1^H NMR (400 MHz, CDCl_3_) δ 1.17 (9 H, s), 1.48 (6 H, s), 2.13 (3 H, s), 2.29 (3 H, s), 2.54 (3 H, s), 2.94 (2 H, s), 6.78–6.81 (1 H, m), 6.89–6.93 (1 H, m), 7.19–7.24 (2 H, m); ^13^C NMR (100 MHz, CDCl_3_) δ 12.6, 18.0, 19.2, 28.6, 31.1, 34.8, 43.1, 87.3, 118.3, 119.4, 119.7, 123.6, 125.2, 125.2, 129.1, 135.3, 141.1, 149.8, 153.1, 160.8; HRMS (ESI^+^) calcd for C_23_H_30_O_4_S (M + Na)^+^ *m*/*z* 425.1757. Found 425.1745.

**3-Phenylphenyl 2,2,4,6,7-pentamethyldihydrobenzofuran-5-sulfonate (12)**

3-Hydroxybiphenol (99 mg, 0.58 mmol) was reacted with **2** (200 mg, 0.69 mmol) according to the General Procedure to afford **12** (118 mg, 48%). ^1^H NMR (400 MHz, CDCl_3_) δ 1.50 (6 H, s), 2.17 (3 H, s), 2.38 (3 H, s), 2.60 (3 H, s), 2.97 (2 H, s), 7.06–7.09 (1 H, m), 7.14–7.16 (1 H, m), 7.32–7.49 (7 H, m); ^13^C NMR (100 MHz, CDCl_3_) δ 12.5, 18.0, 19.3, 28.6, 43.0, 87.4, 118.4, 121.0, 121.3, 125.1, 125.4, 125.5, 127.0, 127.9, 128.9, 129.9, 135.3, 139.8, 141.1, 142.8, 150.1, 160.9; HRMS (ESI^+^) calcd for C_25_H_26_O_4_S (M + Na)^+^ 445.1444. Found 445.1462.

**3-Trifluoromethylphenyl 2,2,4,6,7-pentamethyldihydrobenzofuran-5-sulfonate (13)**

3-Trifluoromethylphenol (71 μl, 0.58 mmol) was reacted with **2** (200 mg, 0.69 mmol) according to the General Procedure to afford **13** (106 mg, 44%). ^1^H NMR (400 MHz, CDCl_3_) δ 1.48 (6 H, s), 2.14 (3 H, s), 2.33 (3 H, s), 2.56 (3 H, s), 2.96 (2 H, s), 7.17–7.19 (1 H, m), 7.26–7.30 (1 H, m), 7.41–7.51 (2 H, m); ^13^C NMR (100 MHz, CDCl_3_) δ 12.4, 17.9, 19.1, 28.4, 42.8, 87.5, 118.5, 119.5 (q, *J* 3.9 Hz), 121.9, 123.4 (q, *J* 3.8 Hz), 124.3, 124.6, 125.4, 126.00, 126.01, 130.2, 131.9 (q, *J* 33.1 Hz), 135.2, 141.0, 149.7, 161.0; HRMS (ESI^+^) calcd for C_20_H_21_F_3_O_4_S (M + Na)^+^ *m*/*z* 437.1005. Found 437.0092.

**3-Nitrophenyl 2,2,4,6,7-pentamethyldihydrobenzofuran-5-sulfonate (14)**

3-Nitrophenol (81 mg, 0.58 mmol) was reacted with **2** (200 mg, 0.69 mmol) according to the General Procedure to afford **14** (111 mg, 49%). ^1^H NMR (400 MHz, CDCl_3_) δ 1.49 (7 H, s), 2.14 (3 H, s), 2.36 (3 H, s), 2.56 (3 H, s), 2.97 (2 H, s), 7.43–7.48 (1 H, m), 7.51 (1 H, t, *J* 8.2 Hz), 7.81 (1 H, t, *J* 2.2 Hz), 8.08–8.13 (1 H, m); ^13^C NMR (100 MHz, CDCl_3_) δ 12.6, 18.0, 19.3, 28.6, 42.9, 87.8, 117.8, 118.8, 121.6, 124.2, 125.8, 128.9, 130.4, 135.3, 141.2, 148.8, 150.0, 161.4; HRMS (ESI^+^) calcd for C_19_H_21_NO_6_S (M + Na)^+^ *m*/*z* 414.0981. Found 414.0996.

**3-Methoxyphenyl 2,2,4,6,7-pentamethyldihydrobenzofuran-5-sulfonate (15)**

3-Methoxyphenol (72 mg, 0.58 mmol) was reacted with **2** (200 mg, 0.69 mmol) according to the General Procedure to afford **15** (66 mg, 30%). ^1^H NMR (400 MHz, CDCl_3_) δ 1.48 (6 H, s), 2.13 (3 H, s), 2.36 (3 H, s), 2.57 (3 H, s), 2.96 (2 H, s), 3.72 (3 H, s), 6.58–6.63 (2 H, m), 6.74–6.80 (1 H, m), 7.13–7.20 (1 H, m); ^13^C NMR (100 MHz, CDCl_3_) δ 12.4, 17.9, 19.1, 28.5, 42.9, 55.4, 87.3, 108.1, 112.7, 114.2, 118.2, 125.2, 129.8, 135.1, 140.9, 150.6, 160.3, 160.7; HRMS (ESI^+^) calcd for C_20_H_24_O_5_S (M + Na)^+^ *m*/*z* 399.1237. Found 399.1220.

**2,3-Dimethylphenyl 2,2,4,6,7-pentamethyldihydrobenzofuran-5-sulfonate (16)**

2,3-Dimethylphenol (71 mg, 0.58 mmol) was reacted with **2** (200 mg, 0.69 mmol) according to the General Procedure to afford **16** (206 mg, 95%). ^1^H NMR (400 MHz, CDCl_3_) δ 1.50 (6 H, s), 2.15 (3 H, s), 2.23 (3 H, s), 2.28 (3 H, s), 2.36 (3 H, s), 2.57 (3 H, s), 2.99 (2 H, s), 6.59 (1 H, d, *J* 8.1 Hz), 6.93 (1 H, t, *J* 7.8 Hz), 7.02 (1 H, d, *J* 7.4 Hz); ^13^C NMR (100 MHz, CDCl_3_) δ 12.5, 13.1, 17.9, 19.1, 20.2, 28.5, 42.9, 87.2, 118.2, 119.0, 125.2, 125.7, 126.4, 128.0, 130.8, 134.7, 139.0, 140.6, 148.4, 160.6; HRMS (ESI^+^) calcd for C_21_H_26_O_4_S (M + Na)^+^ *m*/*z* 397.1444. Found 397.1462.

**3,5-Dichlorophenyl 2,2,4,6,7-pentamethyldihydrobenzofuran-5-sulfonate (17)**

3,5-Dichlorophenol (95 mg, 0.58 mmol) was reacted with **2** (200 mg, 0.69 mmol) according to the General Procedure to afford **17** (101 mg, 42%). ^1^H NMR (400 MHz, CDCl_3_) δ 1.78 (15 H, s), 2.43 (3 H, s), 2.66 (3 H, s), 2.85 (3 H, s), 3.27 (2 H, s), 7.50–7.52 (2 H, m), 7.55 (1 H, s); ^13^C NMR (100 MHz, CDCl_3_) δ 12.6, 18.0, 19.3, 28.5, 42.9, 87.7, 118.7, 121.4, 124.3, 125.6, 127.2, 135.3, 135.3, 141.1, 150.2, 161.3; HRMS (ESI^+^) calcd for C_19_H_20_Cl_2_O_4_S (M + Na)^+^ *m*/*z* 437.0352. Found 437.0368.

***N,N*-Dimethylaminophenyl 2,2,4,6,7-pentamethyldihydrobenzofuran-5-sulfonate (18)**

*N,N*-Dimethylaminophenol (80 mg, 0.58 mmol) was reacted with **2** (200 mg, 0.69 mmol) according to the General Procedure to afford **18** (113 mg, 50%). ^1^H NMR (400 MHz, CDCl_3_) δ 1.48 (6 H, s), 2.13 (3 H, s), 2.38 (3 H, s), 2.57 (3 H, s), 2.85 (6 H, s), 2.96 (2 H, s), 6.29–6.34 (2 H, m), 6.50–6.56 (1 H, m), 7.05–7.11 (1 H, m); ^13^C NMR (100 MHz, CDCl_3_) δ 12.6, 18.0, 19.3, 28.6, 40.4, 43.0, 87.3, 106.1, 109.5, 110.6, 118.2, 125.2, 125.8, 129.7, 135.1, 140.9, 151.0, 151.6, 160.6; HRMS (ESI^+^) calcd for C_21_H_27_NO_4_S (M + H)^+^ *m*/*z* 390.1734. Found 390.1720.

**3-Methylphenyl 2,2,5,7,8-pentamethylbenzochromane-6-sulfonate (19)**

*m*-Cresol (13 mg, 0.121 mmol) was reacted with **4** (50 mg, 0.181 mmol) according to the General Procedure to afford **19** (31 mg, 68%). ^1^H NMR (400 MHz, CDCl_3_) δ 1.34 (6 H, s), 1.84 (2 H, t, *J* 6.9 Hz), 2.14 (3 H, s), 2.30 (3 H, s), 2.47 (3 H, s), 2.53 (3 H, s), 2.65 (2 H, t, *J* 6.8 Hz), 6.78–6.81 (1 H, m), 6.91–6.93 (1 H, m), 7.02 (1 H, d, *J* 7.6 Hz), 7.14 (1 H, t, *J* 7.9 Hz); ^13^C NMR (100 MHz, CDCl_3_) δ 12.2, 17.3, 18.4, 21.3, 21.4, 26.7, 32.5, 74.3, 118.4, 119.0, 123.0, 124.8, 125.9, 127.4, 129.1, 137.5, 137.6, 139.8, 149.5, 155.6; HRMS (ESI^+^) calcd for C_20_H_24_O_4_S (M + Na)^+^ *m*/*z* 383.1288. Found 383.1302.

**Phenyl 2,2,4,6,7-pentamethyldihydrobenzofuran-5-sulfonamide (20)**

Freshly distilled aniline (76 μl, 0.83 mmol) was reacted with **4** (200 mg, 0.69 mmol) according to the General Procedure to afford **20** (110 mg, 46%). ^1^H NMR (400 MHz, CDCl_3_) δ 1.46 (6 H, s), 2.09 (3 H, s), 2.43 (3 H, s), 2.56 (3 H, s), 2.94 (2 H, s), 6.62 (1 H, s), 6.93–7.00 (2 H, m), 7.04–7.10 (1 H, m), 7.19–7.25 (2 H, m); ^13^C NMR (100 MHz, CDCl_3_) δ 12.7, 17.9, 19.6, 28.7, 43.2, 87.1, 118.2, 121.2, 124.9, 125.4, 129.4, 134.5, 137.1, 139.6, 160.1; HRMS (ESI^+^) calcd for C_19_H_23_NO_3_S (M + H)^+^ *m*/*z* 346.1471. Found 346.1466.

**3-Methylphenyl 2,2,4,6,7-pentamethyldihydrobenzofuran-5-sulfonamide (21)**

Freshly distilled *m*-toluidine (62 μl, 0.58 mmol) was reacted with **4** (200 mg, 0.69 mmol) according to the General Procedure to afford **21** (104 mg, 50%). ^1^H NMR (400 MHz, CDCl_3_) δ 1.46 (6 H, s), 2.09 (3 H, s), 2.25 (3 H, s), 2.44 (3 H, s), 2.56 (3 H, s), 2.95 (2 H, s), 6.42 (1 H, s), 6.69–6.77 (1 H, m), 6.76–6.81 (1 H, m), 6.84–6.92 (1 H, m), 7.09 (1 H, t, *J* 7.8 Hz); ^13^C NMR (100 MHz, CDCl_3_) δ 12.5, 17.7, 19.4, 21.3, 28.5, 43.1, 86.9, 117.8, 121.6, 124.8, 125.6, 129.0, 135.1, 137.6, 139.4, 159.9; HRMS (ESI^+^) calcd for C_20_H_25_NO_3_S (M + H)^+^ *m*/*z* 360.1628. Found 360.1609.

**Phenyl *N*-methyl-2,2,4,6,7-pentamethyldihydrobenzofuran-5-sulfonamide (22)**

Freshly distilled *N*-methylaniline (89 μl, 0.83 mmol) was reacted with **4** (200 mg, 0.69 mmol) according to the General Procedure to afford **22** (152 mg, 61%). ^1^H NMR (400 MHz, CDCl_3_) δ 1.45 (6 H, s), 2.08 (3 H, s), 2.27 (3 H, s), 2.42 (3 H, s), 2.91 (2 H, s), 3.20 (3 H, s), 7.19–7.29 (5 H, m); ^13^C NMR (100 MHz, CDCl_3_) δ 12.6, 17.8, 19.3, 28.6, 37.7, 43.1, 86.9, 118.0, 125.1, 126.6, 127.2, 127.4, 129.0, 135.5, 140.9, 142.1, 160.0; HRMS (ESI^+^) calcd for C_20_H_25_NO_3_S (M + Na)^+^ *m*/*z* 382.1447. Found 382.1461.

**3-Chlorophenyl 2,2,5,7,8-pentamethylbenzochromane-6-sulfonate (23)**

3-Chlorophenol (63 μl, 0.63 mmol) was reacted with **4** (200 mg, 0.69 mmol) according to the General Procedure to afford **22** (99.5 mg, 40%). ^1^H NMR (400 MHz, CDCl_3_) δ 1.34 (6 H, s), 1.85 (2 H, t, *J* 6.9 Hz), 2.14 (3 H, s), 2.47 (3 H, s), 2.53 (3 H, s), 2.66 (2 H, t, *J* 6.9 Hz), 6.95–6.97 (1 H, m), 7.06–7.07 (1 H, m), 7.21–7.23 (2 H, m); ^13^C NMR (100 MHz, CDCl_3_) δ 12.5, 17.6, 18.6, 21.6, 26.9, 32.7, 74.7, 118.9, 120.9, 123.2, 125.3, 125.5, 127.2, 130.5, 134.9, 137.8, 137.9, 150.2, 156.1; HRMS (ESI^+^) calcd for C_20_H_23_ClO_4_S (M + Na)^+^ *m*/*z* 397.1049. Found 397.1047.

**2,2,5,7,8-Pentamethylchroman-6-yl 3-methylbenzenesulfonate (24)**

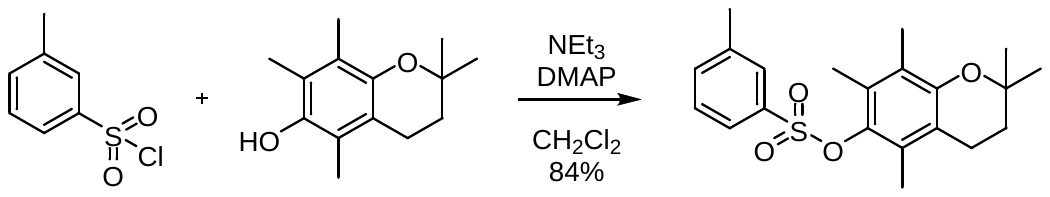

3-Methylbenzene-1-sulfonyl chloride (189 mg, 1.00 mmol, 0.14 mL) was added to a solution of 2,2,5,7,8-pentamethyl-6-chromanol (330 mg, 1.5 mmol), DMAP (12 mg, 0.1 mmol) and Et_3_N (200 μL) in CH_2_Cl_2_ (10 mL) at 0 °C. The reaction was stirred for 24 h and then concentrated. The residue was purified by flash chromatography (ethyl ether/pet. sprits) to afford a white solid **24** (0.316 g, 84%). ^1^H NMR (400 MHz, CDCl_3_) δ 1.30 (6 H, s), 1.79 (2 H, t, *J* 6.9 Hz), 1.95 (3 H, s), 1.97 (3 H, s), 2.04 (3 H, s), 2.44 (3 H, s), 2.56 (2 H, t, *J* 6.8 Hz), 7.41–7.48 (2 H, m), 7.73–7.75 (2 H, m); ^13^C NMR (100 MHz, CDCl_3_) δ 12.0, 13.6, 14.4, 21.1, 21.4, 26.9, 32.7, 73.4, 117.8, 123.7, 125.5, 127.6, 128.6, 129.0, 129.1, 134.7, 137.2, 139.4, 140.3, 150.2; HRMS (ESI^+^) calcd for C_21_H_26_O_4_S (M + H)^+^ *m*/*z* 375.1625. Found 375.1620.

**2,2,5,7,8-Pentamethylbenzochromane (3)**

^1^H NMR

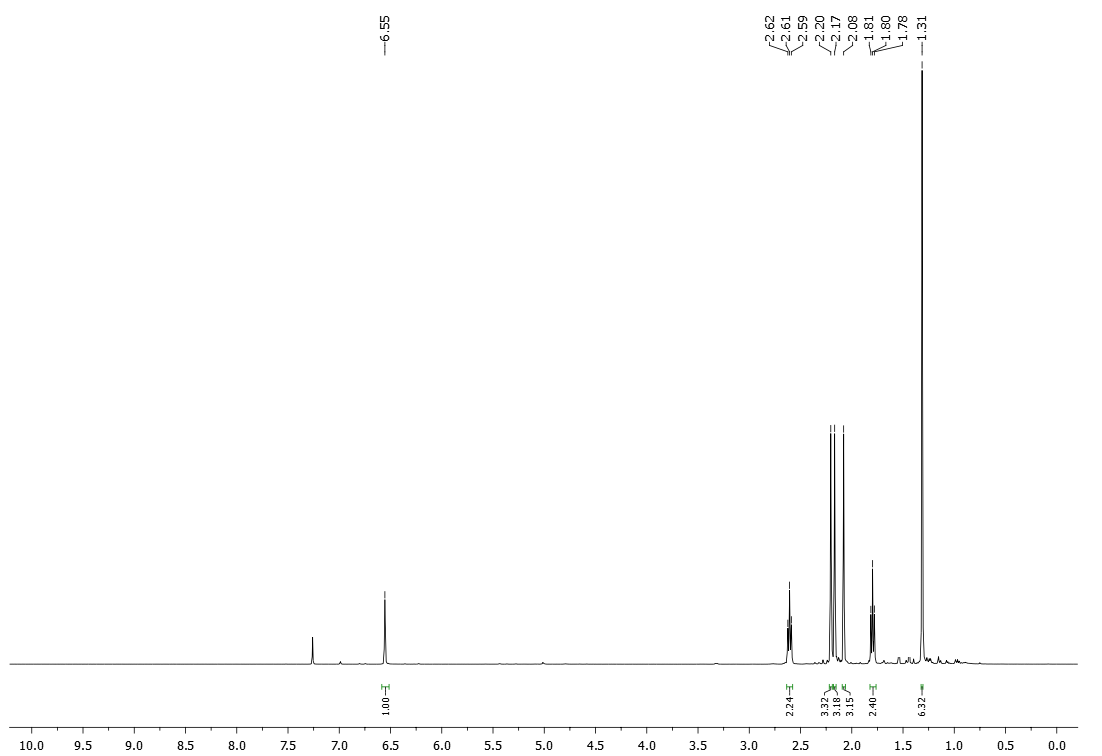

^13^C NMR

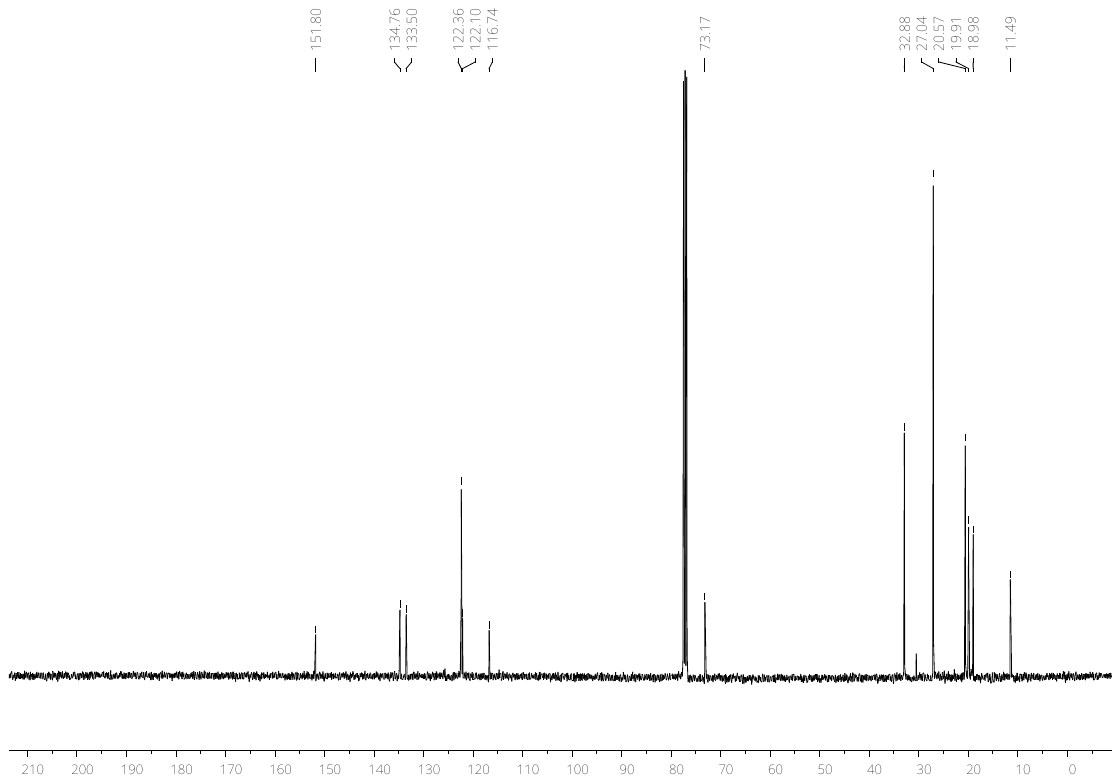

**Phenyl 2,2,4,6,7-pentamethyldihydrobenzofuran-5-sulfonate (5)**

^1^H NMR

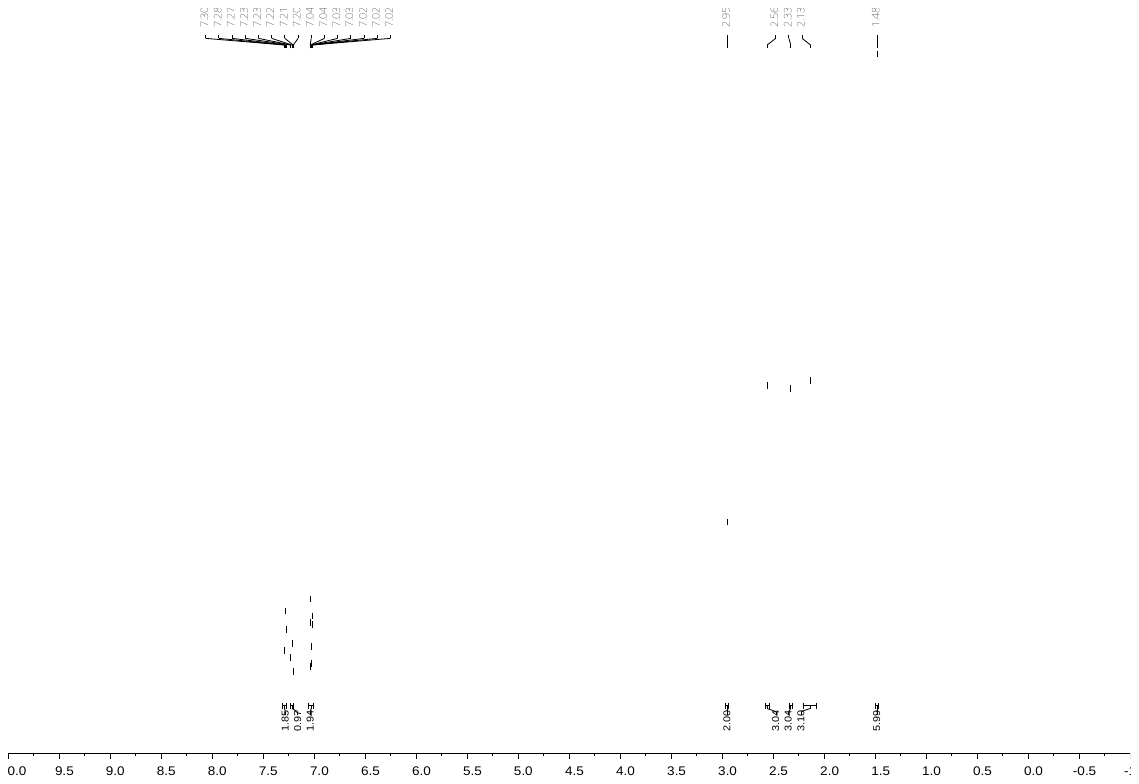

^13^C NMR

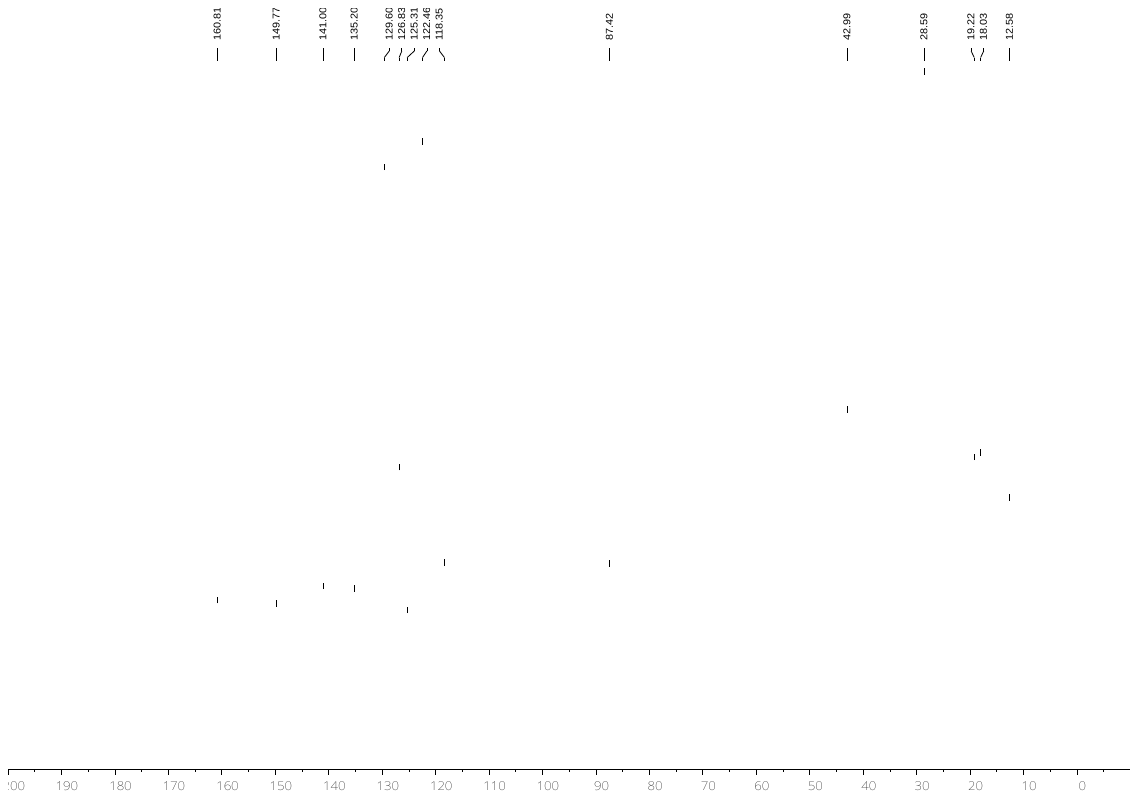

**3-Methylphenyl 2,2,4,6,7-pentamethyldihydrobenzofuran-5-sulfonate (6)**

^1^H NMR

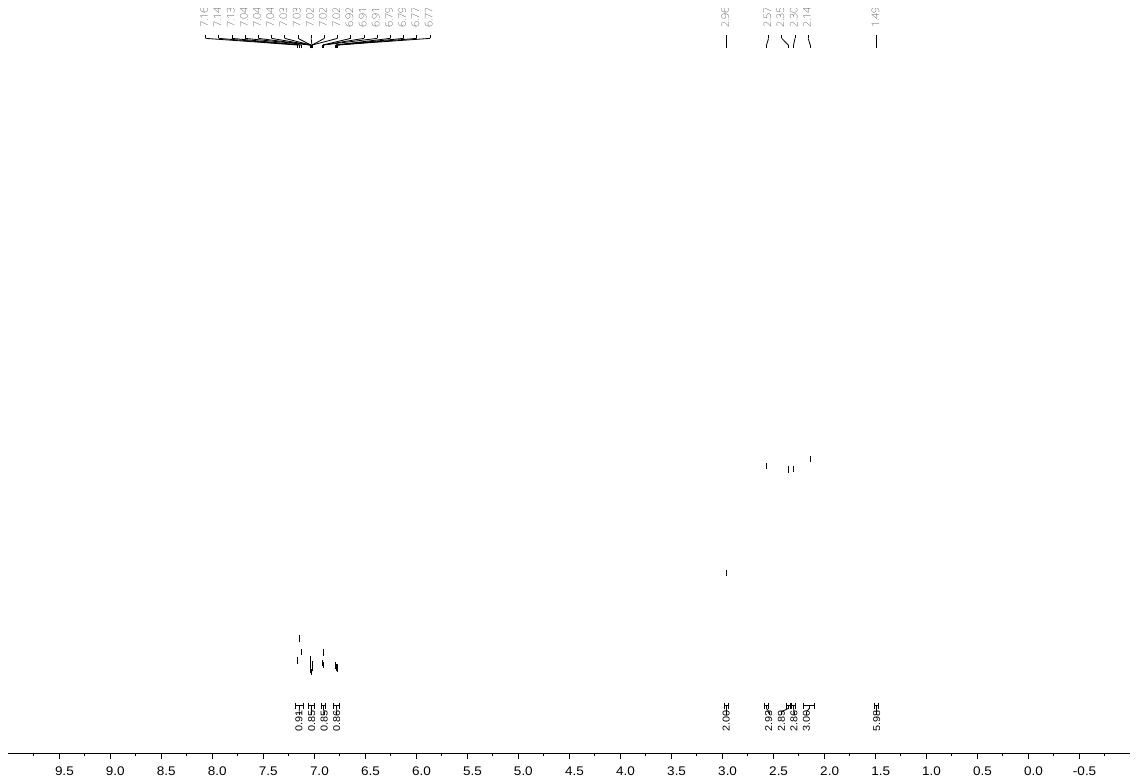

^13^C NMR

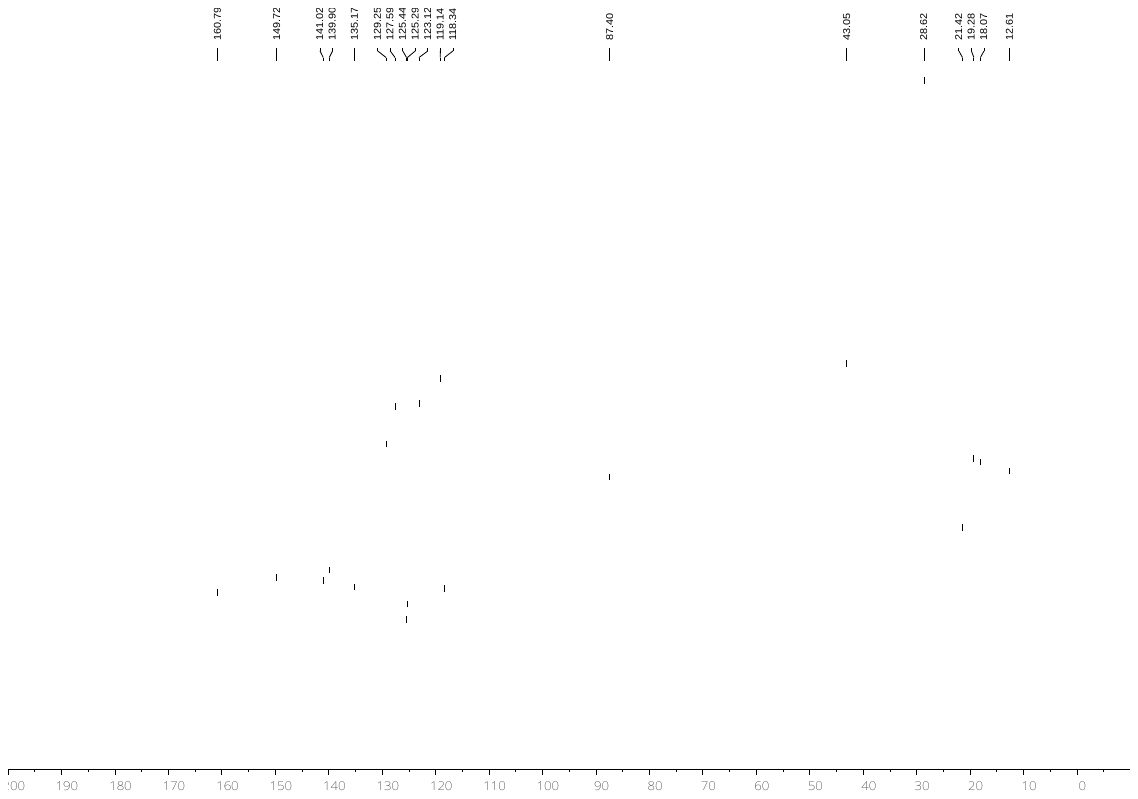

**3-Fluorophenyl 2,2,4,6,7-pentamethyldihydrobenzofuran-5-sulfonate (7)**

^1^H NMR

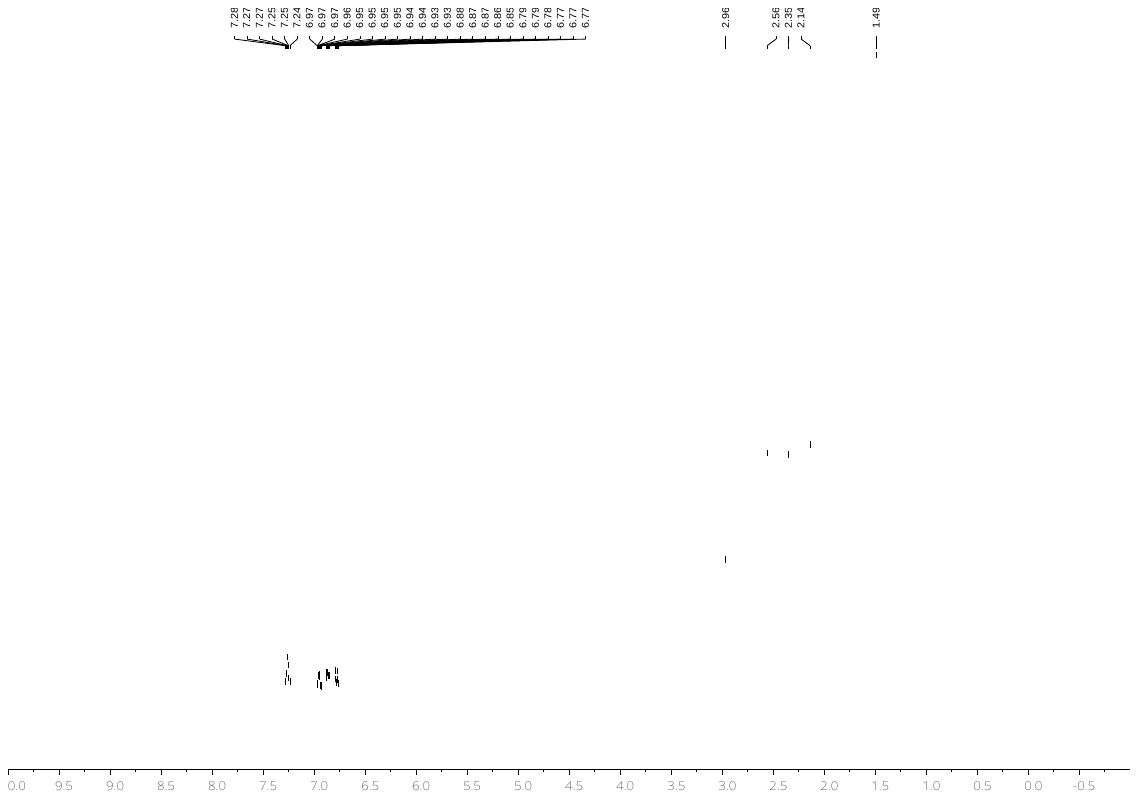

^13^C NMR

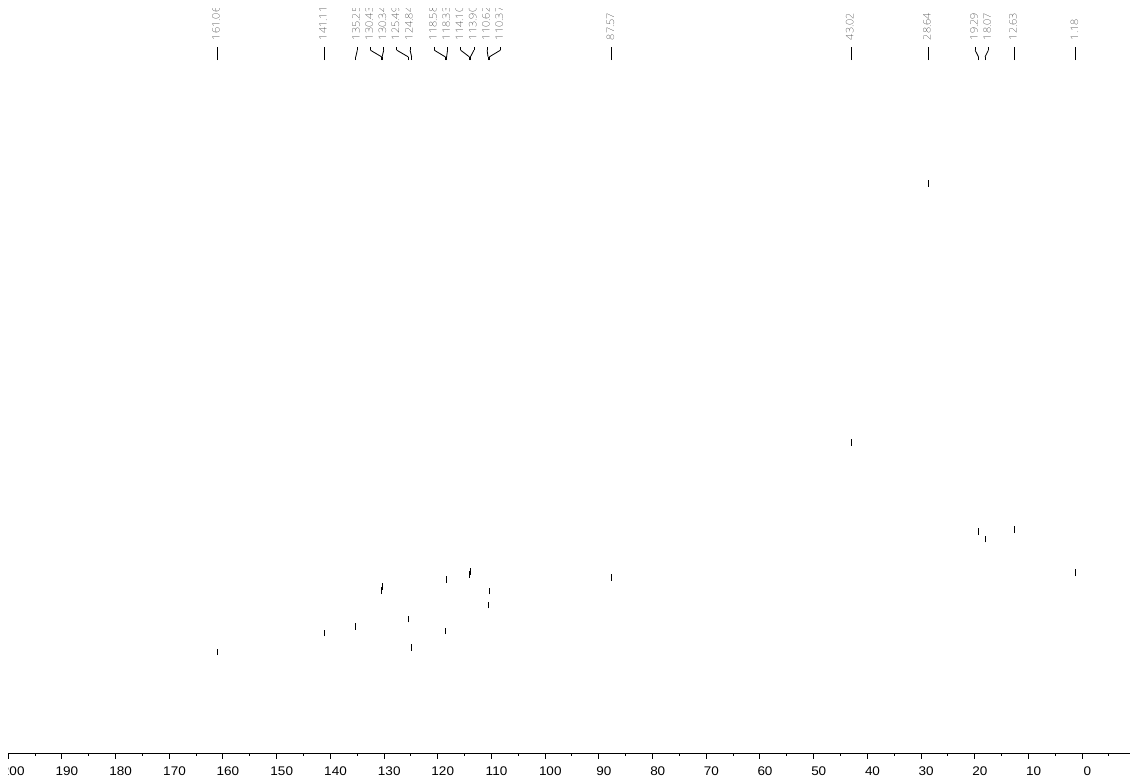

**3-Chlorophenyl 2,2,4,6,7-pentamethyldihydrobenzofuran-5-sulfonate (8)**

^1^H NMR

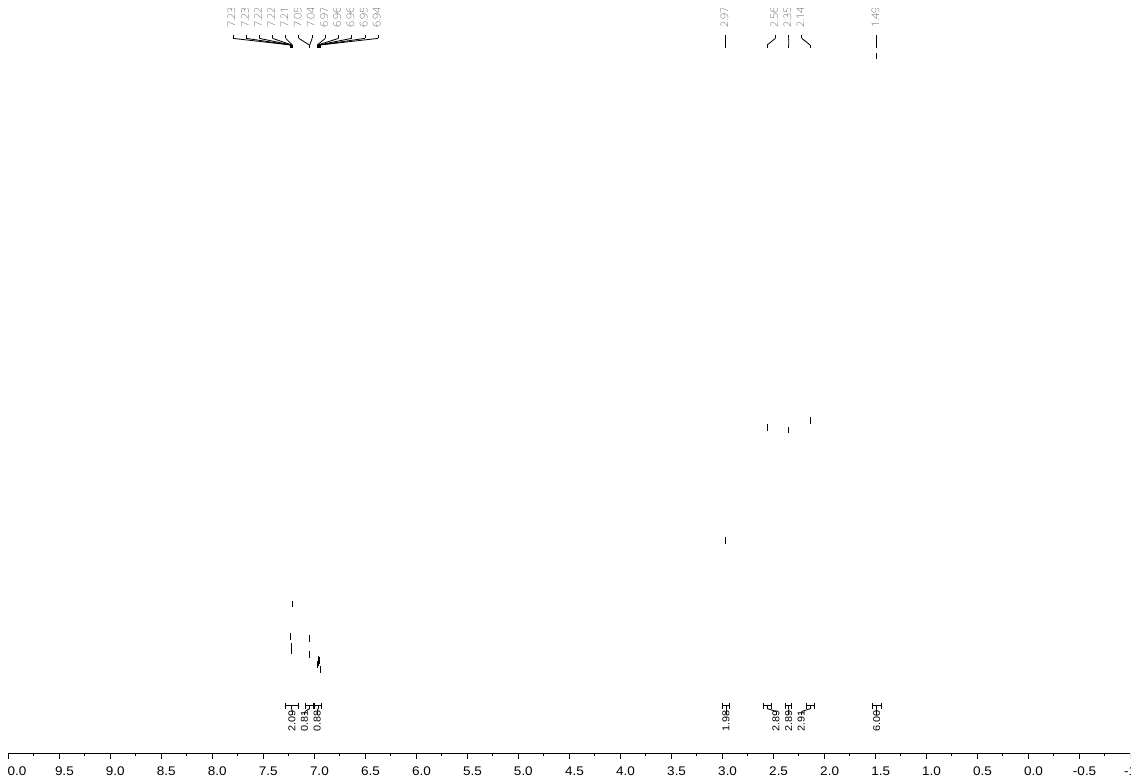

^13^C NMR

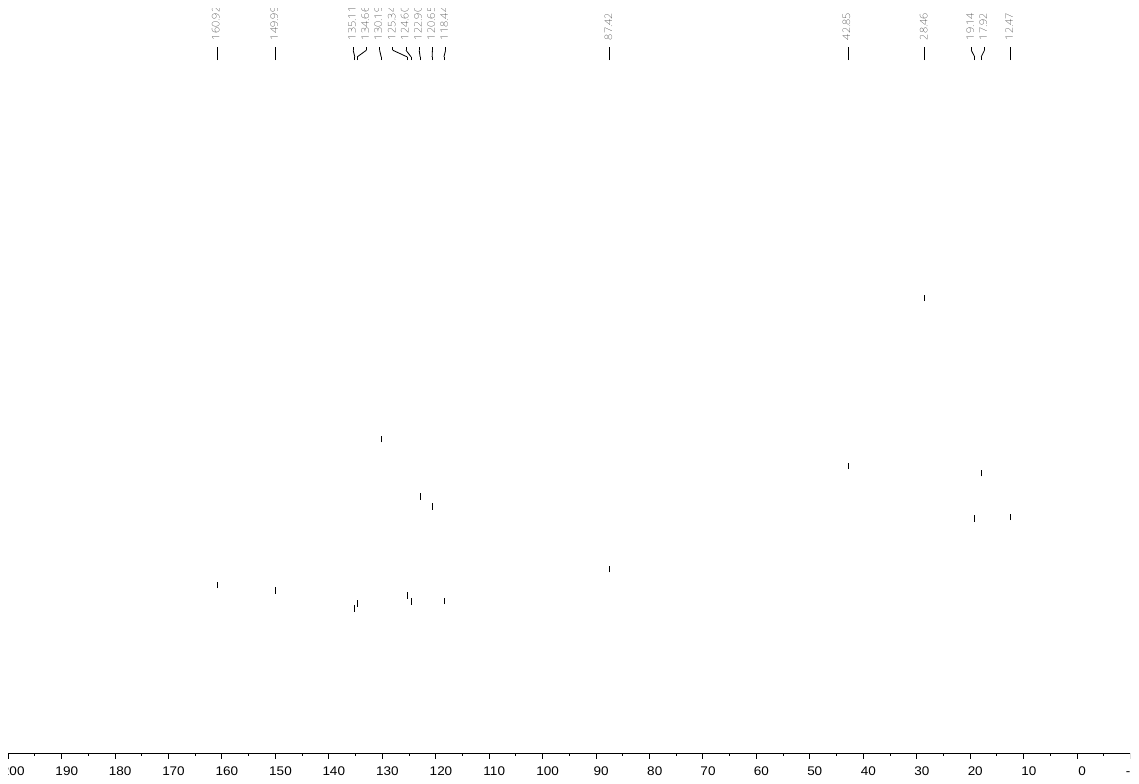

**3-Bromophenyl 2,2,4,6,7-pentamethyldihydrobenzofuran-5-sulfonate (9)**

^1^H NMR

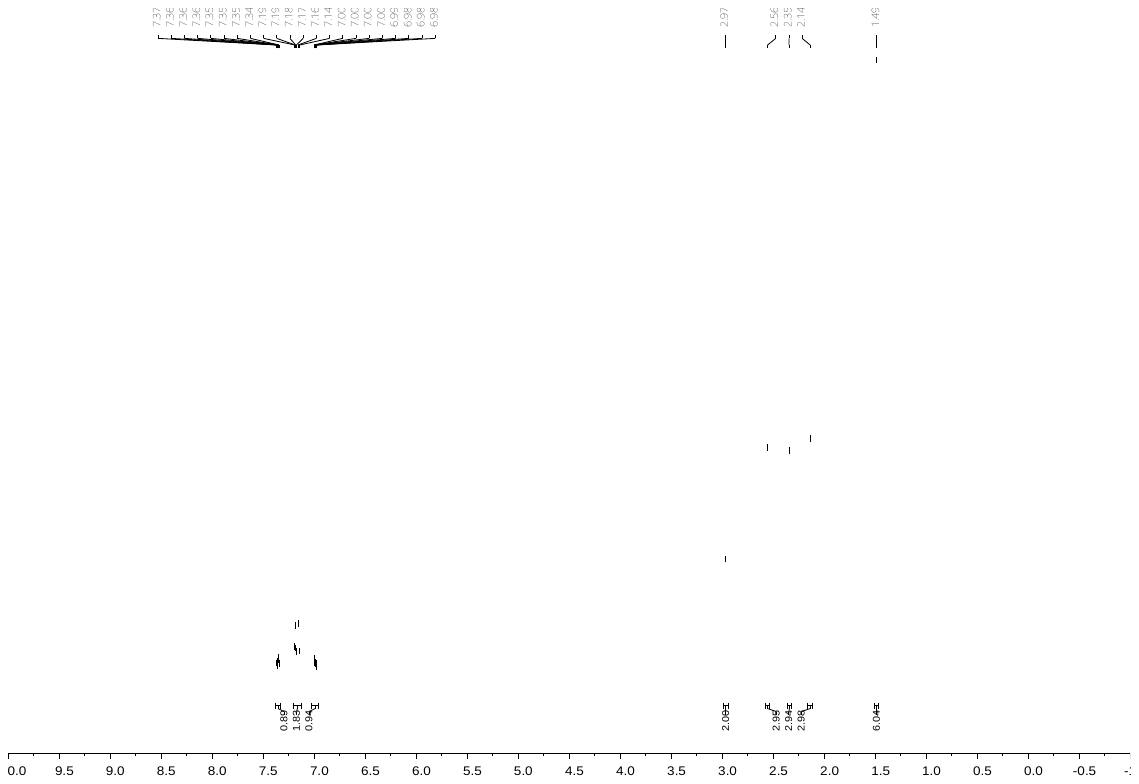

^13^C NMR

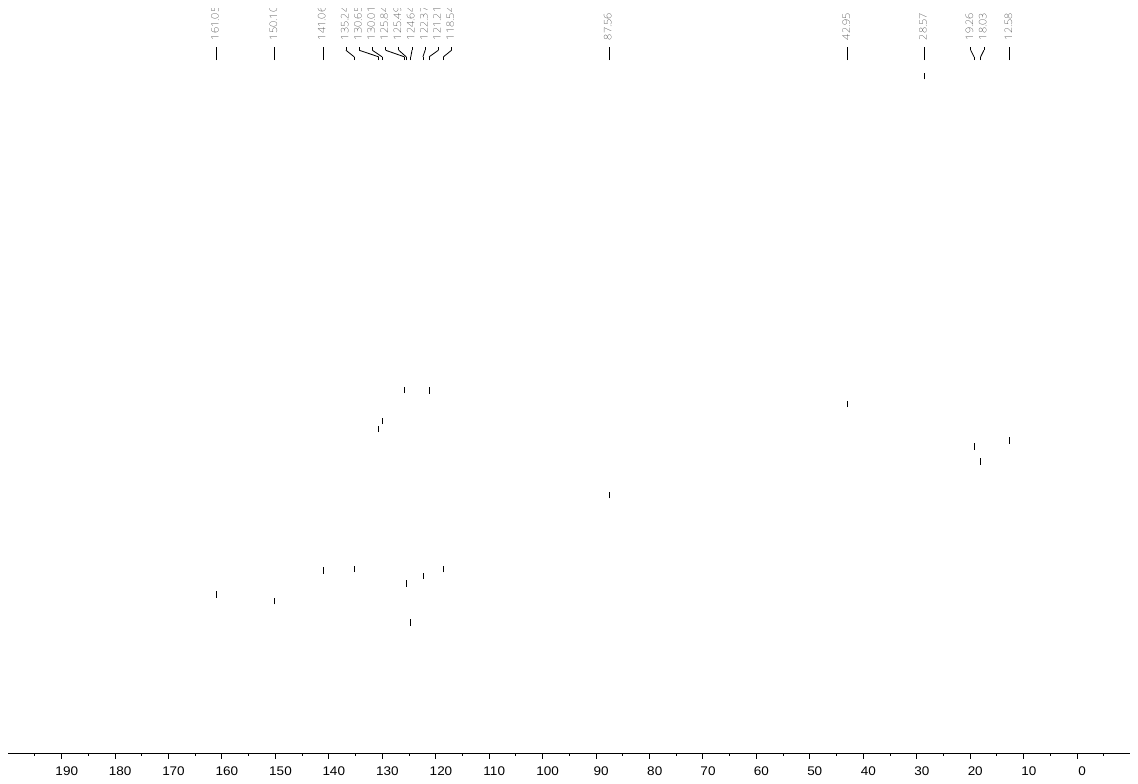

**3-Isopropylphenyl 2,2,4,6,7-pentamethyldihydrobenzofuran-5-sulfonate (10)**

^1^H NMR

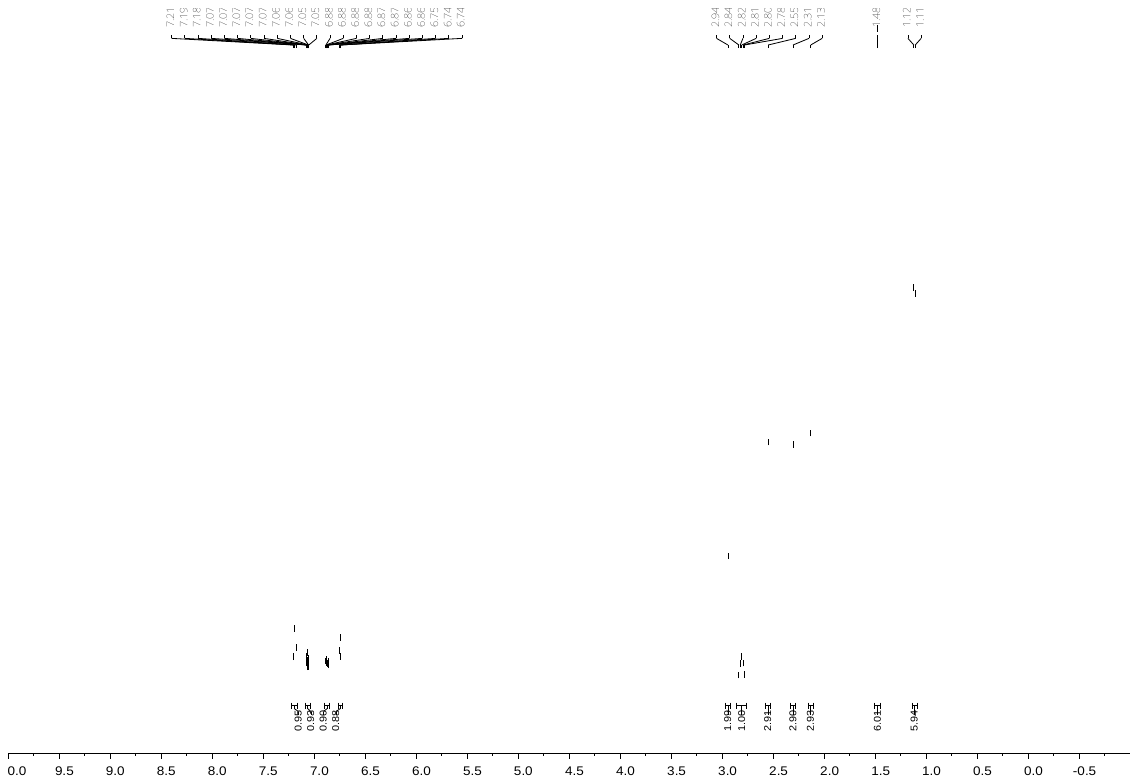

^13^C NMR

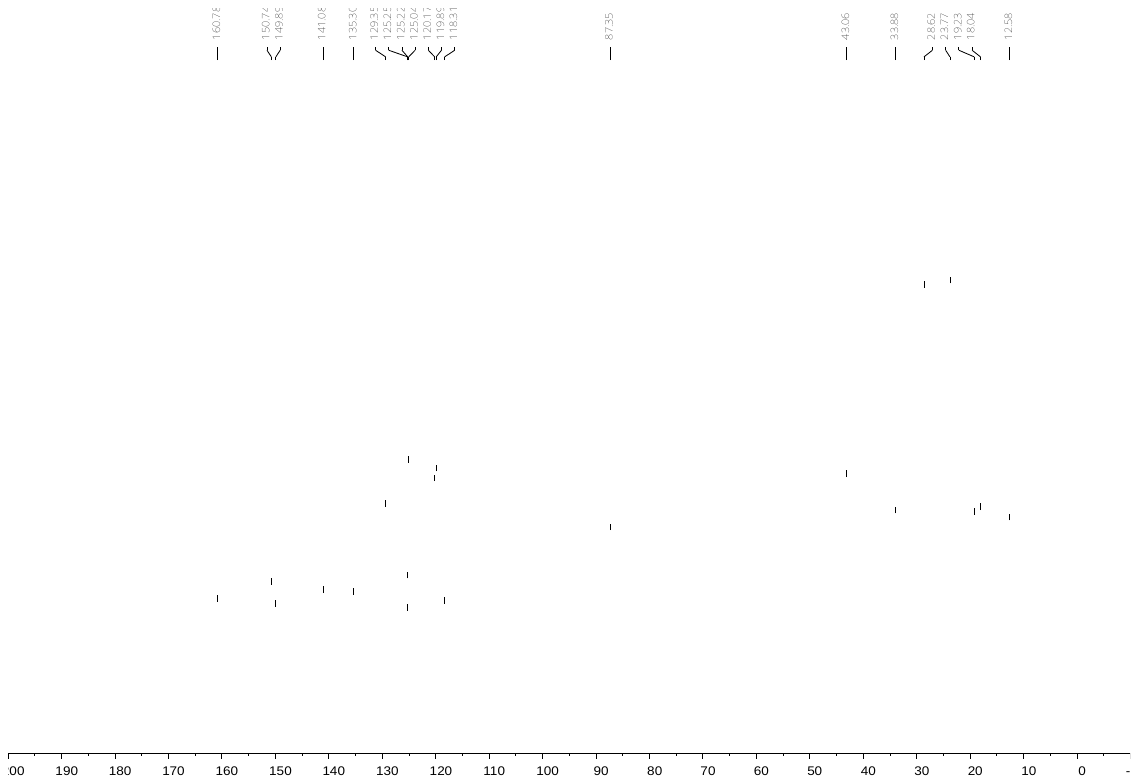

**3-*tert*-Butylphenyl 2,2,4,6,7-pentamethyldihydrobenzofuran-5-sulfonate (11)**

^1^H NMR

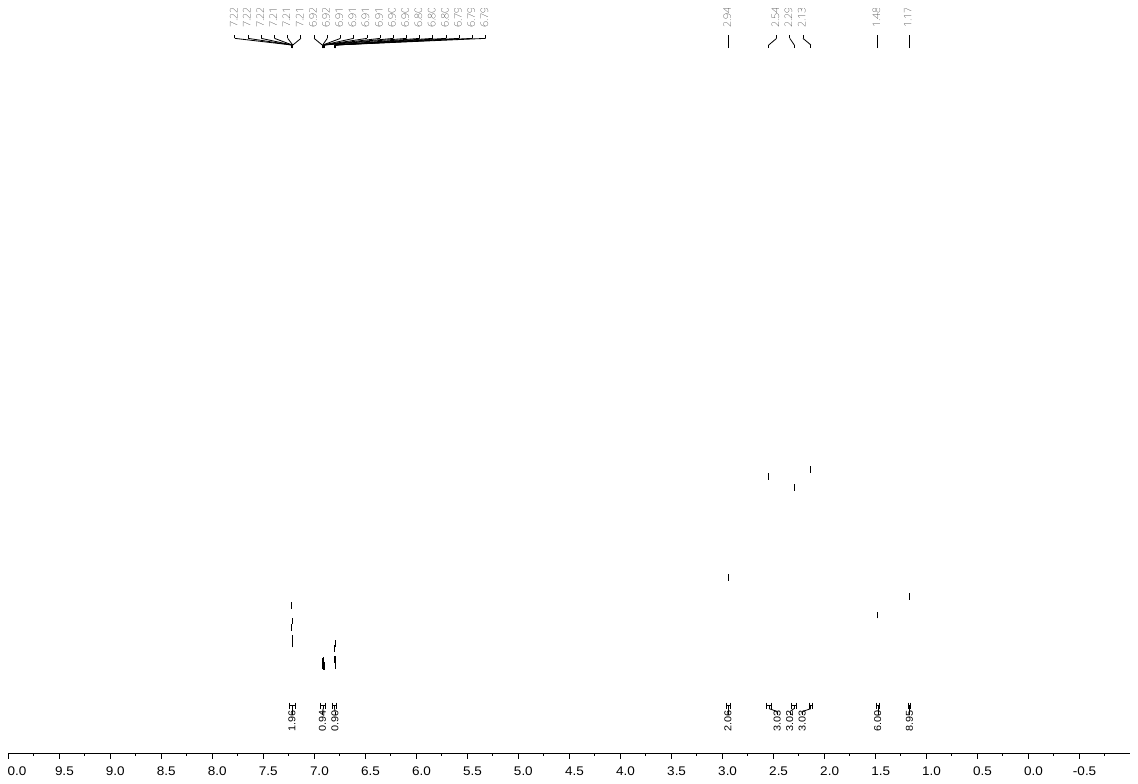

^13^C NMR

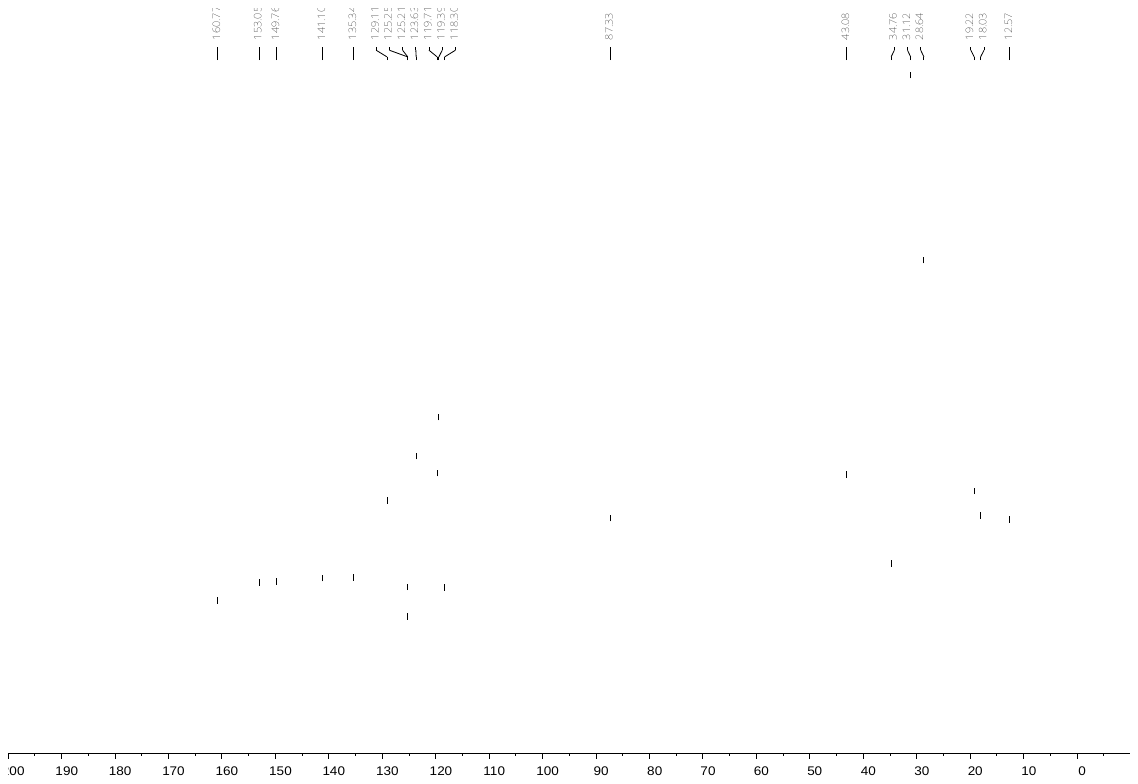

**3-Phenylphenyl 2,2,4,6,7-pentamethyldihydrobenzofuran-5-sulfonate (12)**

^1^H NMR

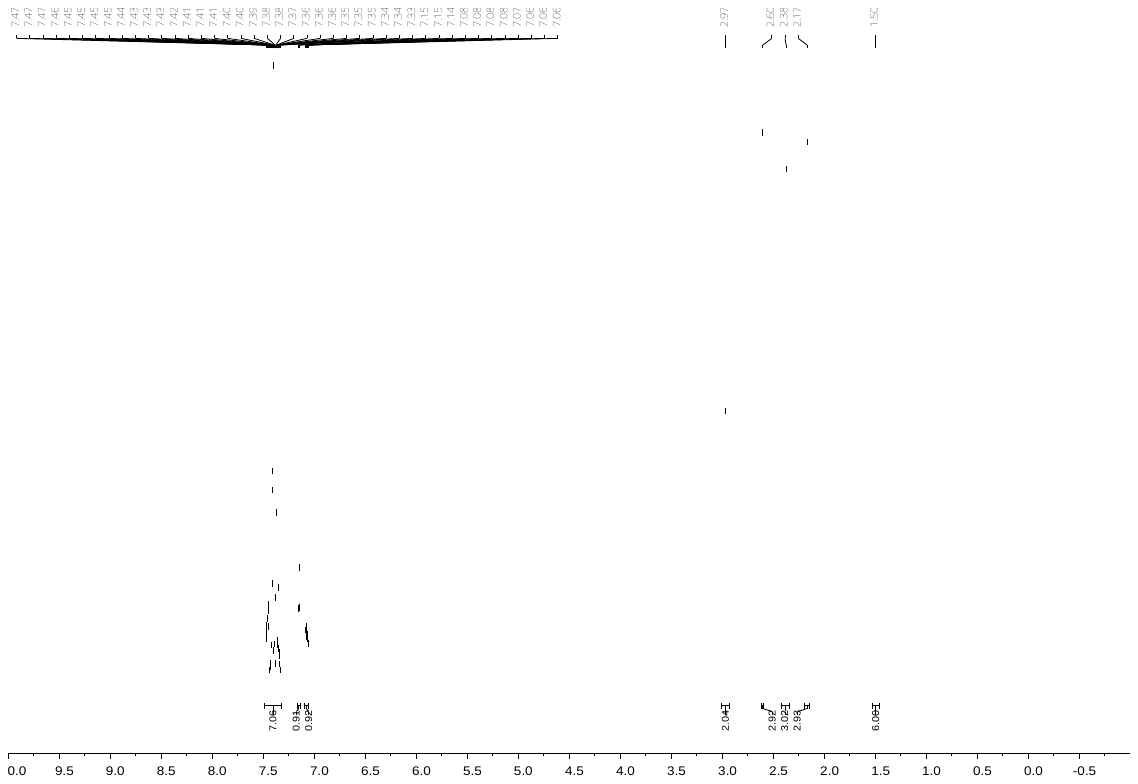

^13^C NMR

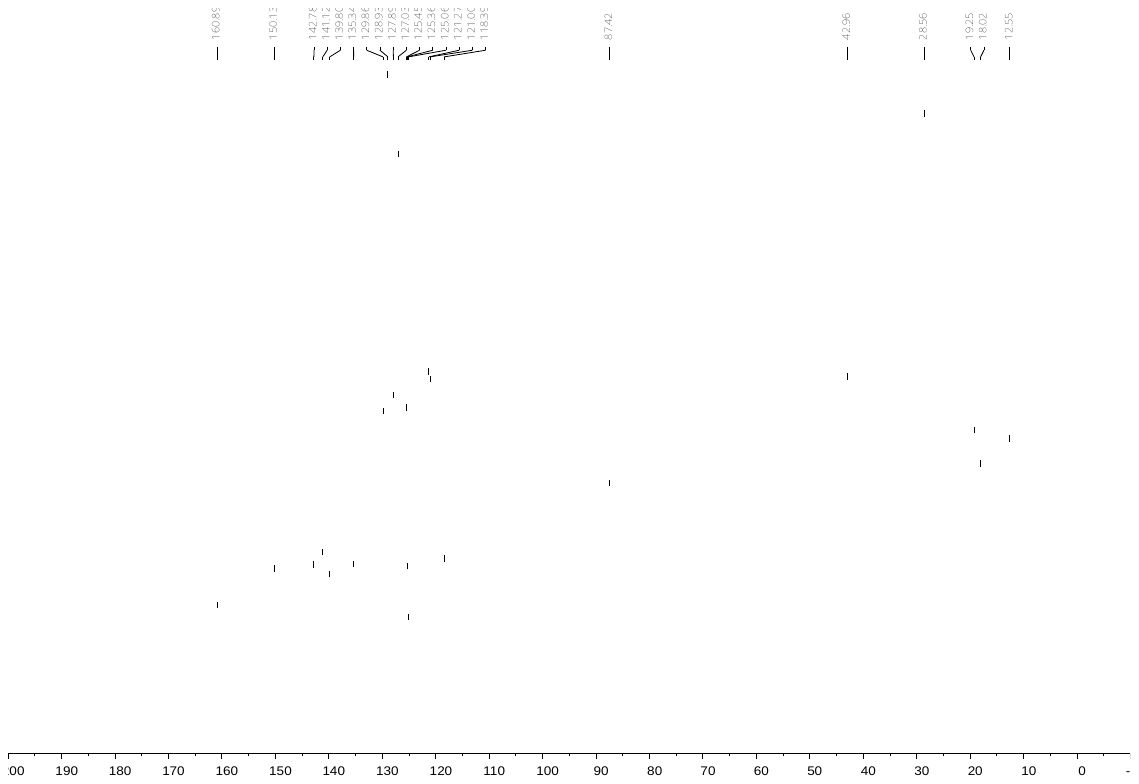

**3-Trifluoromethylphenyl 2,2,4,6,7-pentamethyldihydrobenzofuran-5-sulfonate (13)**

^1^H NMR

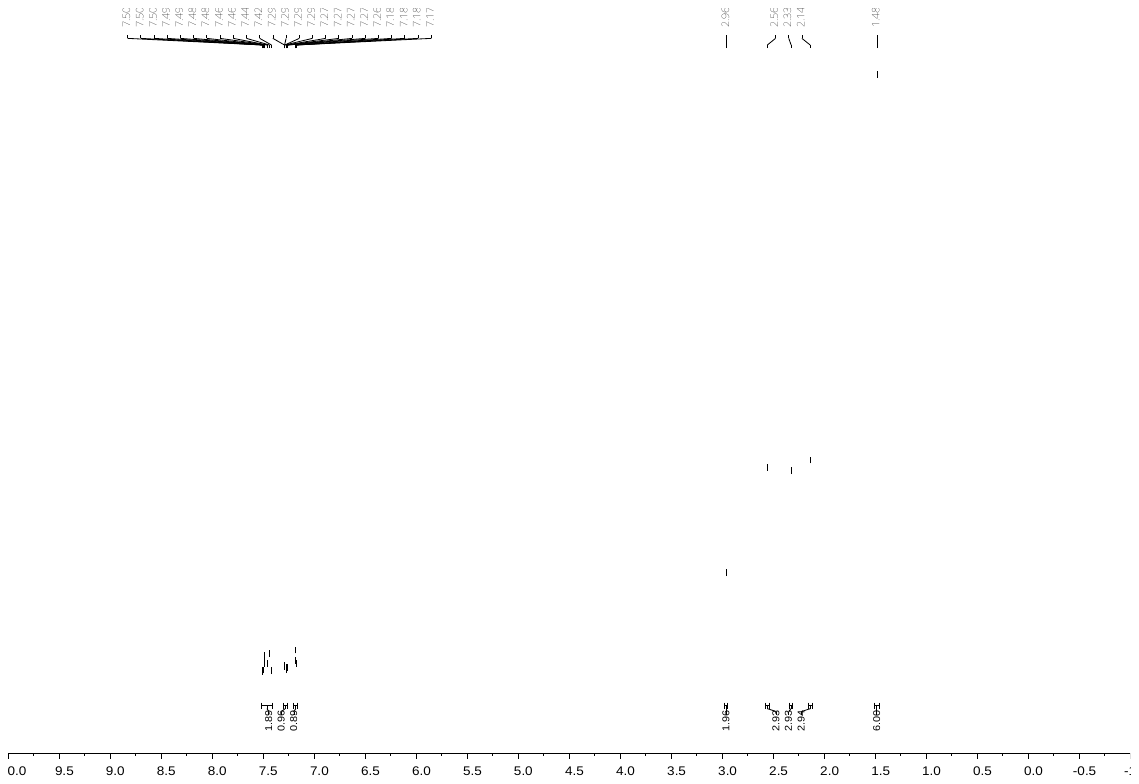

^13^C NMR

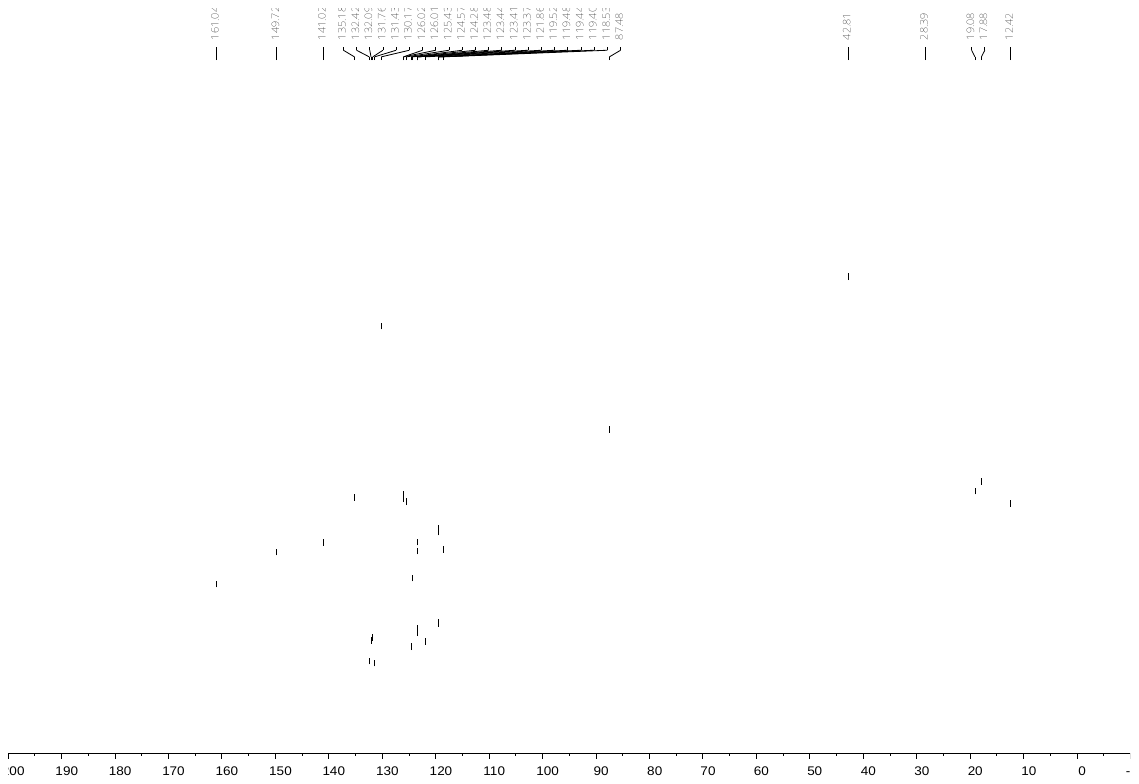

**3-Nitrophenyl 2,2,4,6,7-pentamethyldihydrobenzofuran-5-sulfonate (14)**

^1^H NMR

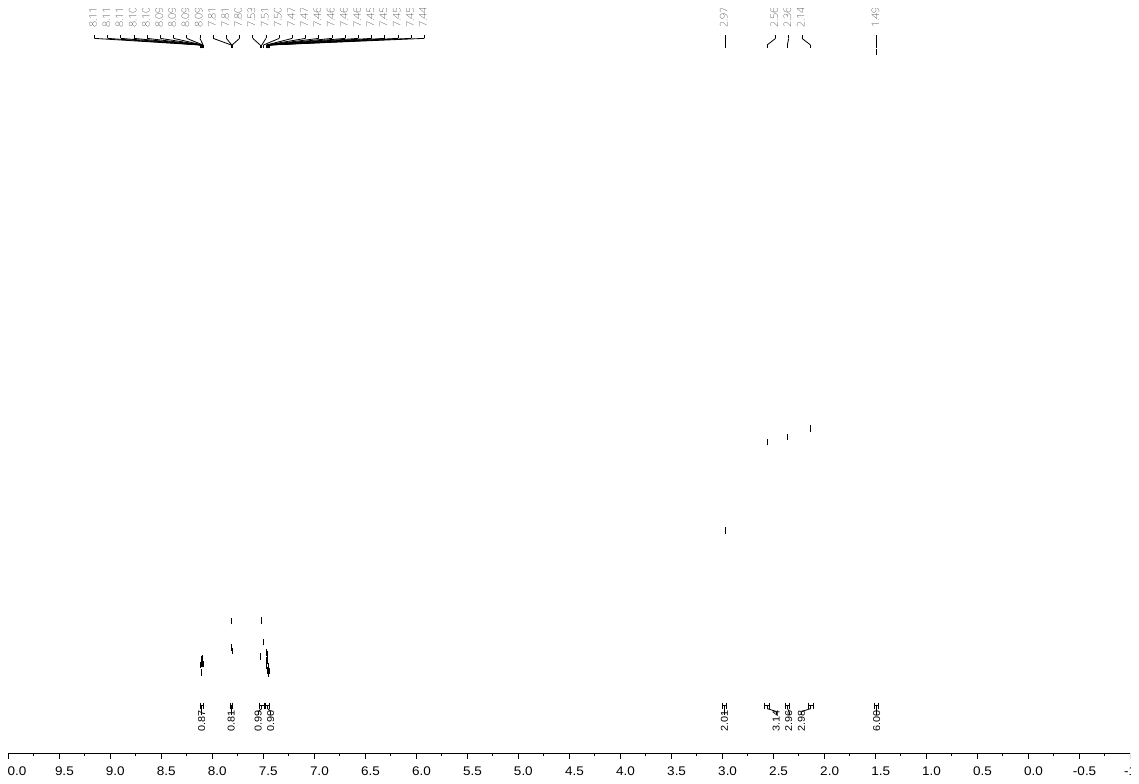

^13^C NMR

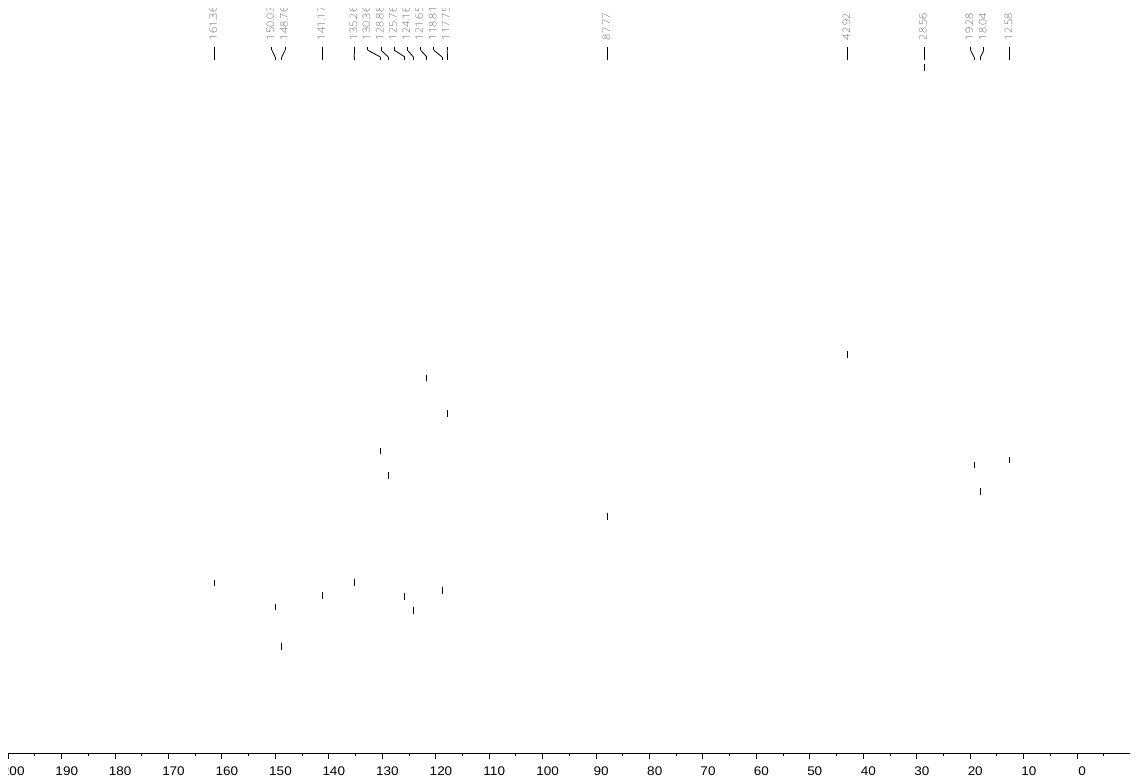

**3-Methoxyphenyl 2,2,4,6,7-pentamethyldihydrobenzofuran-5-sulfonate (15)**

^1^H NMR

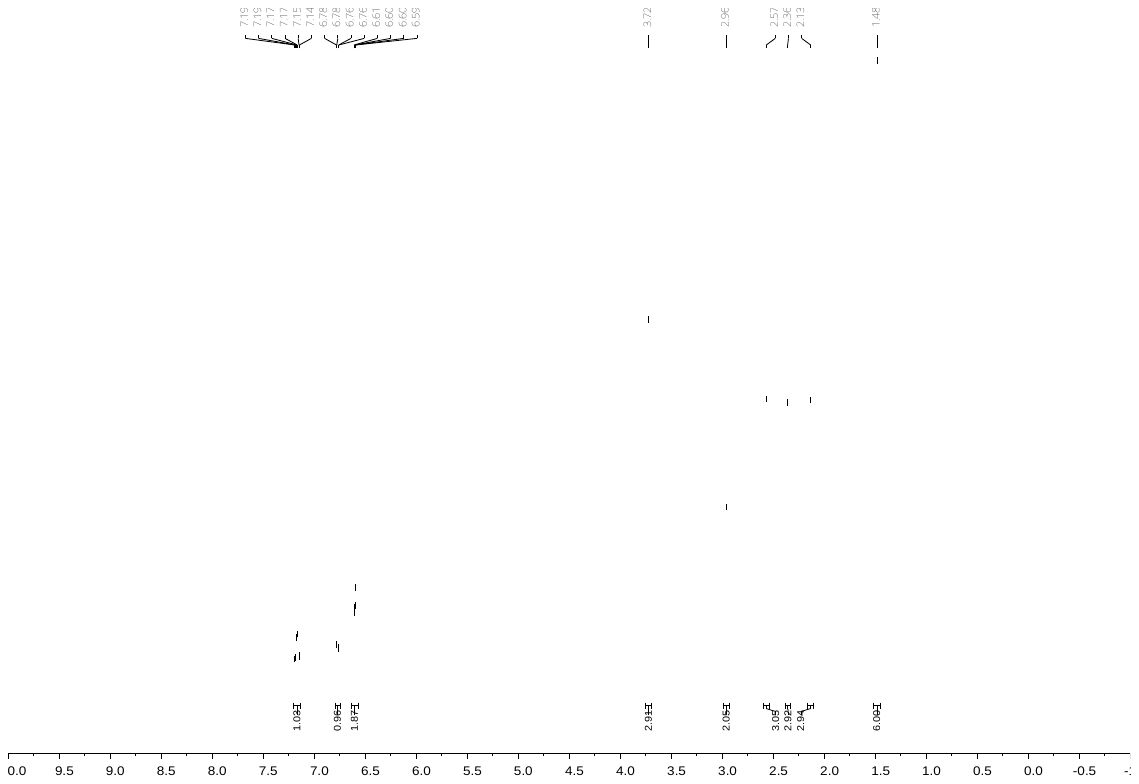

^13^C NMR

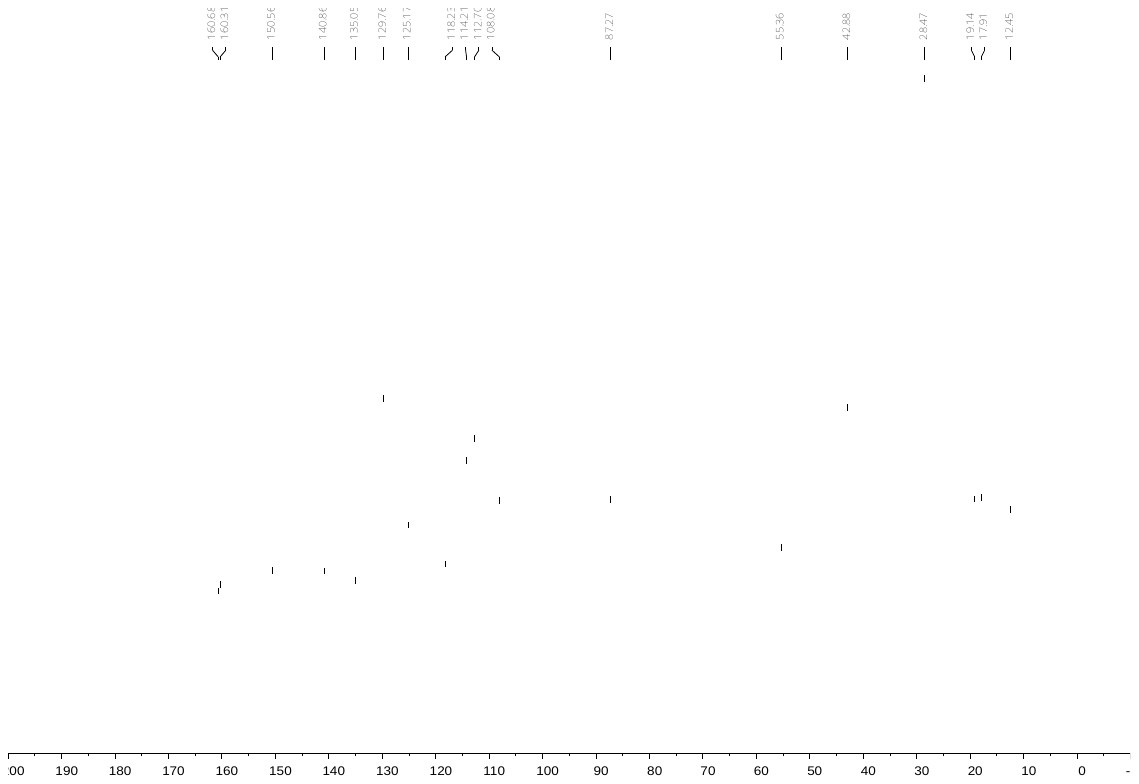

**2,3-Dimethylphenyl 2,2,4,6,7-pentamethyldihydrobenzofuran-5-sulfonate (16)**

^1^H NMR

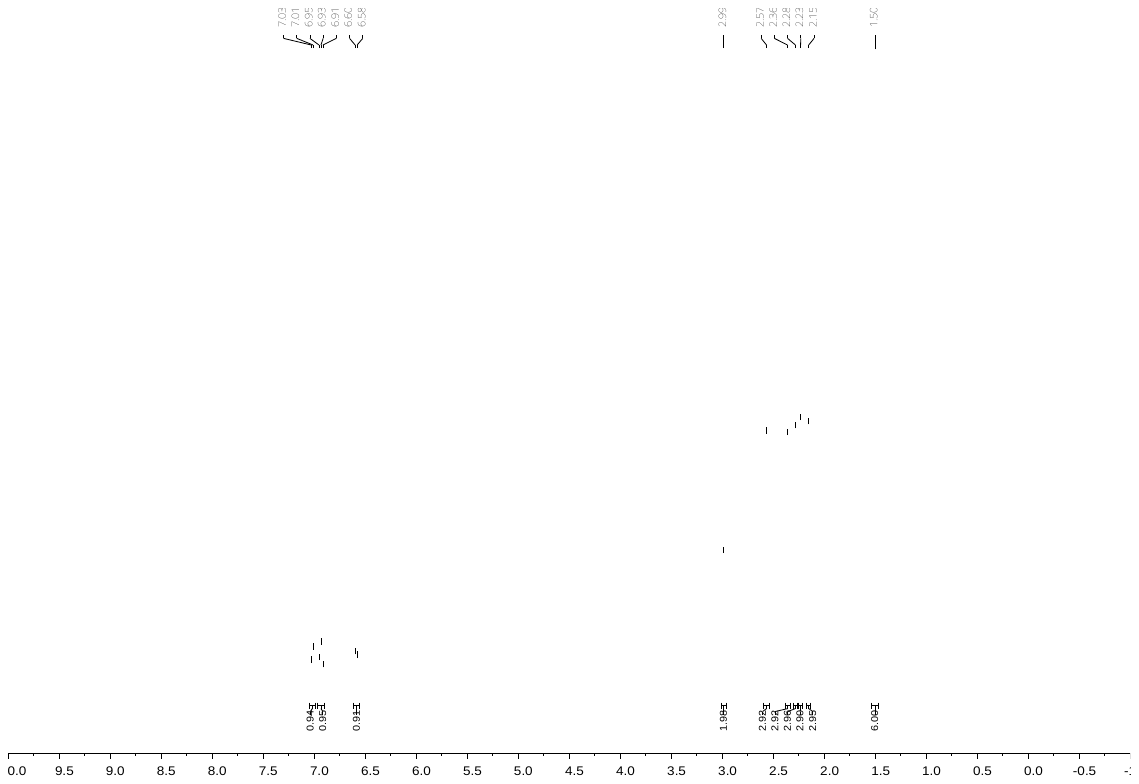

^13^C NMR

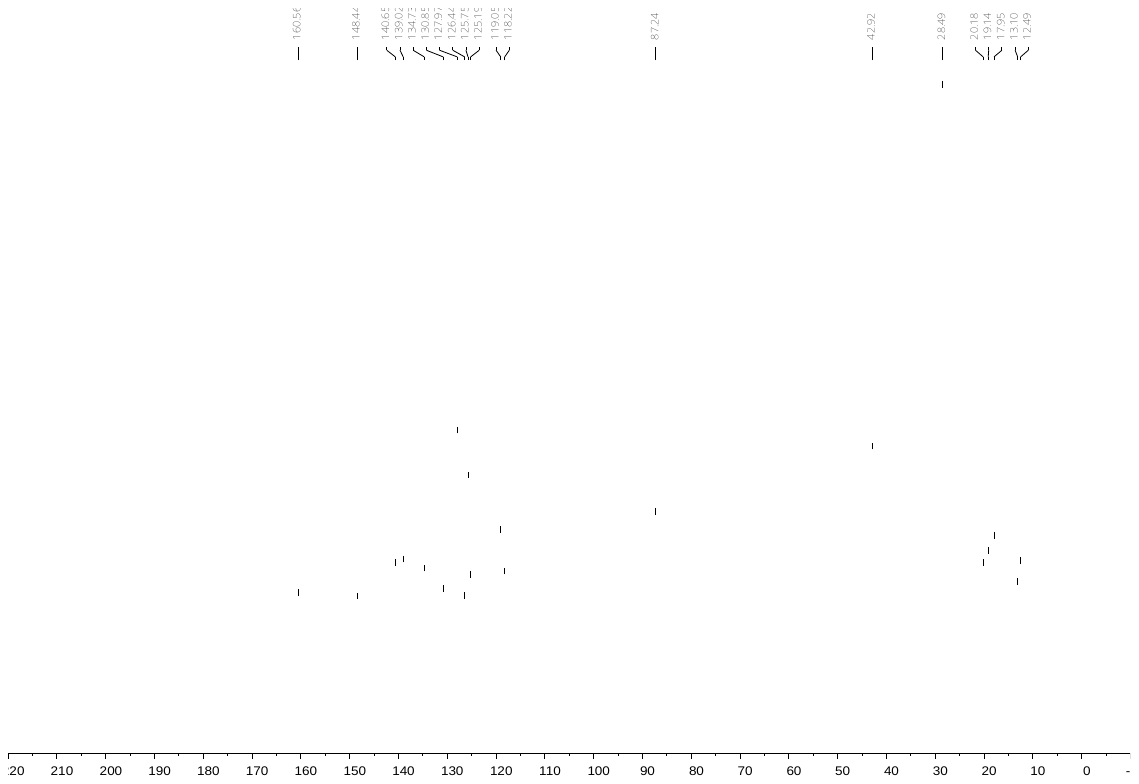

**3,5-Dichlorophenyl 2,2,4,6,7-pentamethyldihydrobenzofuran-5-sulfonate (17)**

^1^H NMR

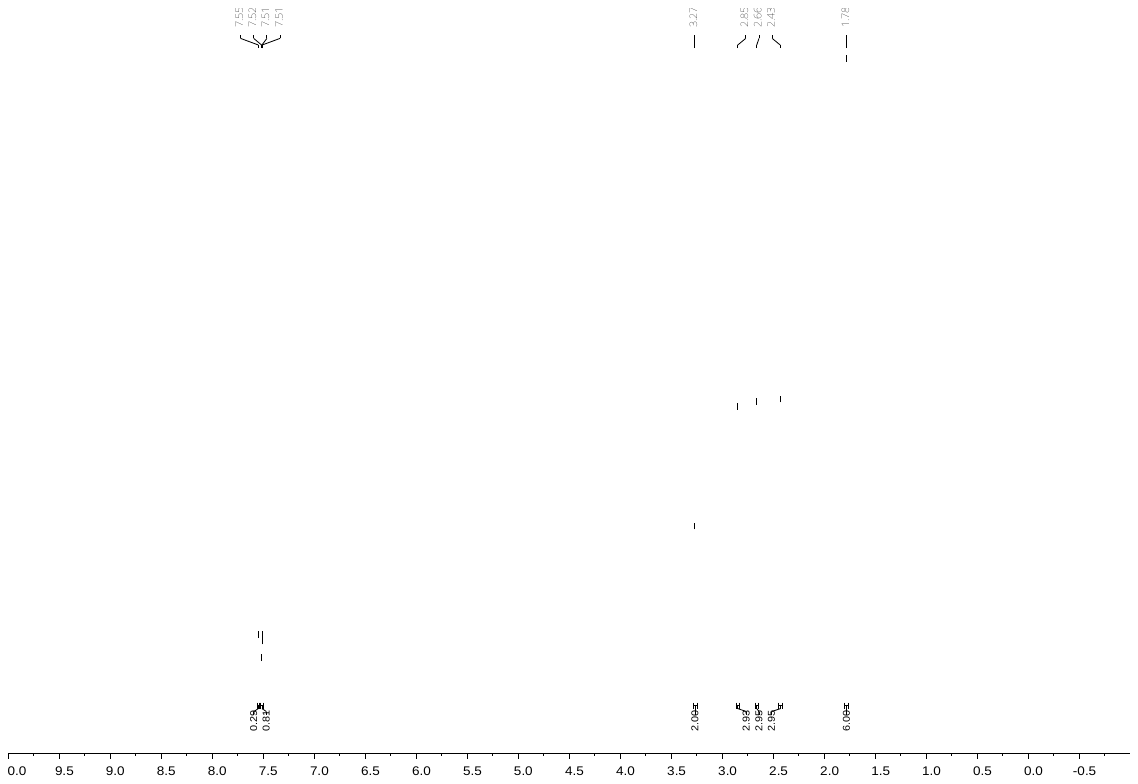

^13^C NMR

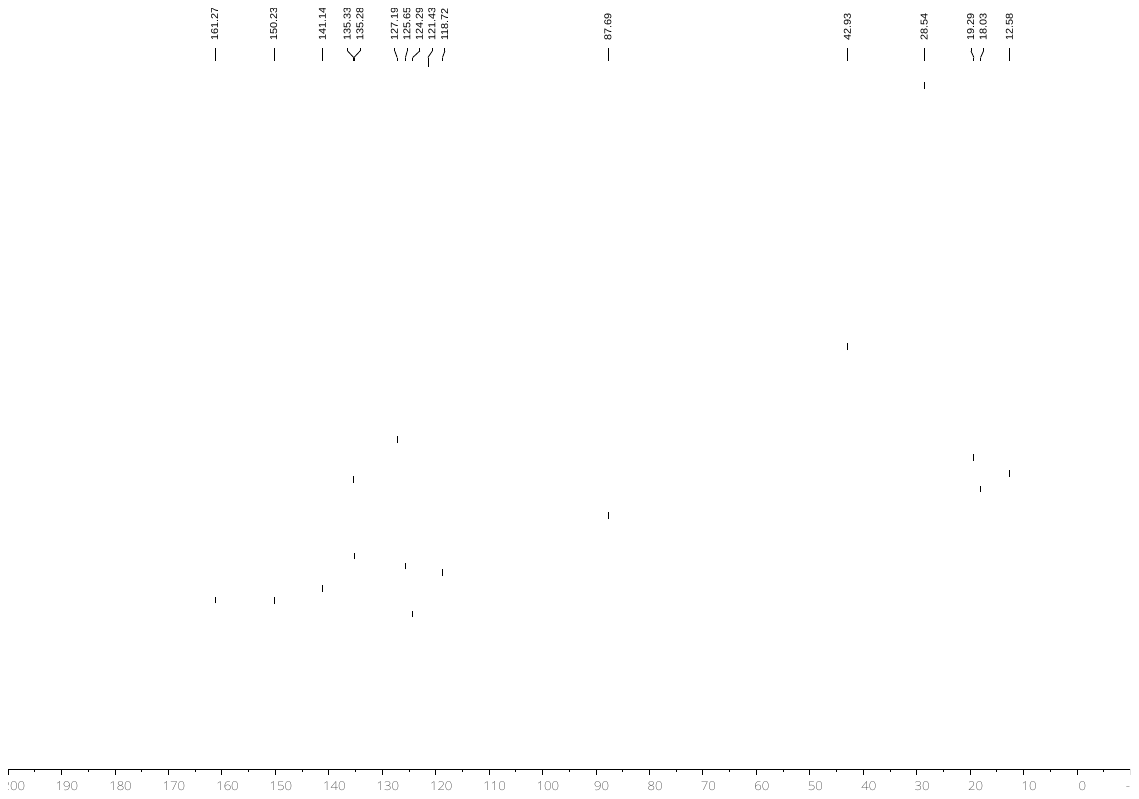

***N*,*N*-Dimethylaminophenyl 2,2,4,6,7-pentamethyldihydrobenzofuran-5-sulfonate (18)**

^1^H NMR

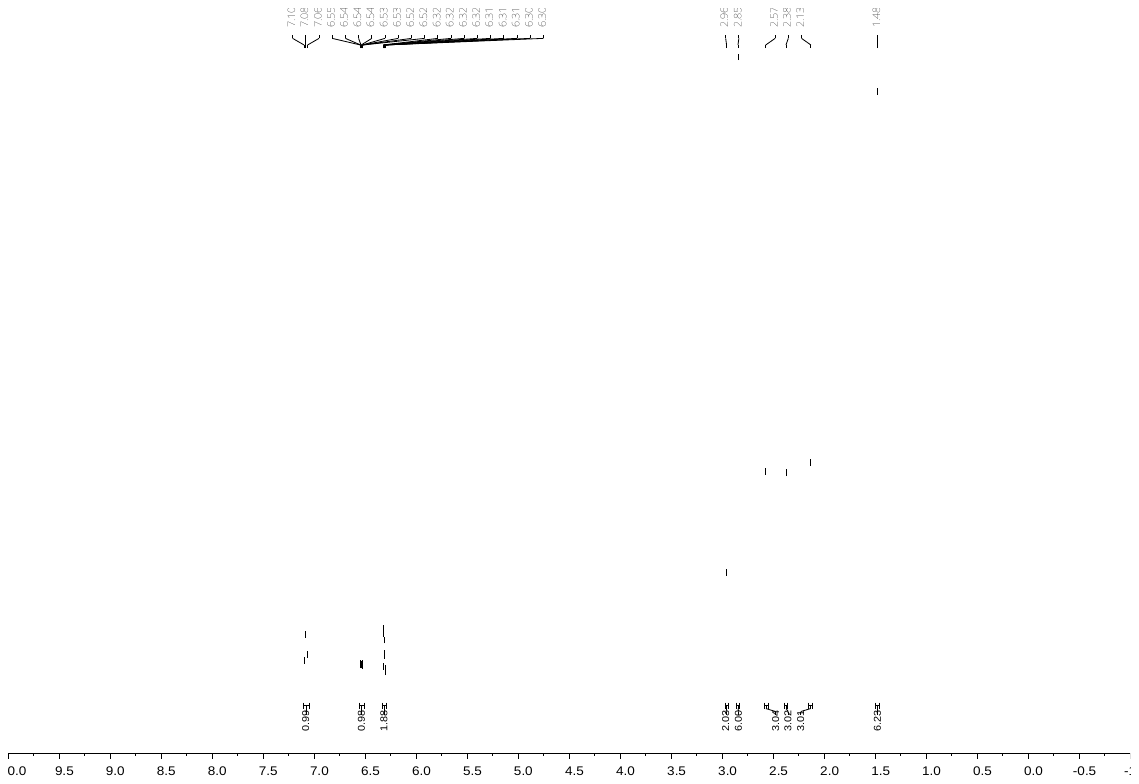

^13^C NMR

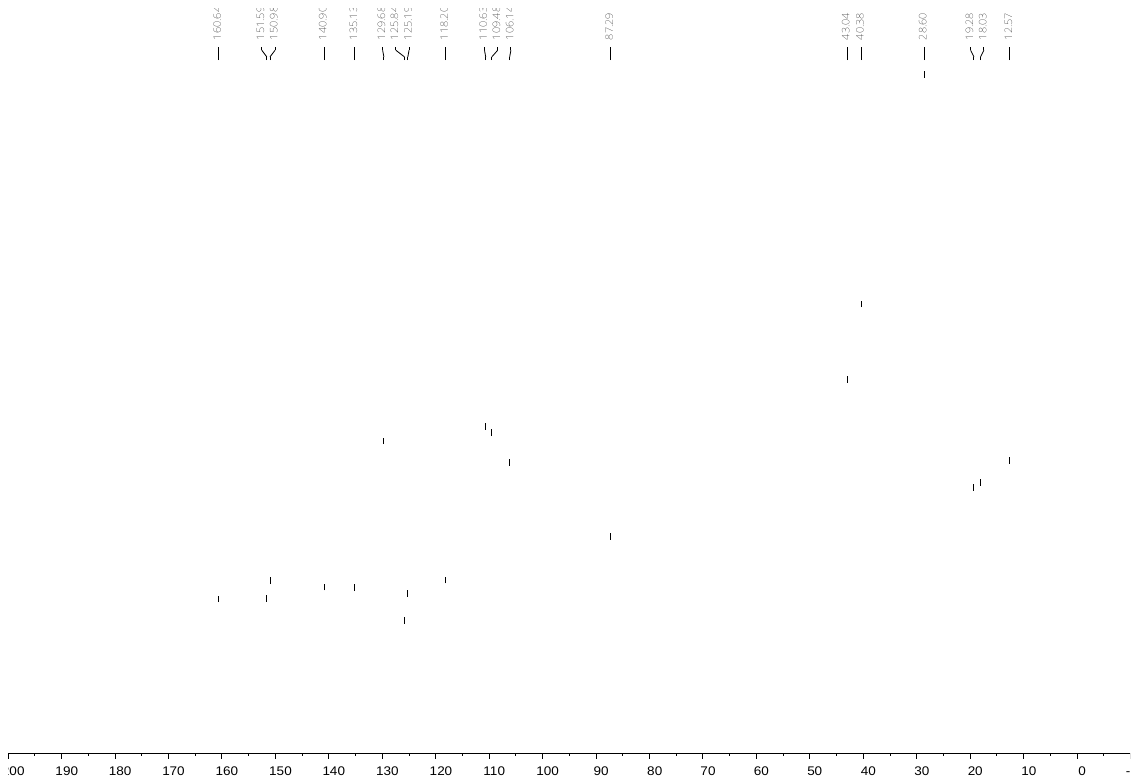

**3-Methylphenyl 2,2,5,7,8-pentamethylbenzochromane-6-sulfonate (19)**

^1^H NMR

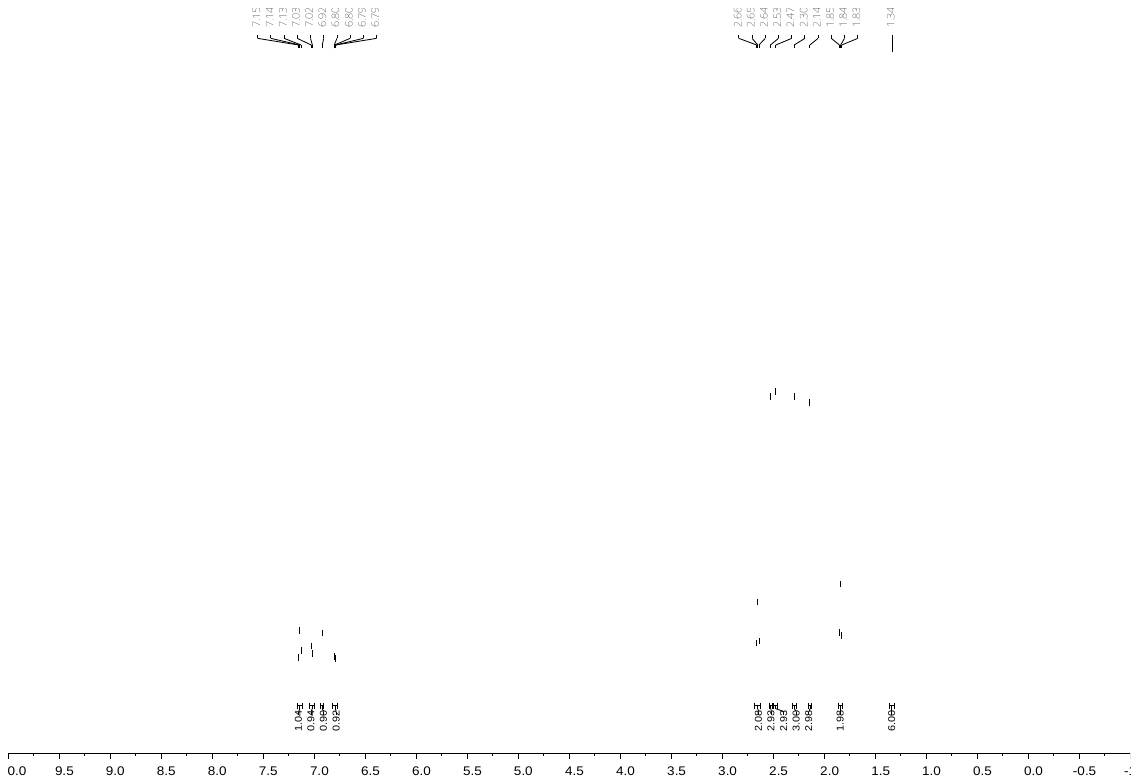

^13^C NMR

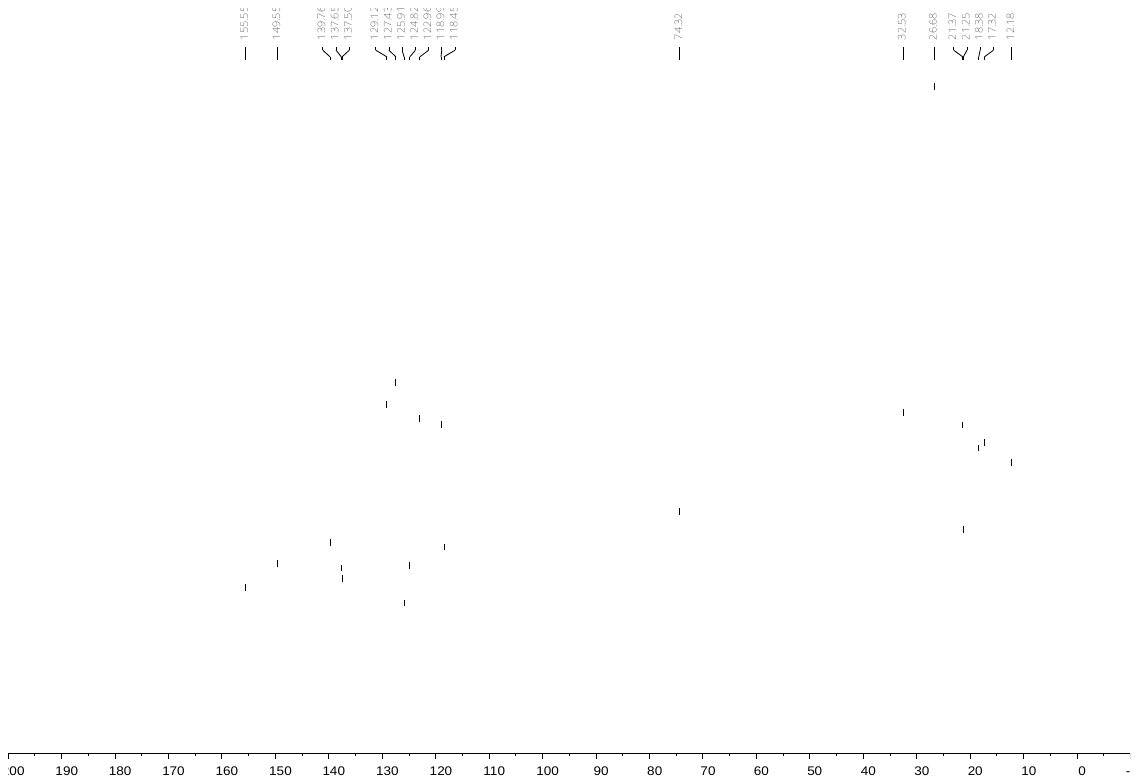

**Phenyl 2,2,4,6,7-pentamethyldihydrobenzofuran-5-sulfamide (20)**

^1^H NMR

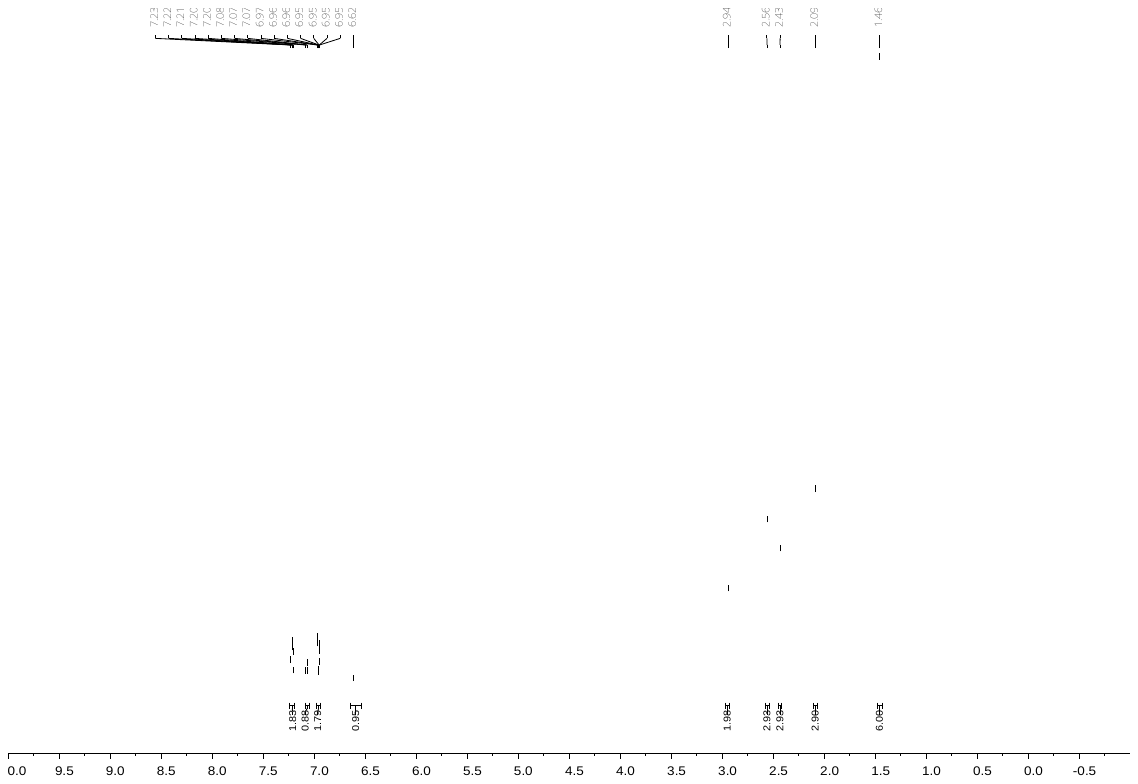

^13^C NMR

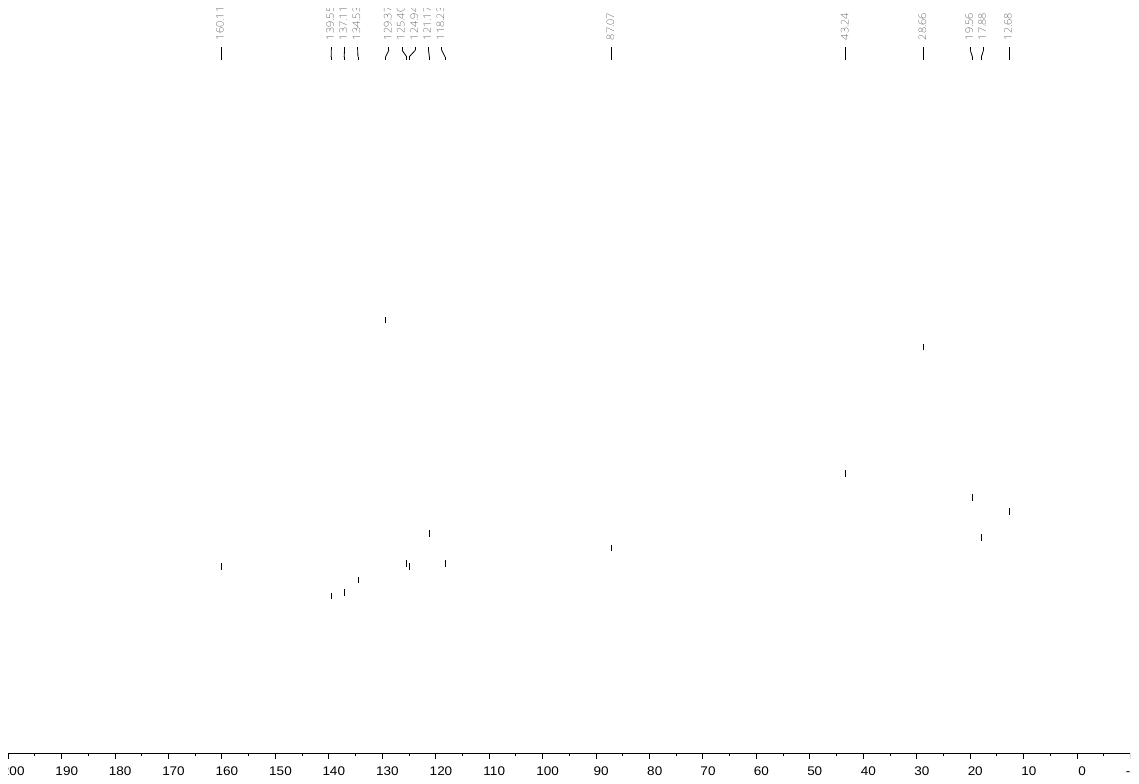

**3-Methylphenyl 2,2,4,6,7-pentamethyldihydrobenzofuran-5-sulfonamide (21)**

^1^H NMR

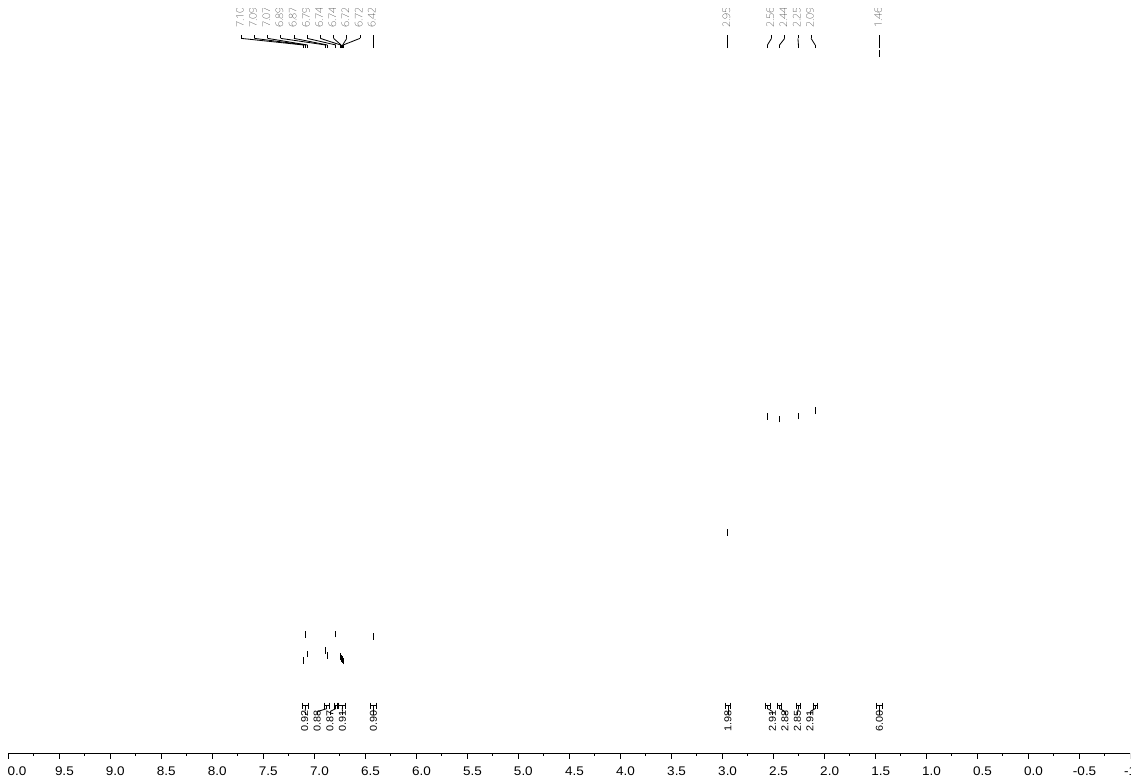

^13^C NMR

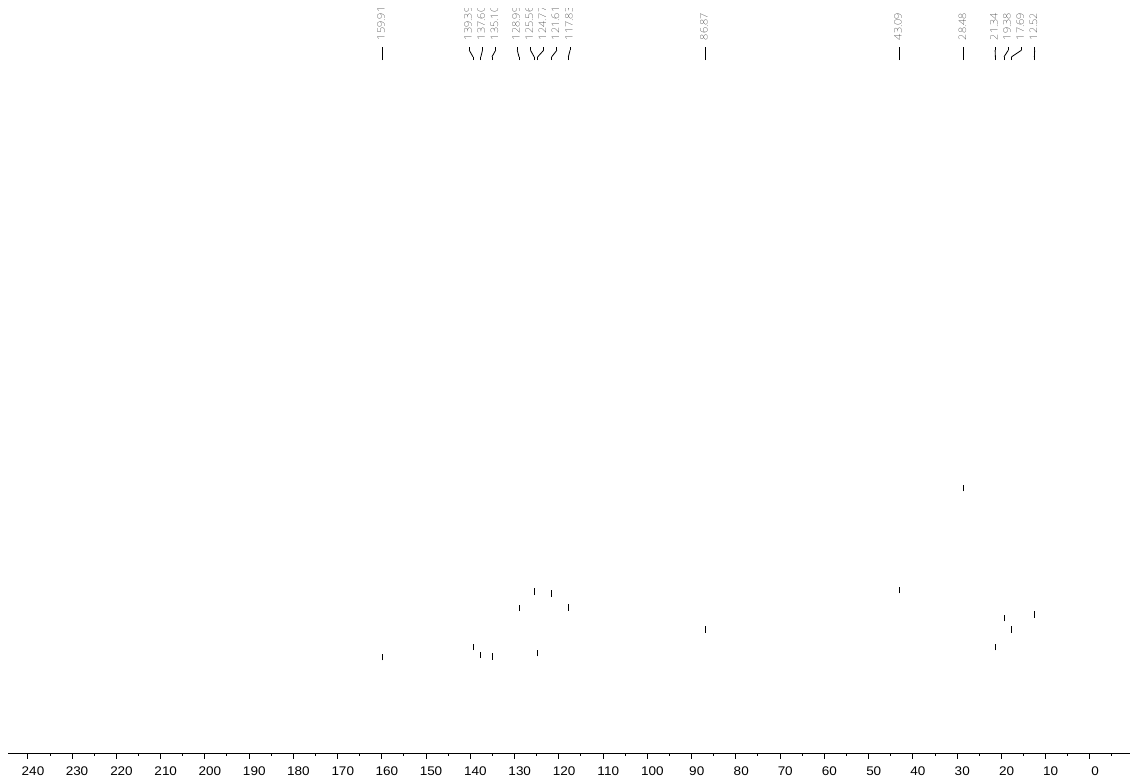

**Phenyl *N*-methyl-2,2,4,6,7-pentamethyldihydrobenzofuran-5-sulfonamide (22)**

^1^H NMR

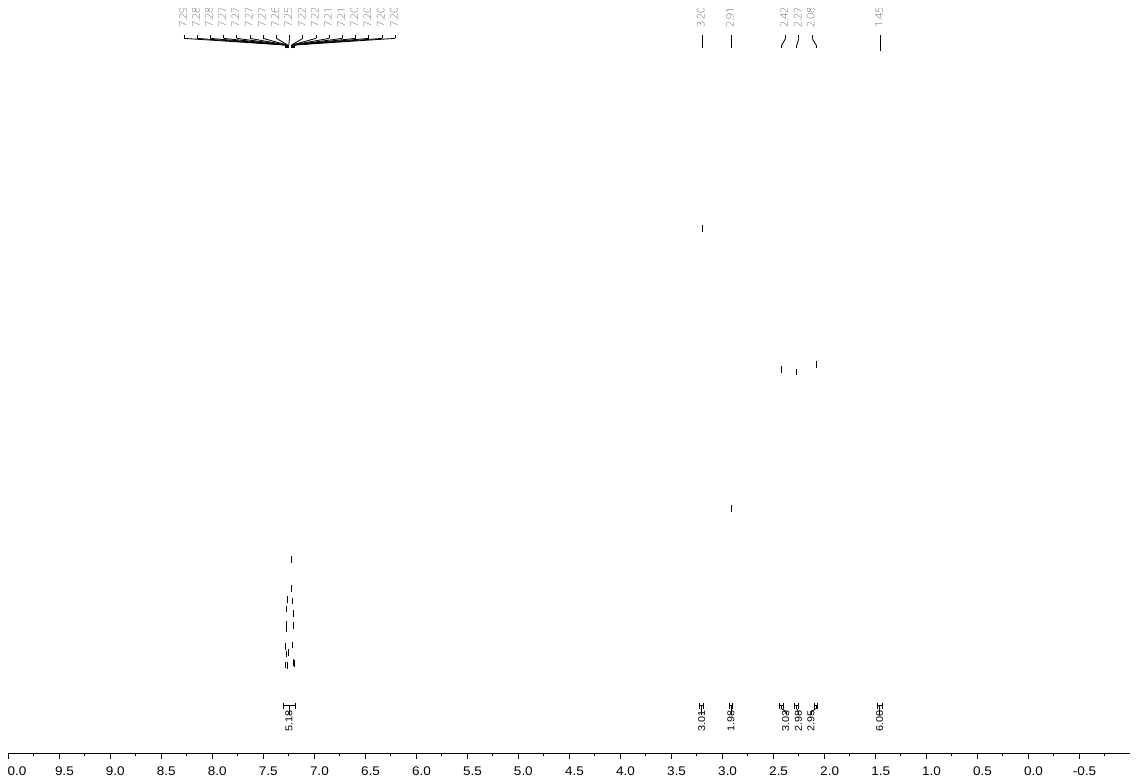

^13^C NMR

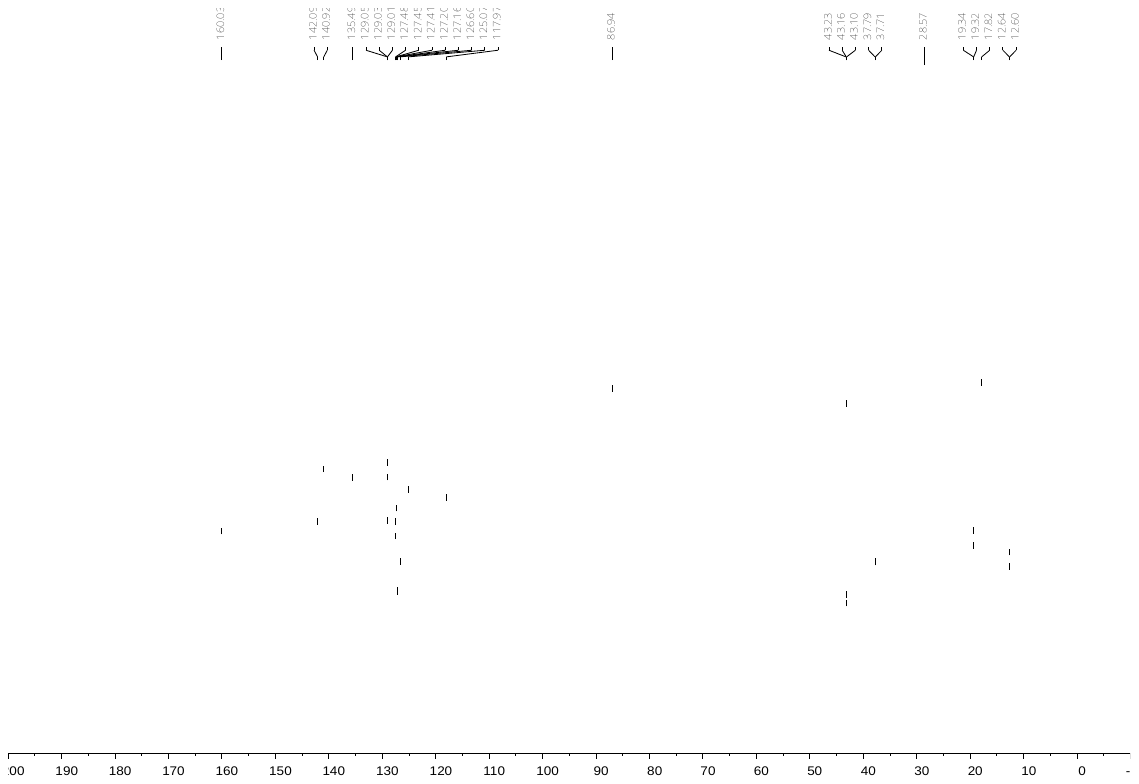

**3-Chlorophenyl 2,2,5,7,8-pentamethylbenzochromane-6-sulfonate (23)**

^1^H NMR

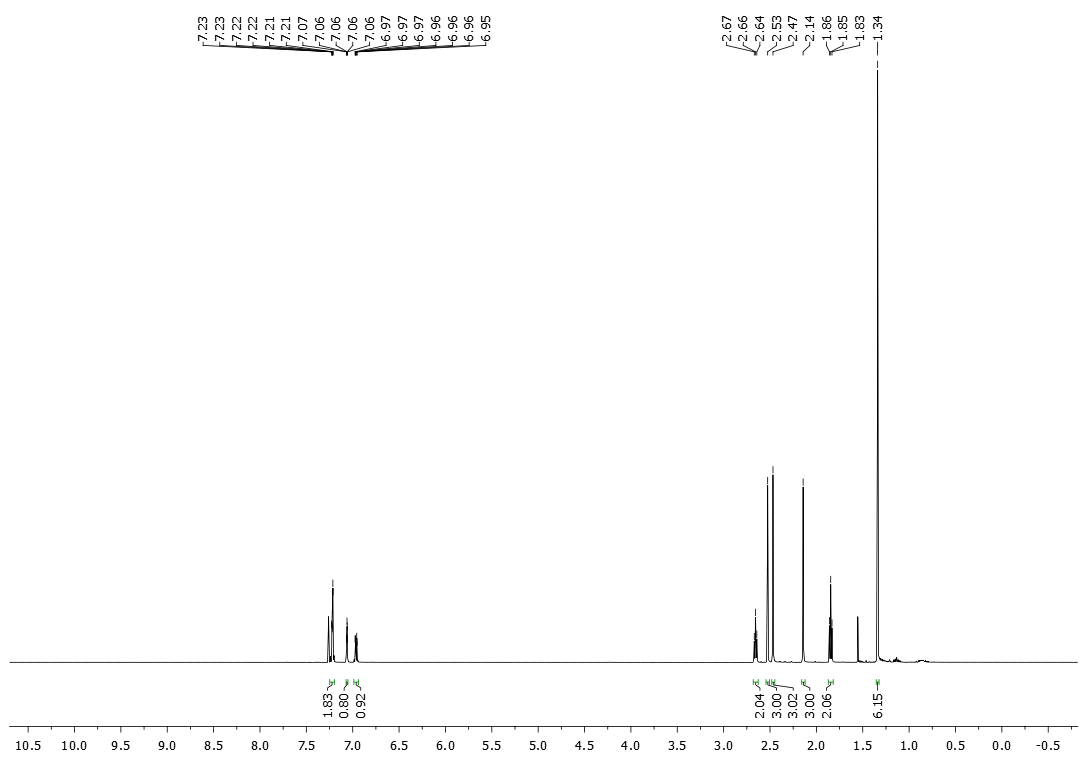

^13^C NMR

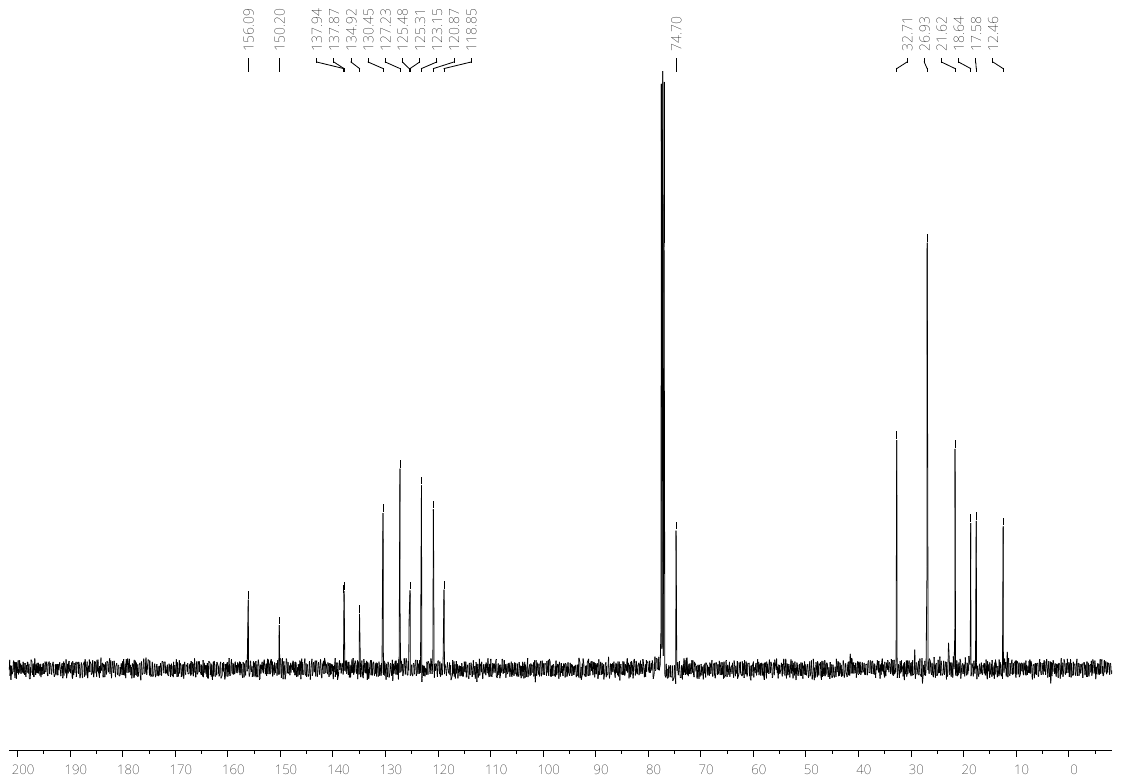

**2,2,5,7,8-Pentamethylchroman-6-yl 3-methylbenzenesulfonate (24)**

^1^H NMR

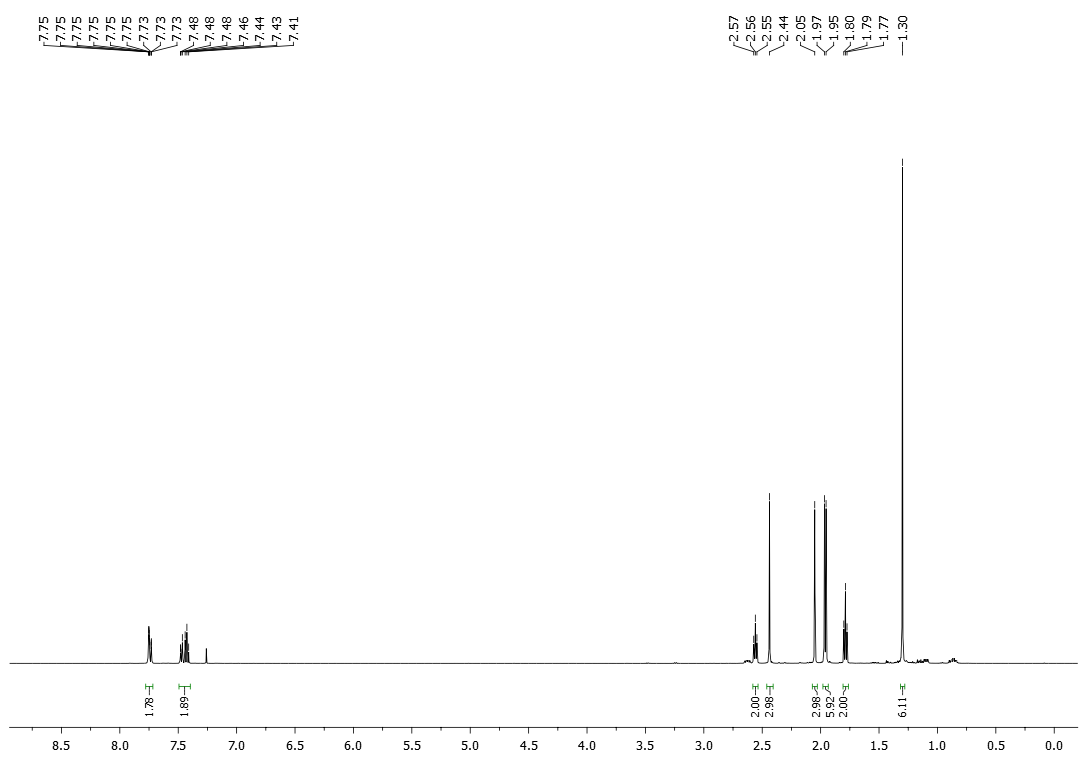

^13^C NMR

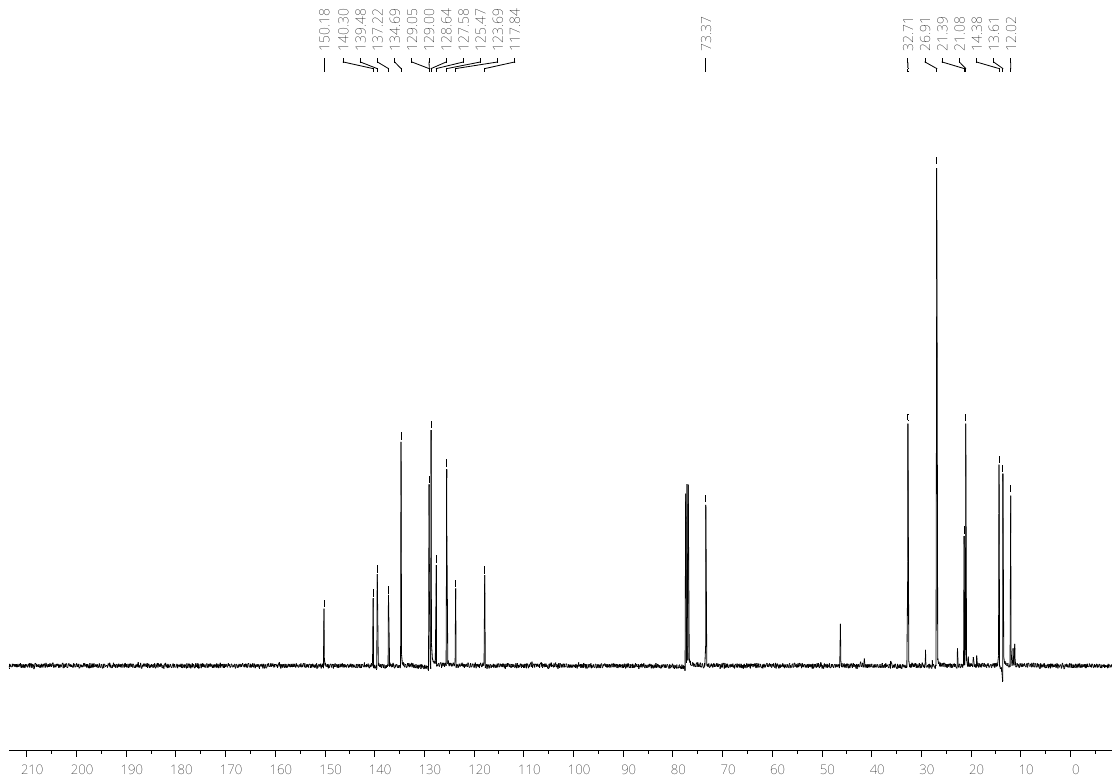
